# Correlations between human mating partners: a comprehensive meta-analysis of 22 traits and raw data analysis of 133 traits in the UK Biobank

**DOI:** 10.1101/2022.03.19.484997

**Authors:** Tanya B. Horwitz, Jared V. Balbona, Katie N. Paulich, Matthew C. Keller

## Abstract

Positive correlations between human mating partners are consistently observed across traits. Such correlations can increase phenotypic variation and, to the extent that they reflect genetic similarity in co-parents, can also increase prevalence for rare phenotypes and bias estimates in genetic designs. We conducted the largest set of meta-analyses on human partner correlations to date, incorporating 480 partner correlations across 22 traits. We also calculated 133 trait correlations between up to 79,074 male-female couples in the UK Biobank (UKB). Estimates of the mean meta-analyzed correlations ranged from *r*_meta_=.08 for extraversion to *r*_meta_*=*.58 for political values. UKB correlations ranged from *r*_UKB_=-.18 for chronotype to *r*_UKB_=.87 for birth year. Overall, attitudes, education, and substance use traits mostly showed the highest correlations, while psychological and biological traits generally yielded lower but still positive correlations. We observed high between-study heterogeneity for most meta-analyzed traits, likely because of both systematic differences between samples and true differences in partner correlations across populations.

## Main

Phenotypic resemblance between human mates, spouses, and co-parents (hereafter, “partners”) is a widely studied area of research with applications to an array of disciplines. Quantitative and behavioral geneticists are interested in the magnitudes and mechanisms underlying partner correlations because both can affect genetic and environmental parameters and estimates of interest, informing how model estimates should be interpreted^1–4^. Sociologists, anthropologists, economists, and other social scientists have studied partner correlations to gain insight into trends in societal values^5^ and the labor market^6,7^, socioeconomic inequalities^5–7^, and relationship satisfaction and divorce rates^8–13^. Research on shared disease/disorder risk factors and health-related behaviors could have public health implication with respect to screening and intervention practices^14–17^ . Meanwhile, research on the nature of partner correlations for behavioral similarities could potentially inform how partners can effectively make lifestyle changes together^18,19^, particularly if partner resemblance is due to mutual influences.

Contrary to the maxim “opposites attract,” nonzero phenotypic correlations between human^8,14,18, 20–36^ and nonhuman^37^ mates and partners are overwhelmingly in the positive direction, with only a handful of examples of significant negative partner correlations (“disassortative mating”) reported in the literature^9,15,21,23,35, 37–43^. Several potential mechanisms leading to partner resemblance in humans have been described. *Phenotypic homogamy* (also *phenotypic assortative mating*) occurs when partners match directly on the trait of interest^44^. While phenotypic homogamy is often conceptualized as partners actively preferring similarity, it can also arise from indirect selection, such as in parent-arranged marriage or when partners form within strata that are associated with trait values (e.g., partner correlations for educational attainment arising indirectly as a consequence of mate choice occurring within profession). *Social homogamy*, on the other hand, occurs when partners assort on non-heritable aspects of a trait^33,45^. This can occur through several different processes, such as when romantic partnerships form within strata that are heavily determined by non-heritable social factors (e.g., religion or social networks). *Genetic homogamy* describes the opposite phenomenon: when partners assort on only heritable aspects of a trait. Genetic homogamy may occur when there is phenotypic homogamy on a trait that is genetically but not environmentally correlated with the trait of interest^44,46^. Finally, *convergence* occurs when partners become more similar over time^20,23^, either due to direct (reciprocal or one-way) phenotypic influences or to the mutual influence of environmental factors shared between partners. Unlike social/phenotypic/genetic homogamy, convergence is not considered a type of assortative mating because it is not a consequence of initial matching (or assortment). Importantly, the processes that lead to partner correlations need not be mutually exclusive, as multiple mechanisms may work in concert to produce an observed correlation between partners for a given trait. For example, it is possible that—across all traits--a mixture of social homogamy and phenotypic homogamy is more common than “pure” social homogamy or “pure” phenotypic homogamy.

Phenotypic and genetic homogamy on heritable traits increases correlations between and within causal genetic loci, which in turn increases the genetic covariance between relatives as well as the trait’s genetic and phenotypic variation. Such an increase in variation could manifest as increased prevalence rates of dichotomous traits such as psychiatric disorders^33,47^, although this effect would only be pronounced for rare, highly heritable disorders for which there are strong positive correlations between partners^33^. Failing to account for phenotypic and genetic homogamy can also lead to biases in association statistics from genome-wide association studies^48^, heritability and genetic correlation estimates based on twin/family designs and single nucleotide polymorphisms^49,59^, and the strength of estimated causal associations in Mendelian randomization studies^50^. Moreover, while convergence and social homogamy have no genetic consequences, if parental traits influence offspring traits, these mechanisms can still increase the environmental and phenotypic variance of traits and the covariance between relatives. Thus, understanding the magnitude of partner correlations across a wide variety of traits is important for setting expectations of what analyses are at risk of bias due to the influences of homogamy and for guiding efforts to uncover the mechanisms that underlie these correlations.

While thousands of studies have reported estimates of partner correlations in humans, we are not aware of any meta-analysis that has done so across a diverse set of traits. In the current report, we use stringent methodology to meta-analyze and compare estimates of partner correlations for 22 frequently investigated traits in this research arena. Our analyses are based on 480 correlational estimates drawn from a total of 199 independent studies, making this the most comprehensive set of meta-analyses on human partner correlations to date. We also calculated 133 trait correlations between up to 79,074 male-female partner dyads from the UK Biobank (UKB) sample. These results can help shed light on contemporary human mating and relationship trends, refine the interpretation of heritability estimates, motivate investigation into the various causes of partner correlations across traits, and aid in the choice of design in genetic and non-genetic studies.

## Results

### Meta-analysis

Our random effects meta-analytic results included a total of 480 partner correlations across the 22 traits for which we were able to find results from at least three samples that satisfied our criteria. The total number of partners for each trait ranged from 2,527 (for generalized anxiety) to 2,727,151 (for diabetes). Supplementary Tables 1 and 2 show all studies that we included in our meta-analysis for continuous and dichotomous traits, respectively, as well as the effect sizes for each sample. For comparability across traits, we focus on Pearson’s, Spearman’s, and tetrachoric correlations for continuous, ordinal, and dichotomous traits, respectively.

Fig. 1 displays the estimates of the mean meta-analyzed correlations for all meta-analyzed traits and, where applicable, the correlations for comparable traits in the UK Biobank (UKB; see **Partner Correlations in the UK Biobank)**, along with the 95% confidence intervals associated with each trait. The estimates of the mean correlations for the meta-analyzed traits (*r*_meta_) were greater than zero at the nominal significance level (*p* < .05) for all but one trait (generalized anxiety) and significant at the Bonferroni-corrected (*p* < .05/22 = 0.00227) significance level for eighteen traits. *r*_meta_ was greater than .35 for nine attitudinal, academic, and substance-related traits (ranging from *r*_meta_ =. 38 for smoking initiation to *r*_meta_*=*.58 for political values; history of problem alcohol use showed the lowest correlation for substance-related traits at *r*_meta_*=*.28). Estimates of mean meta-analyzed correlations for anthropometric traits and (non-substance related) disorder traits were all low-to-moderate (.15 ≤ *r*_meta_ ≤ .24). The three lowest estimates we found were for the Big Five personality traits extraversion (*r*_meta_ = .08), neuroticism (*r*_meta_ = .11), and agreeableness (*r*_meta_ = .11). Point estimates for conscientiousness (*r*_meta_ *=* .16) and openness to experience (*r*_meta_ *=* .21) were slightly higher (see Table 1).

**Figure 1:**
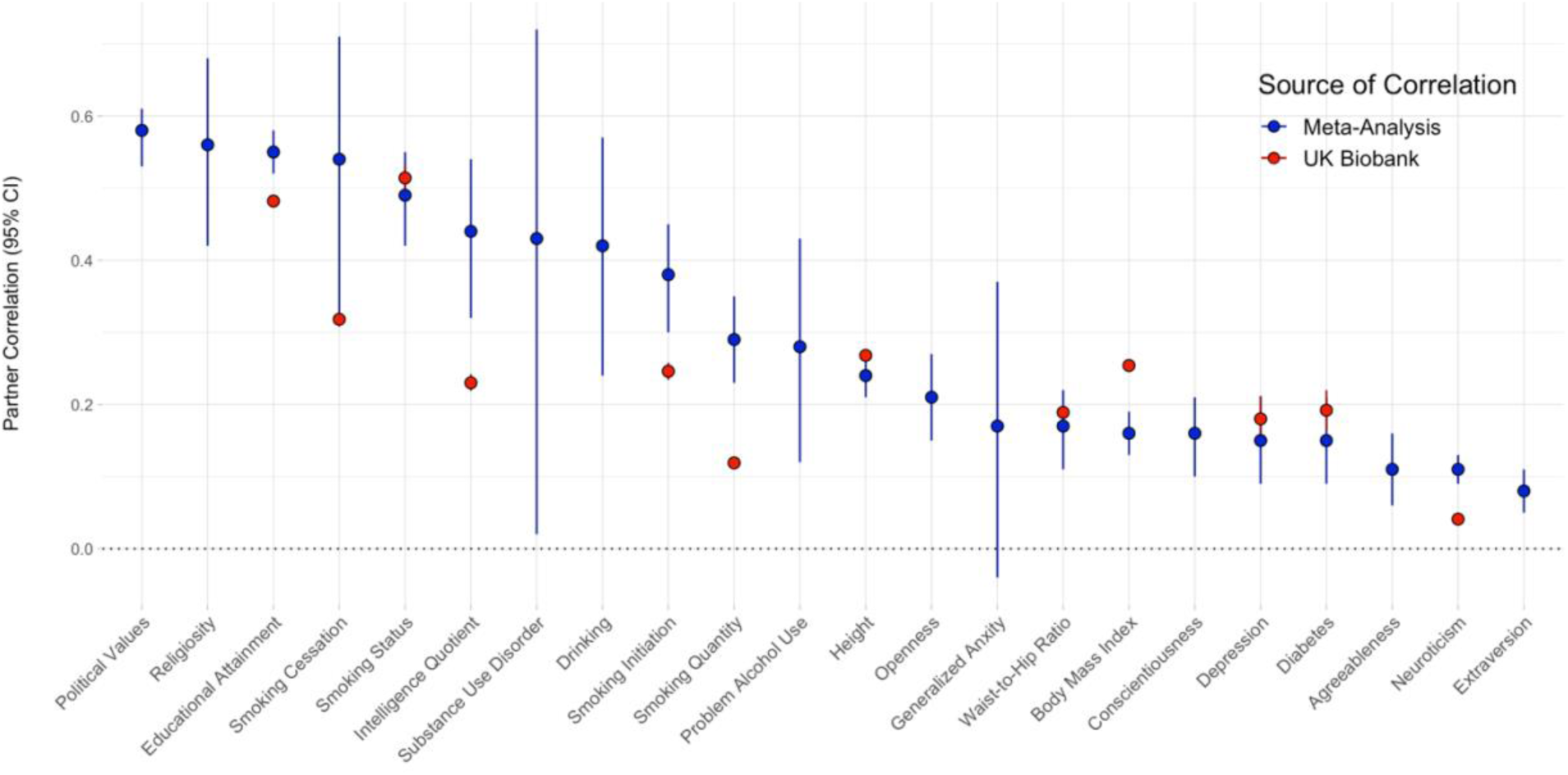
The point estimates of the mean meta-analyzed random effects partner correlations and UK Biobank partner correlations for comparable traits, along with their respective 95% confidence intervals. The dark blue points represent the random effects estimates of the mean meta-analyzed correlations for partners, while the red points on the same vertical axis represent the point estimates for a comparable trait in the UK Biobank, where applicable.

**Table 1:**
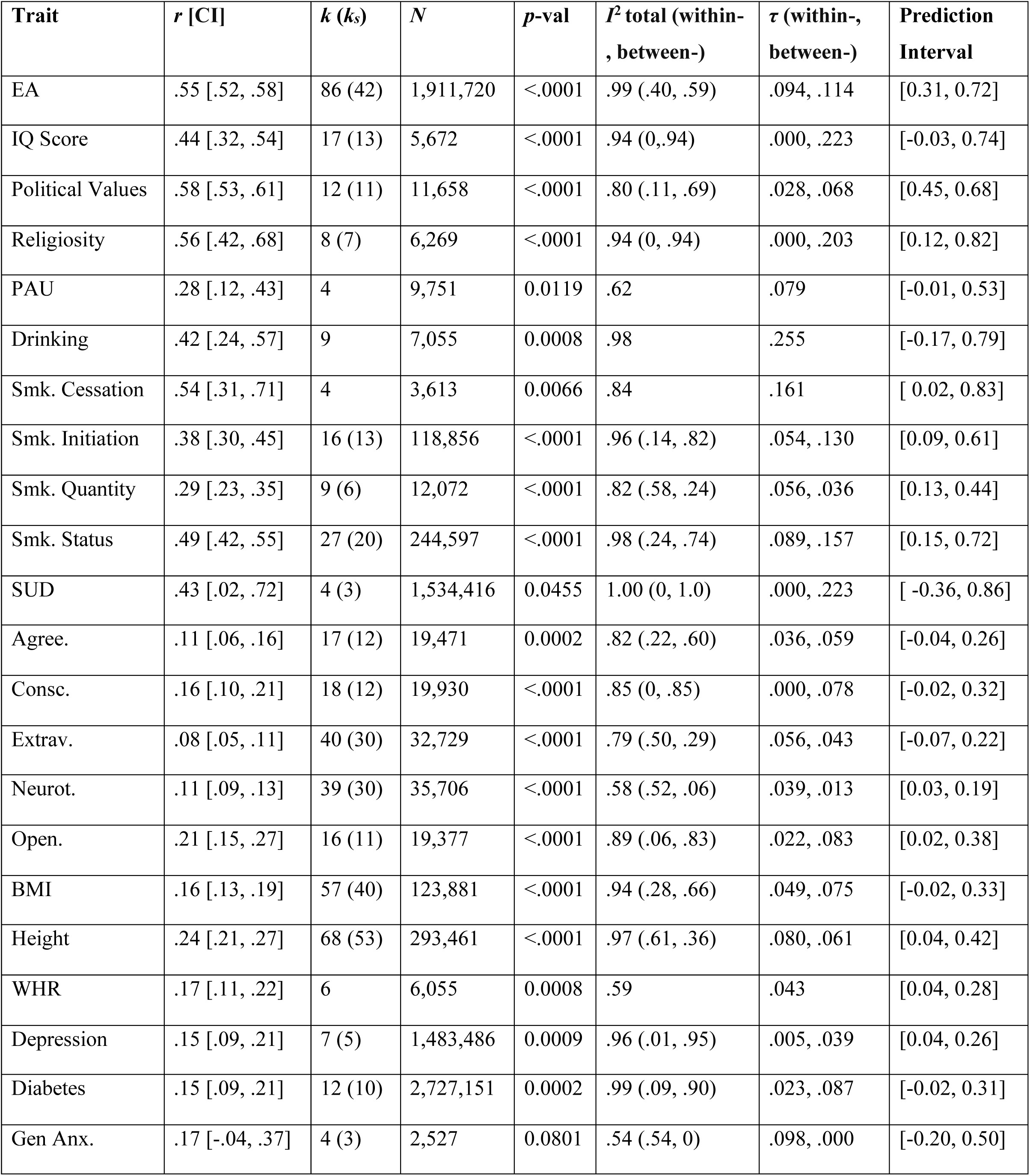
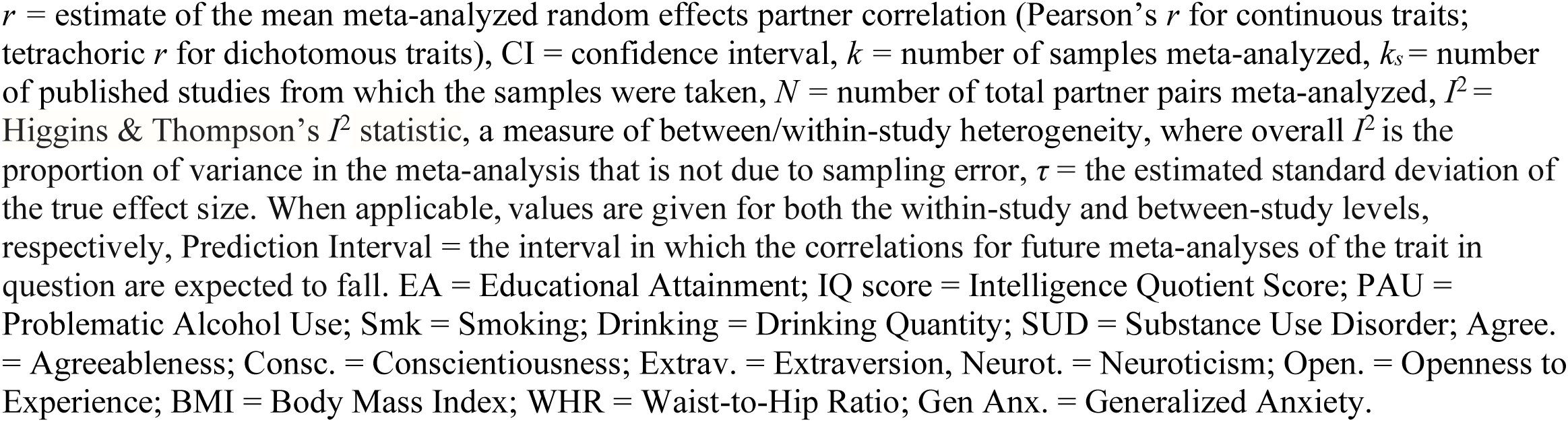
Results for the random effects meta-analyses of mating partner pairs across 22 traits.

| Trait | $r$ [CI] | $k$ ($k_s$) | $N$ | $p$-val | $I^2$ total (within-, between-) | $\tau$ (within-, between-) | Prediction Interval |
| --- | --- | --- | --- | --- | --- | --- | --- |
| EA | .55 [.52, .58] | 86 (42) | 1,911,720 | <.0001 | .99 (.40, .59) | .094, .114 | [0.31, 0.72] |
| IQ Score | .44 [.32, .54] | 17 (13) | 5,672 | <.0001 | .94 (0, .94) | .000, .223 | [-0.03, 0.74] |
| Political Values | .58 [.53, .61] | 12 (11) | 11,658 | <.0001 | .80 (.11, .69) | .028, .068 | [0.45, 0.68] |
| Religiosity | .56 [.42, .68] | 8 (7) | 6,269 | <.0001 | .94 (0, .94) | .000, .203 | [0.12, 0.82] |
| PAU | .28 [.12, .43] | 4 | 9,751 | 0.0119 | .62 | .079 | [-0.01, 0.53] |
| Drinking | .42 [.24, .57] | 9 | 7,055 | 0.0008 | .98 | .255 | [-0.17, 0.79] |
| Smk. Cessation | .54 [.31, .71] | 4 | 3,613 | 0.0066 | .84 | .161 | [ 0.02, 0.83] |
| Smk. Initiation | .38 [.30, .45] | 16 (13) | 118,856 | <.0001 | .96 (.14, .82) | .054, .130 | [0.09, 0.61] |
| Smk. Quantity | .29 [.23, .35] | 9 (6) | 12,072 | <.0001 | .82 (.58, .24) | .056, .036 | [0.13, 0.44] |
| Smk. Status | .49 [.42, .55] | 27 (20) | 244,597 | <.0001 | .98 (.24, .74) | .089, .157 | [0.15, 0.72] |
| SUD | .43 [.02, .72] | 4 (3) | 1,534,416 | 0.0455 | 1.00 (0, 1.0) | .000, .223 | [ -0.36, 0.86] |
| Agree. | .11 [.06, .16] | 17 (12) | 19,471 | 0.0002 | .82 (.22, .60) | .036, .059 | [-0.04, 0.26] |
| Consc. | .16 [.10, .21] | 18 (12) | 19,930 | <.0001 | .85 (0, .85) | .000, .078 | [-0.02, 0.32] |
| Extrav. | .08 [.05, .11] | 40 (30) | 32,729 | <.0001 | .79 (.50, .29) | .056, .043 | [-0.07, 0.22] |
| Neurot. | .11 [.09, .13] | 39 (30) | 35,706 | <.0001 | .58 (.52, .06) | .039, .013 | [0.03, 0.19] |
| Open. | .21 [.15, .27] | 16 (11) | 19,377 | <.0001 | .89 (.06, .83) | .022, .083 | [0.02, 0.38] |
| BMI | .16 [.13, .19] | 57 (40) | 123,881 | <.0001 | .94 (.28, .66) | .049, .075 | [-0.02, 0.33] |
| Height | .24 [.21, .27] | 68 (53) | 293,461 | <.0001 | .97 (.61, .36) | .080, .061 | [0.04, 0.42] |
| WHR | .17 [.11, .22] | 6 | 6,055 | 0.0008 | .59 | .043 | [0.04, 0.28] |
| Depression | .15 [.09, .21] | 7 (5) | 1,483,486 | 0.0009 | .96 (.01, .95) | .005, .039 | [0.04, 0.26] |
| Diabetes | .15 [.09, .21] | 12 (10) | 2,727,151 | 0.0002 | .99 (.09, .90) | .023, .087 | [-0.02, 0.31] |
| Gen Anx. | .17 [-.04, .37] | 4 (3) | 2,527 | 0.0801 | .54 (.54, 0) | .098, .000 | [-0.20, 0.50] |

$r$ = estimate of the mean meta-analyzed random effects partner correlation (Pearson's $r$ for continuous traits; tetrachoric $r$ for dichotomous traits), CI = confidence interval, $k$ = number of samples meta-analyzed, $k_s$ = number of published studies from which the samples were taken, $N$ = number of total partner pairs meta-analyzed, $I^2$ = Higgins & Thompson's $I^2$ statistic, a measure of between/within-study heterogeneity, where overall $I^2$ is the proportion of variance in the meta-analysis that is not due to sampling error, $\tau$ = the estimated standard deviation of the true effect size. When applicable, values are given for both the within-study and between-study levels, respectively, Prediction Interval = the interval in which the correlations for future meta-analyses of the trait in question are expected to fall. EA = Educational Attainment; IQ score = Intelligence Quotient Score; PAU = Problematic Alcohol Use; Smk = Smoking; Drinking = Drinking Quantity; SUD = Substance Use Disorder; Agree. = Agreeableness; Consc. = Conscientiousness; Extrav. = Extraversion, Neurot. = Neuroticism; Open. = Openness to Experience; BMI = Body Mass Index; WHR = Waist-to-Hip Ratio; Gen Anx. = Generalized Anxiety.

In addition to estimates of mean meta-analyzed correlations and sample sizes, Table 1 summarizes heterogeneity estimates for each meta-analysis as well as prediction intervals of future studies’ effect sizes. For traits meta-analyzed using random effects models with two levels, we calculated a single Higgins & Thompson’s *I*^2^, which estimates the proportion of variation resulting from heterogeneity in the effect sizes reported for that trait not attributable to sampling error. For traits meta-analyzed using models with three levels (those for which at least one publication reported two or more effect sizes), Table 1 shows two *I*^2^ values: one representing the proportion of variance resulting from heterogeneity within-study and one representing proportion of variance resulting from between-study heterogeneity, which sum to the total *I*^2^ (see **Meta-analytic Method**)^51^. The median within-study *I*^2^ for three-level analyses was .18 while the median between-study *I*^2^ was .715. Partitioning within-vs. between-study *I*^2^ estimates can yield unstable results, especially in meta-analyses that only involve a small number of effect sizes within a common study, and so should be interpreted with caution. Across our 22 meta-analyses, the median total Higgins & Thompson *I^2^* statistic was .915, reflecting a very high rate of heterogeneity in partner correlations across traits not due to sampling error.

In general, high *I*^2^ values may reflect not only high between-sample heterogeneity in estimated effect sizes but also low within-sample estimation error. Thus, the high *I*^2^ values we observed may be in part due to the high precision of estimates afforded by the large sample sizes of many of the studies included in our analysis. To this end, we also report an alternative metric of heterogeneity, *τ*, that estimates the standard deviation of the true effect size and is unaffected by the precision of individual estimates. The median overall *τ* was .090 and ranged from .039 (Depression) to .255 (Drinking). *τ* tended to be positively associated with *r*_meta_, with the average ratio of *τ* to *r*_meta_ (the coefficient of variation) being .373 (*SD* =.180) across traits.

Overall, our results show that partner correlations are characterized by substantial heterogeneity across samples. Given that traits measured objectively (e.g., height, body mass index, waist-to-hip ratio) were also associated with considerable heterogeneity, our observed *I*^2^ and *τ* estimates suggest that there are substantial differences in true partner correlations for a given trait across populations differentiated by location, time, and/or culture.

We produced funnel plots for each meta-analyzed trait to test for publication bias. Funnel plots demonstrating a large negative regression slope—particularly those with several studies clustering in the bottom-right but not the bottom-left–are indicative of publication bias. Overall, there were not obvious patterns of meaningful asymmetry across traits (Supplementary Fig. 3). The clearest trend across most of the funnel plots was the large number of points outside the expected triangular region, again reflecting the high heterogeneity in correlations observed across samples.

### Partner Correlations in the UK Biobank

We began by investigating partner correlations for 140 traits using up to 79,074 pairs of UK Biobank (UKB) participants inferred to be opposite-sex partners (see **Analysis of Partner Correlations in the UK Biobank**). While seven of these traits are not presented in Fig. 2 or Supplementary Fig. 1 because their analyses were under-powered, results for all 140 traits are presented in Supplementary Table 4, which includes summary statistics, cell frequencies and prevalences (for dichotomous traits), corresponding UKB field IDs, and trait descriptions. Of the 133 adequately-powered UKB traits presented in Fig. 2 and Supplementary Fig. 1--including 59 continuous, 24 ordinal, and 50 dichotomous traits (estimated using Pearson’s, Spearman’s, and tetrachoric correlations, respectively)--128 partner correlations (96%) were significant at the nominal (*p* < .05) threshold and 118 (89%) were significant at the Bonferroni-corrected level (*p* < .00036). The mean partner correlation across the 133 traits was .19 and ranged from *r*_UKB_*=* -.18 (chronotype) to .87 (year of birth).

**Figure 2:**
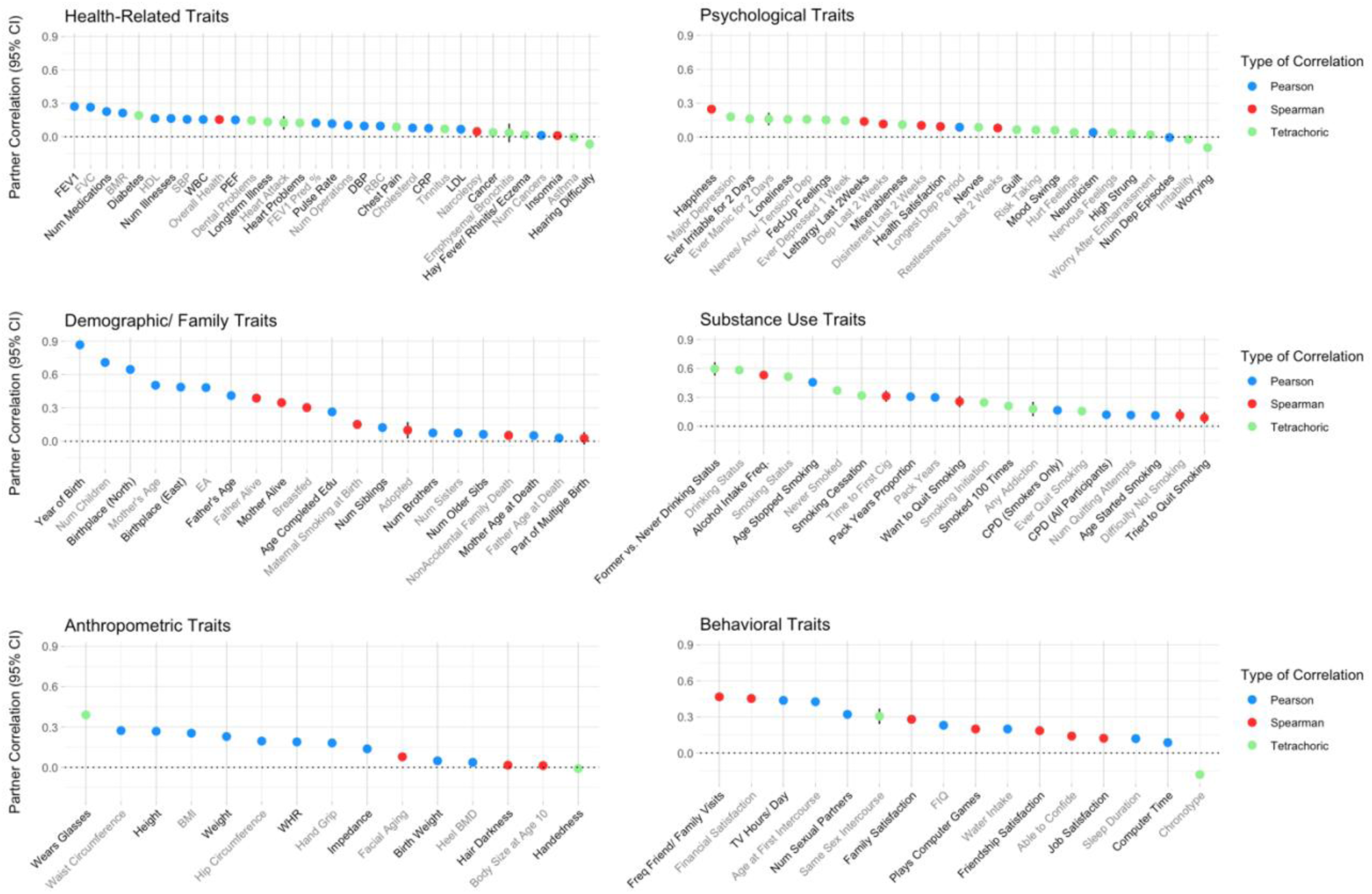
The UK Biobank partner correlation point estimates for 133 traits, along with their respective 95% confidence intervals, grouped by category of the trait. The point estimates on the y-axis represent the estimated partner correlation for the corresponding trait on the x-axis. Traits are grouped according to six categories: health-related, psychological, demographic/family, substance use, anthropometric, and behavioral. Points representing partner correlations for continuous traits (Pearson correlations) are blue; points representing partner correlations for ordinally-coded traits (Spearman correlations) are red; points representing partner correlations for dichotomously-coded traits (tetrachoric correlations) are light green. Num Dep Episodes = Number of Depressive Episodes; Heel BMD = Heel Bone Mineral Density (in the form of a t-score); LDL = Direct Low-density Lipoprotein Cholesterol, CRP = C-reactive Protein; RBC = Red Blood Cell (Erythrocyte) Count; DBP = Diastolic Blood Pressure; CPD (All Participants) = Cigarettes per Day (Includes Current, Former, and Never Smokers); FEV1 Pred % = Forced Expiratory Volume in 1-Second (FEV1), Predicted Percentage; PEF = Peak Expiratory Flow; WBC = White Blood Cell (Leukocyte) Count; SBP = Systolic Blood Pressure; HDL = High-density Lipoprotein Cholesterol; CPD (Smokers Only) = Cigarettes per Day (Restricted to Current or Former Smokers); WHR = Waist-to-hip Ratio; BMR = Basal Metabolic Rate; FIQ = Fluid Intelligence Quotient; BMI = Body Mass Index; FVC = Forced Vital Capacity; Time to First Cig = Time to First Cigarette; EA = Educational Attainment.

Twelve of the traits that we analyzed in the UKB were also meta-analyzed, allowing us to compare the meta-analytical estimates to the partner correlations found in the UKB (Fig. 1). Pearson and tetrachoric correlations for all but one (cigarettes per day) of these twelve overlapping UKB traits fell within the prediction intervals (distinct from the confidence interval) of corresponding traits from our meta-analyses, and the Spearman correlation for cigarettes per day in the UKB fell in the prediction interval for Smoking Quantity in the meta-analysis. While correlations for several UKB traits were significantly different than the estimates of the mean meta-analyzed correlations for the corresponding trait, there were notable overall trends: for example, estimates for educational attainment and current smoking status ranked amongst the highest correlations while estimates of partner correlations for neuroticism were uniformly low across the two analyses. Our UKB results, when compared to our meta-analytical results, recapitulate the primary findings from the meta-analyses: partner correlations tend to be positive but the true correlational values differ notably between populations.

Using weighted least squares regression 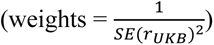, we regressed the UKB partner correlation point estimates on the category of trait (a factor with six levels: anthropometric, health-related, psychological, behavioral, demographic/family, and substance use) for the 133 adequately-powered UKB traits, controlling for how the correlations were estimated (Pearson’s, Spearman’s, or Tetrachoric). There were significant differences between mean correlations depending on the category of trait (*F*(5,125) = 28.57, *p* < 2e-16). In a follow-up analysis, we found that traits that were anthropometric (*r̄_UKB_* = .15, SE = .031), health-related (*r̄_UKB_* = .11, SE = .013), or psychological (*r̄_UKB_* = .09, SE = .014) yielded significantly lower partner correlations on average than traits related to behavior (*r̄_UKB_* = .24, SE = .042), demography/family (*r̄_UKB_* = .29, SE = .055), or substance use (*r̄_UKB_* = .29, SE = .036), (*t*(129) = 6.13, *p =* 1e-08). Controlling for category of trait, magnitude of partner correlations did not differ based on correlation type (*F*(2,125) = 1.79, *p* = .170).

Several of the highest positive zero-order UKB partner correlations for demographic variables are unsurprising given that individuals tend to marry and enter into long-term relationships with individuals who are of similar ages and from nearby geographic locations (e.g., *r*_UKB_ for north coordinate of birthplace *=* .65) and with whom they often share children (*r*_UKB_ for number of children *=* .71). More interestingly, partner correlations for several substance use traits were also very high. For adequately-powered substance use traits, we calculated the highest correlation for former vs. never drinking status (amongst only current non-drinkers, whether or not a person is a previous or never drinker; *r*_UKB_ *=* .60), followed by drinking status (whether or not a person currently drinks alcohol; *r*_UKB_ *=* .58); however, drinking status yielded a larger odds ratio (OR = 10.40) than former vs. never drinking status (OR = 6.32), as did smoking status (OR = 7.21). We evaluated smoking behavior with multiple measures and, of these, found the highest correlations for smoking status (whether or not a person currently smokes tobacco; *r*_UKB_ = .51) and the age that individuals stopped smoking (restricted to former regular smokers; *r*_UKB_ *=* .46) and the lowest correlation for presence and duration of quitting attempts (measured ordinally) in current regular smokers (*r*_UKB_ *=* .09). While we were underpowered in the UKB to examine concordance for addictions to specific narcotic or prescribed substances (e.g., alcohol, over-the-counter medication), addiction to ‘any substance or behavior’ showed a significant correlation (*r*_UKB_ *=* .18), suggesting a modest positive relationship between partners for general addiction.

Consistent with the results from our meta-analyses, anthropometric, health-related, and psychological traits generally showed low-to-moderate correlations, with some of these traits showing no significant correlation between partners. Correlations for height and waist-to-hip ratio (*r*_UKB_ *=* .27 and *r*_UKB_ *=* .19, respectively) were similar to *r*_meta_ for the corresponding traits in our meta-analysis, though the UKB estimate for body mass index (*r*_UKB_ *=* .23) was notably higher than *r*_meta_ for body mass index. Partner correlation for overall neuroticism score was low (*r*_UKB_ *=* .04) and partner correlations for the twelve dichotomous (yes/no) items comprising the neuroticism scale ranged from -.09 to .16 (see Supplementary Table 4). Finally, three dichotomous depression traits that we analyzed in the UKB all yielded modest correlations that were similar to one another: *r*_UKB_ = .15-.18.

Meanwhile, Fluid IQ score (classified as a behavioral trait) was correlated at *r*_UKB_ = .23 between partners in the UKB, and educational attainment (classified as a demographic/family trait and measured with respect to level of academic qualification/degree achieved) yielded a partner correlation of *r*_UKB_ = .48. Both of these correlations were lower than their respective comparable *r*_meta_ traits.

Additionally, we explored multiple traits that did not overlap with those we examined in our meta-analysis. The largest negative correlation across all UKB traits was for chronotype, which was dichotomized to indicate whether an individual is a “morning person” or an “evening person” (*r*_UKB_ = -.18). Furthermore, we found moderate correlations for traits related to sexuality (age of first intercourse: *r*_UKB_ = .43; number of lifetime sexual partners: *r*_UKB_ = .32; ever had same-sex intercourse: *r*_UKB_ = .31). Lifetime number of same-sex sexual partners yielded a non-significant Pearson’s correlation of .04 and Spearman correlation of .29; however, this trait was only measured in the 40 couples in which both members reported having had same-sex intercourse. Finally, general happiness (*r*_UKB_ = .25) and measures of satisfaction with various life aspects (financial situation: *r*_UKB_ = .45, family relationships: *r*_UKB_ = .28, friendships: *r*_UKB_ = .19, job *r*_UKB_ = .12, and health *r*_UKB_ = .09) showed small-to-moderate, positive partner correlations.

## Discussion

In this study, we collated and synthesized results from a large number of studies on same-trait correlations in opposite-sex/gender couples, spousal pairs, and co-parents (collectively, “partners”) in order to provide a better understanding of the degree to which partners correlate across traits and the degree of heterogeneity in those correlations across samples. We then analyzed partner correlation for 133 traits in putative partners in the UK Biobank (UKB). To our knowledge, this is the largest and most comprehensive set of analyses on human partner correlations to date.

We found widespread evidence of positive correlations across traits, with significant variation across traits with respect to degree of partner similarity. Across both the meta-analyses and UKB raw data analyses, correlations for educational attainment, substance use measures, attitudinal traits, and behavioral traits were often moderate and relatively higher, whereas correlations for anthropometric, psychological, personality, and health-related traits were mostly low or moderate. In our meta-analysis, we calculated the highest mean meta-analyzed partner correlations for political and religious values, educational attainment, IQ score, and some substance use traits. Though estimates for other traits were smaller, all but one was nominally significant (*p* < .05), with 18 of 22 traits reaching Bonferroni-level significance. Meanwhile, partner correlations for the vast majority of traits reached Bonferroni-corrected significance in the UKB. Overall, patterns of correlations in the UKB were similar though not identical to what we observed in our meta-analysis.

In addition to finding widespread positive associations in mating pairs across traits, our meta-analysis found evidence of substantial within-trait heterogeneity across different samples. Because of this large degree of heterogeneity and because we performed random (rather than fixed) effects meta-analyses, the results presented here should not be interpreted as estimates of a single true partner correlation for a given trait, but rather as estimates of the typical level of partner similarity across many possible levels that might be observed in samples drawn from different populations. Our results suggest that much of the observed between-study and between-sample heterogeneity is due to true differences that exist across populations (cultures, time periods, etc.) from which our samples were drawn. This is sensible given that resemblance between partners can result from personal preferences, social stratification, and/or couple dynamics that are unlikely to be consistent across different cultural contexts for a given trait. Although population size and/or mobility across populations may impact the pool of a person’s potential partners, there was insufficient diversity within the samples we included in most of our meta-analyses to test whether there were significant differences in partner correlations across specific geographical regions. Similarly, we were unable to formally assess changes in partner resemblance over time, as publication year served as an inaccurate measure of the year in which partners entered into a relationship, a variable which few studies reported. Thus, more stringently examining differences in partner concordance across cultures and time periods remains a direction for future studies.

In addition to the possible causes described above, the between-study heterogeneity in effect size estimates may also be partially due to differences in how constructs were assessed across studies and samples (e.g., differences in the measurement batteries used, participant interpretations of battery items, and clinical thresholds employed). Potentially consistent with this possibility, we observed that the prevalence rates of dichotomous traits were highly heterogeneous across supposedly non-ascertained samples, which may have contributed to the heterogeneity we observed in our correlation coefficients (two identical odds ratios translate into different tetrachoric correlations if the prevalence rates in the samples differ). Nevertheless, we observed high levels of heterogeneity in correlations for traits—such as height and body mass index—that are not dichotomous and are measured in standardized ways, suggesting that measurement invariance and ascertainment are unlikely to be a complete explanation.

There are several implications for the consistent evidence of partner similarity we documented across traits in our meta-analysis. First, partner similarity due to phenotypic or genetic homogamy within a population can bias estimates of genetic correlations^52^, latent variances from twin/family studies^25,53,54^, Mendelian randomization^50^, and SNP-heritability^49^. To the degree that observed levels of partner similarity are due to these mating mechanisms, our results suggest that, for many traits, some degree of bias due to assortative mating may exist in the aforementioned estimates and designs. Second, genetically informed designs that attempt to model assortative mating often must assume that degree of assortment has been stable across time and place; our observations of high heterogeneity across samples calls this assumption into question. Third, to the degree that partner similarity reflects genetic or phenotypic homogamy, partner similarity can increase the genetic variance and prevalence of disorders. Although the increase in prevalence for common disorders may not be large (e.g., ∼10%), homogamy for a rare, highly heritability trait could increase the trait’s prevalence more substantially^33^. Additionally, phenotypic homogamy, genetic homogamy, social homogamy, and convergence can each increase phenotypic variation for the trait in question in future generations. Finally, partner correlations have social and interpersonal implications that are relevant to outcomes of mating partners regardless of whether they procreate. For example, there is longitudinal evidence that smokers with a smoking partner are less likely to quit smoking^19^, while other findings suggest that being married reduces drinking quantity among men but increases drinking quantity among women, with divorce having the opposite effect^55^. Thus, knowledge of patterns in partner correlations offers an important understanding of the context shaping complex human traits and refines our interpretations of existing results.

While the current paper mostly targeted partners who were in a relationship at the time of study, previous research in dating and married couples concerning the association between partner similarity and likelihood of couples staying together has further social implications that extend beyond downstream genetic and environmental consequences in procreating pairs. Past research on this topic has found that partner similarity for variables such as academic performance and aspirations, involvement in the relationship^56^, political and religious beliefs^38^, mental distress, exercise, drinking, and smoking^57^ may be predictive of relationship longevity. Further investigation of these trends may be of interest to social scientists studying human relationships.

This study had limitations that we were not able to address. First, the vast majority of samples in our meta-analyses were drawn from Europe and the United States, with a smaller number coming from East Asia; meanwhile, samples residing in Sub-Saharan Africa, South Asia, Latin America and the Caribbean, Southwest Asia and Northern Africa, and Australia and Oceania (the last of which is notably less populous than the other regions) were each present in only a handful of studies. Additionally, participants in the UKB are disproportionately healthy, less likely to live in socioeconomically deprived areas, and less likely to smoke and drink on a daily basis; a large majority of those in the UKB are also of white European ancestry, which is reflective of the United Kingdom but not of the overall human population^58^. Thus, findings from both our meta-analysis and partners in the UKB are unlikely to generalize to all human populations and time periods. Relatedly, despite comprising a non-negligible share of couples and parents throughout time, partners and co-parents who did not mutually select each other— either because of cultural norms around marriage or because of coercion—are unlikely to be proportionally reflected in recent volunteer samples. Next, there was a broad range of total sample sizes used to calculate partner correlations across traits, particularly in the meta-analysis, suggesting some degree of imbalance in power across analyses. Finally, the results of the present study must be interpreted as cross-sectional estimates of partner resemblance and so are not directly informative with respect to selection mechanisms or longitudinal trends. Future studies will therefore be necessary to understand the specific processes underlying partner similarity (i.e., phenotypic/social/genetic homogamy and convergence), including how these processes have changed over time.

In sum, we have conducted the largest and most comprehensive set of analyses of human partner correlations to date, with estimates based on nearly a century of research and millions of partners. Across both our meta-analyses and our study of partners in the UKB, we generally found the highest estimates of similarity for traits related to substance use, educational attainment, and attitudes/behaviors and smaller correlations for personality, psychological, anthropometric, and disorder traits—though there was variability with respect to partner correlations within trait categories. We also observed high levels of heterogeneity in point estimates across studies for most traits investigated, suggesting that partner resemblance may differ across time and/or place. Research into the resemblance between partners and co-parents provides valuable information about the nature of relationships and cultural norms around human mating, as well as information pertinent to the etiological roots of disorders and to the interpretation of results from genetically informed designs.

## Methods

### Literature Search for the Meta-analysis

For the meta-analytic portion of this paper, we conducted a systematic review of English-language studies that examined concordance rates in partners for the same—or very similar— complex traits. All included studies were published in peer-reviewed journals on or before August 16, 2022, and we excluded studies from books. The analysis was not pre-registered and no protocol was prepared because it was not a hypothesis-driven study.

We selected an initial 26 traits on which to conduct our meta-analysis. While partner concordance has been analyzed for many traits, we focused on those most studied in the literature as well as some less commonly studied dichotomous traits. First, we searched for words pertaining to the traits of interest (with the exception of height; see below and **Inclusion and Exclusion Criteria**) in conjunction with thirteen terms pertaining to partner correlations and assortative mating (see Supplementary Table 6 for exact search terms used) in the search engines PubMed (using the RISmed package^59^ in R) and ScienceDirect. We excluded educational attainment (EA) and body mass index (BMI) from our PubMed search because we had already obtained large sample sizes using only ScienceDirect (1,911,720 partner pairs from 86 samples for EA and 123,881 partner pairs from 57 samples for BMI).

Three authors then worked independently to evaluate whether each paper met the relevant criteria for meta-analysis and then manually collected data from relevant publications. One author then double-checked that each report determined to be applicable was indeed eligible for inclusion; if it was, all relevant findings were recorded in Supplementary Table 1 or 2 and included for meta-analysis. Supplementary Figs. 2a-2v display forest plots for every trait; each includes every correlation—as presented in Supplementary Table 1 or 2—associated with each included sample, with publications ordered by year and color-coded by geographic region(s) of the sample. Citations that were collated for assessment but that did not meet inclusion criteria are shown in Supplementary Table 3 (with the exception of a small number of studies corresponding to excluded traits, which were included in Supplementary Table 2; see *Dichotomous traits*), along with the reasons for each exclusion.

For each of the traits that we meta-analyzed, Supplementary Fig. 4 presents the number of results for each set of search terms and the number of studies included and excluded for each trait, grouped by reasons for exclusion, in accordance with PRISMA guidelines^60,61^.

### Inclusion and Exclusion Criteria

We restricted our analysis to studies of co-parents, engaged pairs, married pairs, and/or cohabitating pairs, with some studies containing a small number of divorced couples. We refer to the couples or co-parenting dyads as “partners” and refer to the female vs. male designation of individuals as “sex/gender” unless referring to a specific sample that uses one of these terms. We excluded samples and studies with a high proportion of same-sex/gender partners, which we believe warrant separate analysis because same- and opposite-sex/gender partners have shown different patterns of similarity for some traits^62,63^, because there is less data on the former, and because downstream genetic implications are contingent on whether or not a pair produces offspring. For traits such as personality, we required that measures be self-reported, but we permitted case statuses of psychiatric traits reported by family members or partners due to the low numbers of applicable studies for such traits. Except for studies that ascertained partners for the trait of interest, we excluded studies in which partners were ascertained in a way that might have affected the magnitude of concordance for the trait of interest (e.g., concordance for depression in parents of depressed children). We limited our analysis to studies with sample sizes equal to or greater than 100 partner pairs. When sample sizes were reported along with percentages for each cell of a contingency table, we inferred the sample size of each cell by multiplying the percentages by *n*. When studies reported partner concordance at different waves over time in the same sample, we reported the results from the first wave. If the *n* for a sample was not provided or inferable or we discovered an inconsistency or other point of confusion in a study that was conducted in or after 2000, we contacted the corresponding author(s) for clarification. In total, we contacted authors from nine different studies and five responded. Although we initially gathered data on rare psychiatric disorders, we did not formally meta-analyze the tetrachoric correlations for these traits because too few studies met our inclusion criteria (see Supplementary Tables 2 and 3).

For consistency, when reported concordance rates were stratified by a variable (i.e., age, length of marriage, or zygosity of a twin sample), we included results for each stratum as a separate entry. We calculated effect sizes from the data reported in primary studies rather than from other meta-analyses or reviews. However, because partner correlations for height have already been meta-analyzed extensively by Stulp *et al.* (2017)^24^, we analyzed only samples includes in the Stulp *et al.* paper for height in place of conducting a literature review for this trait. We pooled results from Stulp *et al.* in the same way we analyzed other continuous traits, after eliminating studies included in Stulp *et al.*’s meta-analysis in accordance with our exclusion criteria.

When samples for a given trait in multiple studies overlapped or were likely to overlap based on information provided in the publication, we only used the study that had the largest sample size or that we otherwise determined to be more appropriate. We also excluded samples taken from the UK Biobank (UKB) because we calculated partner correlations in this sample ourselves as a separate analysis. Finally, we restricted our meta-analysis to traits for which there were at least three samples that met our criteria.

For continuous and ordinal traits, we only included Pearson/interclass correlations and Spearman rank correlations. When a study presented cross-partner contingency tables for each level of an ordinal trait but a concordance statistic was not reported, we directly calculated the Spearman rank correlation and its corresponding standard error. Because most studies reported them, we analyzed raw correlations when studies reported both the raw correlation and the partial correlation(s) controlling for covariates. However, we included some correlations that controlled for age when the raw correlation was not available because age was a common covariate; additionally, we made occasional exceptions for types of adjustments (i.e. whether the participant was a twin) that attempted to correct for heterogeneity induced by the sampling or measurement scheme used in the study (see Supplementary Tables 1 and 2 for covariates associated with each sample). In every other case, we excluded studies that did not report raw partner correlations.

### Dichotomous traits

Within the context of ascertained studies (i.e., those that used probands and controls rather than randomly sampling from a population), we excluded those involving probands taken from a clinical setting (i.e., a hospital or other treatment facility) when partners of probands were not required to be. Although such ascertainment can be dealt with if all the applicable populations’ (i.e., inpatient, outpatient, and those who have never received treatment) prevalence rates are known, it was typically difficult to know these rates. We eliminated any ascertained studies in which there was a >∼two-fold difference in male and female prevalence if there was not enough information to identify discordant partners based on sex/gender. Simulation results suggested that mixing male and female probands when their prevalence rates were more discrepant than this would lead to unacceptable levels of bias. Because of possible differences in the strength of calculated concordance based on an all-male proband sample versus that based on an all-female proband sample, we excluded studies that only included single-sex/gender probands. However, when data was available for *both* a female proband and a male proband (only a single study^64^), estimates based on each proband (female and male) were included as separate results.

We also restricted our meta-analysis to studies with expected contingency table cell frequencies of five or greater for all cells and observed cell frequencies greater than zero. Four of the traits in our supplementary tables posed a problem because they are rare (bipolar disorder and schizophrenia) or have not been studied in enough sufficiently large samples (panic disorder and phobia), resulting in contingency tables with zero frequency cells or with expected cell frequencies that were less than five in many instances. As a result, there were not enough studies meeting our inclusion criteria to justify formally meta-analyzing these four traits, though we show results from studies on these traits that mostly met our criteria in Supplementary Table 2.

We required that either odds ratios (OR), phi coefficients (Φ), contingency tables (from which an OR and tetrachoric correlation can be calculated), or—if the study was not ascertained (see **Effect Size Conversion**)–tetrachoric correlations, be reported for dichotomous traits since other effect sizes would not be appropriate for pooling with these statistics. No studies that met our inclusion criteria only reported risk ratios and so we do not further discuss them. All effect sizes for dichotomous traits that were not already in the form of a tetrachoric correlation were then converted to a tetrachoric correlation (see **Effect Size Conversion**).

### Effect Size Conversion

To make estimates across continuous and dichotomous traits more comparable, we converted all effect sizes for studies examining dichotomous traits to tetrachoric correlations (if the study did not report one). If the contingency table was unknown but the OR was reported, we first inferred the contingency table using an R function described in the supplementary methods of Peyrot *et al.* (2016)^33^ (which the authors provided to us) and then used the polychor() function from the “polycor” package^65^ in R^66^ to convert it to a tetrachoric correlation; if the values of the contingency table cells were unknown but the Φ was available, we simply used the phi2tetra() function from the “psych” package in R^66^ to convert it to a tetrachoric correlation. Both conversions required inputting male and female prevalences, each of which was based on published male and female prevalence rates for the trait or—if provided or inferable in the study and the sample was not ascertained—the prevalence rates indicated in the study (prevalences for both males and females are reported in Supplementary Table 2 for each sample in which a dichotomous trait was examined). When a contingency table was provided, we simply calculated the associated OR manually and, if the study was not ascertained, we calculated the tetrachoric correlation based on this contingency table using polychor().

For studies in which probands were ascertained, it was necessary to use the aforementioned R function from Peyrot *et al.* (2016)^33^ to produce an expected population (non-ascertained) contingency table based on the OR and the *population* prevalence prior to converting to a tetrachoric correlation. This correction is necessary because the case-to-control ratio in an ascertained sample is usually higher than in the general population, and so simply evaluating the correlation based on the uncorrected contingency table, which is based around the *sample* prevalence, would lead to a (usually upwardly) biased estimate.

### Meta-analytic Method

We conducted two- or three-level (see below) random effects meta-analyses based on Fisher Z-transformed correlations and their respective variances; we reversed this transformation when reporting final results (e.g., *r*_meta_) for ease of interpretation. We used two-level meta-analysis when none of the publications for a given trait reported multiple effect sizes and used three-level meta-analysis when at least one publication for a given trait reported multiple effect sizes. Two-level random effects meta-analyses allow for the estimation of heterogeneity due to sampling variance versus that due to true effect size differences between samples. Three-level meta-analyses further break down the true effect size differences to capture heterogeneity resulting from variation between studies versus heterogeneity resulting from variation within studies^51^. Partner correlations estimated in the same study could be more similar due to, for example, measures being taken at the same time or in the same culture or due to the use of common instruments, even if the samples in a study were collected independently.

We utilized restricted maximum likelihood (REML) for both two-level and three-level meta-analyses. For the two-level meta-analyses, we used the rma.uni() function in the “metafor” package^67^ in R to specify two levels (i.e., standard random effects modeling). For three-level analyses, we used the rma.mv() function, within which we set individual effect sizes (the Z-transformed partner correlations in different samples) to be nested within individual studies for the “random” argument. For both two-level and three-level meta-analyses, we set the test of the coefficient to follow a t-distribution and report two-tailed *p*-values.

For continuous traits, we specified the effect sizes as Fisher Z-transformed Pearson or Spearman correlations and specified their variances as 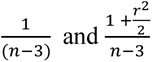, respectively, in accordance with Bonett & Wright (2000)^68^. For dichotomous traits, we converted the ascertained contingency table to a tetrachoric correlation corrected for ascertainment, as described above, and used the Fisher Z-transformed tetrachoric correlations in the meta-analyses. We used polychor() to calculate the untransformed standard error (*se_ut_*) for non-ascertained studies examining dichotomous traits. For ascertained studies examining dichotomous traits, we estimated *se_ut_* by creating 1,000 bootstrapped contingency tables, each of size *n* (the number of partner pairs) and sampling from the study’s (raw, ascertained) contingency table, with replacement, and then estimating the standard deviation of the 1,000 bootstrapped tetrachoric correlations arising from the contingency tables. We then transformed all *se_ut_*’s for tetrachoric correlations using the delta method^69^, wherein the standard error of the Z-transformed tetrachoric correlation is equivalent to 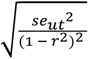, and used the square of these transformed standard errors for the variance argument in the rma.uni/rma.mv function. For reporting purposes, all meta-analyzed point estimates and their associated confidence intervals were back-transformed to their original format (Pearson’s correlation or tetrachoric correlation and their respective confidence intervals).

We estimated two sets of heterogeneity parameters, *τ* and *I*^2^, at the between-study level and—when applicable—within-study level. *τ* estimates the standard deviation of the true effect size at a given level. The total Higgins & Thompson’s *I*^2^ represents the percentage/proportion of variance not due to sampling error and is the sum of *I*^2^ at the between-study level and *I*^2^ at the within-study level. The heterogeneity statistics are detailed in Table 1.

Because studies on partner correlations ostensibly focus more on effect size than hypothesis testing, we expected that publication bias was unlikely to be a major factor for the study results we meta-analyzed. Nevertheless, we created funnel plots (Supplementary Fig. 3)--which plot study effect size (Fisher Z-transformed correlations here) on the x-axis against their standard errors on the y-axis--to visually inspect whether there was evidence for asymmetry, a potential indicator of publication bias.

### Analysis of Partner Correlations in the UK Biobank

In addition to our main meta-analysis, we analyzed correlations between inferred male-female partners in the UK Biobank (UKB) for an initial 140 ordinal, dichotomous, and continuous traits. We detected putative partners using a similar approach to past studies of partner correlations/assortative mating in the UKB^47,52^. Beginning with 502,414 individuals, we first restricted our sample to individuals who reported living with a “Husband, wife, or partner” and who did not report living with an unrelated roommate (based on data-field 6141), leaving 359,189 individuals. Using a co-location variable that was previously provided by the UKB, we subsequently limited our sample to pairs of individuals who were living at the same address at the time of recruitment, which narrowed the pool down to 200,707 participants. Afterward, using a series of genetic relationship matrices (GRMs) to calculate within genetic ancestry groups, we removed pairs of related individuals (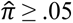), leaving 175,250 individuals; because GRMs were calculated within-ancestry, cross-ancestry pairs were removed at this stage. We then kept only pairs who were concordant for the number of people in their household, whether they own or rent their property, and their Townsend Deprivation Index (based on data-fields 709, 680, and 189, respectively), leaving 160,738 participants. Same-sex partners (data-field 31) were then removed, leaving 159,998 participants. Because most discrepancies in self-reported household income (data-field 738) were due either to the pairs choosing two adjacent options (e.g., one individual choosing “18,000 to 30,999” and the other “31,000 to 51,999”) or to non-responses, we removed only pairs whose reported household incomes differed by more than one category, leaving 158,350 individuals. Finally, pairs with age differences greater than 20 years were removed to minimize the possibility of including stepchild/stepparent pairs, leaving a final sample of 158,148 individuals comprising 79,074 putative mating pairs.

Notably, this subsample contains substantially more partners than certain previous studies using inferred partners in the UKB; for example, despite using similar procedures and criteria to detect partners, Yengo *et al*. (2018)^47^ detected only 18,984 pairs within the UKB. The discrepancy between these two sample sizes can be attributed to several causes, including our sample containing partners of non-European descent, us not requiring perfect matches on household income, and our use of the co-location variable. Additionally, Yengo *et al.* required that partners be concordant for the number of smokers in their household—a question only asked of non-smokers in the UKB, thus eliminating all pairs that contain one or more current smoker. Nonetheless, as shown in Supplementary Table 5, the magnitude of correlations within our subsample do not differ substantially from those of Yengo *et al.*’s subsample for the majority of traits analyzed in both studies, despite their study’s more conservative partner detection criteria that inadvertently ascertained for non-smoking concordance.

In some cases, we slightly altered the nature of traits by combining information from multiple traits or changing trait coding schemes (see Supplementary Table 4 for traits that were altered and the UKB descriptions of each trait we included). In line with our meta-analytic methods, we used responses from the initial wave when responses were collected at multiple times for the same trait. For each continuous trait, we eliminated partners containing at least one member with an absolute z-score greater than four to reduce the number of invalid responses and then calculated both a Pearson correlation and a Spearman rank correlation to provide two different estimates in the event that unusually-distributed data biased correlations. For zero-inflated data, both Pearson and Spearman rank correlations provide a slight underestimate of the true underlying correlation of two continuous measures where values below some threshold provide an observation of zero, but Spearman correlations tend to be less biased^70^. For dichotomous traits, we calculated tetrachoric correlations and odds ratios, and for ordinal traits, we calculated only Spearman rank correlations. In addition to the zero-order correlations for UKB partners, we also present partial correlations in Supplementary Table 4 that control for year of birth, whether an individual was born in the British Isles, and each individual’s first ten ancestral principal components. To avoid the re-introduction of covariate effects^71^, we ranked our ordinal data prior to residualizing. Out of the initial 140 traits, six dichotomous traits did not have expected cell frequencies of five or greater, and one continuous trait was under-powered. These traits were therefore included in Supplementary Table 4 but not in Fig. 2 or Supplementary Fig. 1, where we visualize the point estimates and confidence intervals for the remaining 133 traits.

## Data availability

Studies included in the meta-analysis are listed in Supplementary Tables 1 and 2, and studies excluded from the meta-analysis are listed in Supplementary Table 3. Raw data from the UK Biobank is not publicly available, but summary statistics for most traits are available on the UK Biobank website: https://biobank.ndph.ox.ac.uk/showcase/search.cgi.

## Code availability

The code for the analyses is available from the authors upon request.

## Author contributions statement

TBH contributed to study design, statistical analyses, manuscript writing/editing, assessment of meta-analyzed studies, and creation of figures and tables; JVB contributed to study design, statistical analyses, manuscript writing/editing, assessment of meta-analyzed studies, and creation of figures and tables; KNP contributed to manuscript editing, assessment of meta-analyzed studies, and creation of tables; MCK contributed to study design, statistical analyses, manuscript writing/editing, assessment of meta-analyzed studies, and creation of tables.

## Additional information

The authors declare no competing interests. MCK was supported by grants NIMH MH130448 and MH100141. This research has been conducted using the UK Biobank Resource under Application Number 16651.

## Notes

### Competing Interest Statement

The authors have declared no competing interest.

### Summary of Updates

We are submitting a revision of this manuscript because we have made substantial changes to the original preprint: namely, we added calculated partner correlations for putative partners in the UK Biobank, we re-conducted our literature review using more stringent criteria for the meta-analytic portion of the paper, and we applied multi-level modeling in our meta-analysis where applicable; we have also made other minor alterations to the paper and changed some language and terminology; finally, two additional authors have been added to the paper, and we have revised the main figures and supplementary material.

## References

1. Fisher, R. A. The Correlation between Relatives on the Supposition of Mendelian Inheritance. Earth Environ. Sci. Trans. R. Soc. Edinb. 52, 399–433 (1919).

2. Kong, A. et al. The nature of nurture: Effects of parental genotypes. Science 359, 424–428 (2018).

3. Eaves, L. The use of twins in the analysis of assortative mating. Heredity 43, 399–409 (1979).

4. Border, R. et al. Assortative mating biases marker-based heritability estimators. Nat. Commun. 13, 1–10 (2022).

5. Lavryashina, M. B. et al. Interethnic Dynamics among the Indigenous Peoples of Southern Siberia (Demographic Aspect)*. Archaeol. Ethnol. Anthropol. Eurasia 41, 131–142 (2013).

6. Blundell, R., Joyce, R., Norris Keiller, A. & Ziliak, J. P. Income inequality and the labour market in Britain and the US. Honor Sir Tony Atkinson 1944-2017 162, 48–62 (2018).

7. Schwartz, C. R. Trends and Variation in Assortative Mating: Causes and Consequences. Annu. Rev. Sociol. 39, 451–470 (2013).

8. Luo, S. Assortative mating and couple similarity: patterns, mechanisms, and consequences. Soc. Personal. Psychol. Compass 11, e12337 (2017).

9. Watson, D. et al. Match makers and deal breakers: analyses of assortative mating in newlywed couples. J. Pers. 72, 1029–1068 (2004).

10. Vandermeer, M. R. J., Kotelnikova, Y., Simms, L. J. & Hayden, E. P. Spousal Agreement on Partner Personality Ratings is Moderated by Relationship Satisfaction. J. Res. Personal. 76, 22–31 (2018).

11. Glicksohn, J. & Golan, H. Personality, cognitive style and assortative mating. Personal. Individ. Differ. 30, 1199–1209 (2001).

12. Dufouil, C. & Alpérovitch, A. Couple similarities for cognitive functions and psychological health. J. Clin. Epidemiol. 53, 589–593 (2000).

13. Feng, D. & Baker, L. Spouse similarity in attitudes, personality, and psychological well-being. Behav. Genet. 24, 357–364 (1994).

14. Jurj, A. L. et al. Spousal correlations for lifestyle factors and selected diseases in Chinese couples. Ann. Epidemiol. 16, 285–291 (2006).

15. Stimpson, J. P., Masel, M. C., Rudkin, L. & Peek, M. K. Shared health behaviors among older Mexican American spouses. Am. J. Health Behav. 30, 495–502 (2006).

16. Di Castelnuovo, A. et al. Cardiovascular risk factors and global risk of fatal cardiovascular disease are positively correlated between partners of 802 married couples from different European countries. Thromb. Haemost. 98, 648–655 (2007).

17. Knuiman, M. W., Divitini, M. L., Bartholomew, H. C. & Welborn, T. A. Spouse correlations in cardiovascular risk factors and the effect of marriage duration. Am. J. Epidemiol. 143, 48–53 (1996).

18. Ask, H., Rognmo, K., Torvik, F. A., Røysamb, E. & Tambs, K. Non-random mating and convergence over time for alcohol consumption, smoking, and exercise: the Nord-Trøndelag Health Study. Behav. Genet. 42, 354–365 (2012).

19. Cobb, L. K. et al. The association of spousal smoking status with the ability to quit smoking: the Atherosclerosis Risk in Communities Study. Am. J. Epidemiol. 179, 1182–1187 (2014).

20. Burgess, E. W. & Wallin, P. Homogamy in social characteristics. Am. J. Sociol. 49, 109–124 (1943).

21. Alford, J. R., Hatemi, P. K., Hibbing, J. R., Martin, N. G. & Eaves, L. J. The politics of mate choice. J. Polit. 73, 362–379 (2011).

22. Rawlik, K., Canela-Xandri, O. & Tenesa, A. Indirect assortative mating for human disease and longevity. Heredity 123, 106–116 (2019).

23. Price, R. A. & Vandenberg, S. G. Spouse similarity in American and Swedish couples. Behav. Genet. 10, 59–71 (1980).

24. Stulp, G., Simons, M. J. P., Grasman, S. & Pollet, T. V. Assortative mating for human height: a meta-analysis. Am. J. Hum. Biol. 29, e22917 (2017).

25. Allison, D. B. et al. Assortative mating for relative weight: genetic implications. Behav. Genet. 26, 103–111 (1996).

26. Mascie-Taylor, C. G. N. Spouse similarity for IQ and personality and convergence. Behav. Genet. 19, 223–227 (1989).

27. Meyler, D., Stimpson, J. P. & Peek, M. K. Health concordance within couples: a systematic review. Soc. Sci. Med. 64, 2297–2310 (2007).

28. Stimpson, J. P. & Peek, M. K. Concordance of chronic conditions in older Mexican American couples. Prev. Chronic. Dis. 2, (2005).

29. Jeong, S. & Cho, S.-I. Concordance in the health behaviors of couples by age: a cross-sectional study. J. Prev. Med. Pub. Health 51, 6 (2018).

30. Di Castelnuovo, A., Quacquaruccio, G., Donati, M. B., de Gaetano, G. & Iacoviello, L. Spousal concordance for major coronary risk factors: a systematic review and meta-analysis. Am. J. Epidemiol. 169, 1–8 (2009).

31. Hippisley-Cox, J. Married couples’ risk of same disease: cross sectional study. BMJ 325, 636–636 (2002).

32. Galbaud Du Fort, G., Bland, R. C., Newman, S. C. & Boothroyd, L. J. Spouse similarity for lifetime psychiatric history in the general population. Psychol. Med. 28, 789–802 (1998).

33. Peyrot, W. J., Robinson, M. R., Penninx, B. W. J. H. & Wray, N. R. Exploring boundaries for the genetic consequences of assortative mating for psychiatric traits. JAMA Psychiatry 73, 1189–1195 (2016).

34. McLeod, J. D. Social and psychological bases of homogamy for common psychiatric disorders. J. Marriage Fam. 201–214 (1995).

35. Maes, H. H. et al. Assortative mating for major psychiatric diagnoses in two population-based samples. Psychol. Med. 28, 1389–1401 (1998).

36. Mathews, C. A. & Reus, V. I. Assortative mating in the affective disorders: a systematic review and meta-analysis. Compr. Psychiatry 42, 257–262 (2001).

37. Jiang, Y., Bolnick, D. I. & Kirkpatrick, M. Assortative mating in animals. Am. Nat. 181, E125–E138 (2013).

38. Botwin, M. D., Buss, D. M. & Shackelford, T. K. Personality and mate preferences: five factors in mate selection and marital satisfaction. J. Pers. 65, 107–136 (1997).

39. Eysenck, H. J. & Wakefield, J. A. Psychological factors as predictors of marital satisfaction. *Adv*. Behav. Res. Ther. 3, 151–192 (1981).

40. McCrae, R. R. et al. Personality trait similarity between spouses in four cultures. J. Pers. 76, 1137–1164 (2008).

41. Markey, P. M. & Markey, C. N. Romantic ideals, romantic obtainment, and relationship experiences: the complementarity of interpersonal traits among romantic partners. J. Soc. Pers. Relatsh. 24, 517–533 (2007).

42. Al-Sharbatti, S. S., Abed, Y. I., Al-Heety, L. M. & Basha, S. A. Spousal concordance of diabetes mellitus among women in Ajman, United Arab Emirates. Sultan Qaboos Univ. Med. J. 16, e197 (2016).

43. Ober, C. et al. HLA and mate choice in humans. Am. J. Hum. Genet. 61, 497–504 (1997).

44. Keller, M. C., Medland, S. E. & Duncan, L. E. Are extended twin family designs worth the trouble? A comparison of the bias, precision, and accuracy of parameters estimated in four twin family models. Behav. Genet. 40, 377–393 (2010).

45. Cloninger, C. R. Interpretation of intrinsic and extrinsic structural relations by path analysis: theory and applications to assortative mating. Genet. Res. 36, 133–145 (1980).

46. Robinson, M. R. et al. Genetic evidence of assortative mating in humans. *Nat*. Hum. Behav. 1, 1–13 (2017).

47. Yengo, L. et al. Imprint of assortative mating on the human genome. *Nat*. Hum. Behav. 2, 948–954 (2018).

48. Lee, J. J. et al. Gene discovery and polygenic prediction from a genome-wide association study of educational attainment in 1.1 million individuals. Nat. Genet. 50, 1112–1121 (2018).

49. Border, R. et al. Assortative mating biases marker-based heritability estimators. Nat. Commun. 13, 660 (2022).

50. Hartwig, F. P., Davies, N. M. & Davey Smith, G. Bias in Mendelian randomization due to assortative mating. Genet. Epidemiol. 42, 608–620 (2018).

51. Harrer, M., Cuijpers, P., Furukawa, T. A. & Ebert, D. D. Doing Meta-Analysis in R. (2019).

52. Border, R. et al. Cross-trait assortative mating is widespread and inflates genetic correlation estimates. Science 378, 754–761 (2022).

53. Coventry, W. L. & Keller, M. C. Estimating the extent of parameter bias in the classical twin design: a comparison of parameter estimates from extended twin-family and classical twin designs. Twin Res. Hum. Genet. 8, 214–223 (2005).

54. Cloninger, C. R., Rice, J. & Reich, T. Multifactorial inheritance with cultural transmission and assortative mating. II. a general model of combined polygenic and cultural inheritance. Am. J. Hum. Genet. 31, 176–198 (1979).

55. Reczek, C., Pudrovska, T., Carr, D., Thomeer, M. B. & Umberson, D. Marital histories and heavy alcohol use among older adults. J. Health Soc. Behav. 57, 77–96 (2016).

56. Hill, C. T., Rubin, Z. & Peplau, L. A. Breakups before marriage: the end of 103 affairs. J. Soc. Issues 32, 147–168 (1976).

57. Torvik, F. A., Gustavson, K., Røysamb, E. & Tambs, K. Health, health behaviors, and health dissimilarities predict divorce: results from the HUNT study. BMC Psychol. 3, 13 (2015).

58. Fry, A. et al. Comparison of Sociodemographic and Health-Related Characteristics of UK Biobank Participants With Those of the General Population. Am. J. Epidemiol. 186, 1026– 1034 (2017).

59. Kovalchik, S. RISmed: Download Content from NCBI Databases. (2021).

60. Page, M. J. et al. The PRISMA 2020 statement: an updated guideline for reporting systematic reviews. BMJ n71 (2021) doi:10.1136/bmj.n71.

61. Page, M. J. et al. PRISMA 2020 explanation and elaboration: updated guidance and exemplars for reporting systematic reviews. BMJ n160 (2021) doi:10.1136/bmj.n160.

62. Schwartz, C. R. & Graf, N. L. Assortative matching among same-sex and different-sex couples in the United States, 1990–2000. Demogr. Res. 21, 843 (2009).

63. Verbakel, E. & Kalmijn, M. Assortative mating among Dutch married and cohabiting same-sex and different-sex couples. J. Marriage Fam. 76, 1–12 (2014).

64. Nordsletten, A. E. et al. Patterns of nonrandom mating within and across 11 major psychiatric disorders. JAMA Psychiatry 73, 354–361 (2016).

65. Revelle, W. psych: Procedures for Psychological, Psychometric, and Personality Research. (2021).

66. polycor: Polychoric and Polyserial Correlations version 0.8-1 from R-Forge. https://rdrr.io/rforge/polycor/.

67. Viechtbauer, W. metafor: Meta-Analysis Package for R. (2022).

68. Bonett, D. G. & Wright, T. A. Sample size requirements for estimating pearson, kendall and spearman correlations. Psychometrika 65, 23–28 (2000).

69. Ver Hoef, J. M. Who Invented the Delta Method? Am. Stat. 66, 124–127 (2012).

70. Huson, L. W. Performance of Some Correlation Coefficients When Applied to Zero-Clustered Data. J. Mod. Appl. Stat. Methods 6, 530–536 (2007).

