## Supplementary Figures and Note for "Correlations between human mating partners: a comprehensive meta-analysis of 22 traits and raw data analysis of 133 traits in the UK Biobank"

**Supplementary Table S1.** Studies on continuous traits included in the meta-analysis, including sample size, descriptions of sample(s), and (where relevant), measure(s)/instrument(s) used and any covariates used.

**Supplementary Table S2.** Studies on dichotomous traits included in the meta-analysis, including sample size, relevant effect size(s), raw contingency table values not corrected for ascertainment (where available), expected population contingency table values (corrected for any ascertainment), prevalences of the trait in females and in males, descriptions of sample(s), measure(s)/instrument(s) used, and any covariates/matching criteria used.

**Supplementary Table S3.** Citations for studies that were returned in our literature search for the meta-analysis that we excluded, accompanied by each study's reason for exclusion.

**Supplementary Table S4.** Summary statistics and zero order correlations between inferred partners in the UK Biobank for 140 traits, as well as corresponding Field IDs and trait descriptions; summary statistics and partial correlations between inferred partners in the UK Biobank for the same traits controlling for each partner's first ten ancestral principal components, their year of birth, and whether or not they were born within the British Isles, as well as corresponding Field IDs and trait descriptions; discrepancies between zero-order correlations and partial correlations for the same trait.

**Supplementary Table S5.** Statistical comparison of correlations in our UK Biobank partner sample and that of Yengo *et al.* (2018) for each trait examined in both studies.

**Supplementary Table S6.** The list of search terms we used to detect studies related to partner correlations/assortative mating in our literature search, in addition to the terms used in combination with these terms to detect studies related to each meta-analyzed trait/set of traits.

### Table of Contents

**1 Supplementary Figure 1: Partner correlations for 133 UK Biobank traits**…………6

**2 Supplementary Figures 2a-2v: Forest plots, sorted by year of publication and color-coded by region**

**2.1. Smoking Status Forest Plot**………………………………………………...7

**2.2. Smoking Quantity Forest Plot**……………………………………………..8

**2.3. Extraversion Forest Plot**…………………………………………………...9

**2.4. Neuroticism Forest Plot**…………………………………………………..10

**2.5. Openness Forest Plot**……………………………………………………...11

**2.6. Drinking Quantity Forest Plot**……………………………………………12

**2.7. Agreeableness Forest Plot**………………………………………………...13

**2.8. Waist-to-Hip Ratio Forest Plot**…………………………………………...14

**2.9. Depression Forest Plot**…………………………………………………….15

**2.10. Diabetes Forest Plot**………………………………………………………16

**2.11. Generalized Anxiety Forest Plot**…………………………………………17

**2.12. Political Values Forest Plot**………………………………………………18

**2.13. Religiosity Forest Plot**…………………………………………………….19

**2.14. Smoking Initiation Forest Plot**…………………………………………...20

**2.15. Smoking Cessation Forest Plot**…………………………………………...21

**2.16. Problematic Alcohol Use Forest Plot**…………………………………….22

**2.17. Substance Use Disorder Forest Plot**……………………………………...23

**2.18. Body Mass Index Forest Plot**……………………………………………..24

**2.19. Height Forest Plot**……………………………………………………………25

**2.20. Educational Attainment Forest Plot**………………………………………..26

**2.21. Intelligence Quotient Score Forest Plot**…………………………………….27

**2.22. Conscientiousness Forest Plot**……………………………………….........28

**3 Supplementary Figure 3: Funnel Plots**

**3.1. Funnel Plots for Smoking Status, Height, Smoking Quantity, Extraversion, Neuroticism, Openness, Conscientiousness, Drinking Quantity, Agreeableness, Intelligence Quotient Score, Waist-to-Hip Ratio, and Educational Attainment**……………………………………………………………………….29

**3.2. Funnel Plots for Depression, Diabetes, Generalized Anxiety, Political Values, Religiosity, Smoking Initiation, Smoking Cessation, Problematic Alcohol Use, Substance Use Disorder, and Body Mass Index**………………………………….30

**4 Supplementary Figure 4: PRISMA Flow Diagrams**

**4.1. Substance Use Disorder Flow Diagram**…………………………………….31

**4.2. Problematic Alcohol Use Flow Diagram**…………………………………...32

**4.3. Drinking Quantity Flow Diagram**…………………………………………..33

**4.4. Waist-to-Hip Ratio Flow Diagram**..………………………………………...34

**4.5. Height Flow Diagram**……………………………………………………......35

**4.6. Body Mass Index Flow Diagram**……………………………………………36

**4.7. Extraversion Flow Diagram**………………………………………………...37

**4.8. Neuroticism Flow Diagram**…………………………………………………38

**4.9. Openness Flow Diagram**…………………………………………………….39

**4.10. Conscientiousness Flow Diagram**………………………………………….40

**4.11. Agreeableness Flow Diagram**……………………………………………41

**4.12. Intelligence Quotient Score Flow Diagram.**………………………………42

**4.13. Diabetes Flow Diagram**……………………………………………………..43

**4.14. Educational Attainment Flow Diagram**………………………………...44

**4.15. Smoking Status Flow Diagram**…………………………………………….45

**4.16. Smoking Initiation Flow Diagram**………………………………………46

**4.17. Smoking Cessation Flow Diagram**………………………………………47

**4.18. Smoking Quantity Flow Diagram**……………………………………….48

**4.19. Religiosity Flow Diagram**…………………………………………………..49

**4.20. Political Values Flow Diagram**……………………………………...……..50

**4.21. Depression Flow Diagram**………………………………………………….51

**4.22. Generalized Anxiety Flow Diagram**…………………………………….52

**5 Supplementary Note**

**5.1. Trait descriptions for Educational Attainment, Intelligence Quotient Score, Political Values, Religiosity, Problematic Alcohol Use, Drinking Quantity, and Smoking Cessation**……………………………………………………………53-54

**5.2. Trait descriptions for Smoking Initiation, Smoking Quantity, Smoking Status, Substance Use Disorder, and Agreeableness**……………………….54-55

**5.2. Trait descriptions for Conscientiousness, Extraversion, Neuroticism, and Openness**……………………………………………………………………55-56

**5.3. Trait descriptions for Body Mass Index, Height, Waist-to-Hip Ratio, Depression, Diabetes, and Generalized Anxiety**……………………….…56-57

**Supplementary Figure 1: Partner correlations and 95% confidence intervals for 133 traits in the UK Biobank**

The visualized traits represent partner correlations for all of the adequately-powered UK Biobank traits (out of an original 140). Each estimate is color-coded by the correlation type—Pearson (in blue), Spearman (in red), and tetrachoric (in green), used for continuous, ordinal, and binary traits, respectively—with the lines depicting the 95% confidence interval for each trait (the majority of which do not extend beyond the dot). Num Dep Episodes = Number of Depressive Episodes; Heel BMD = Heel Bone Mineral Density (in the form of a t-score); LDL = Direct Low-density Lipoprotein Cholesterol, CRP = C-reactive Protein; RBC = Red Blood Cell (Erythrocyte) Count; DBP = Diastolic Blood Pressure; CPD (All Participants) = Cigarettes per Day (Includes Current, Former, and Never Smokers); FEV1 Pred % = Forced Expiratory Volume in 1-Second (FEV1), Predicted Percentage; PEF= Peak Expiratory Flow; WBC = White Blood Cell (Leukocyte) Count; SBP = Systolic Blood Pressure; HDL = High-density Lipoprotein Cholesterol; CPD (Smokers Only) = Cigarettes per Day (Restricted to Current or Former Smokers); WHR = Waist-to-hip Ratio; BMR = Basal Metabolic Rate; FIQ = Fluid Intelligence Quotient; BMI = Body Mass Index; FVC = Forced Vital Capacity; Time to First Cig = Time to First Cigarette; EA = Educational Attainment

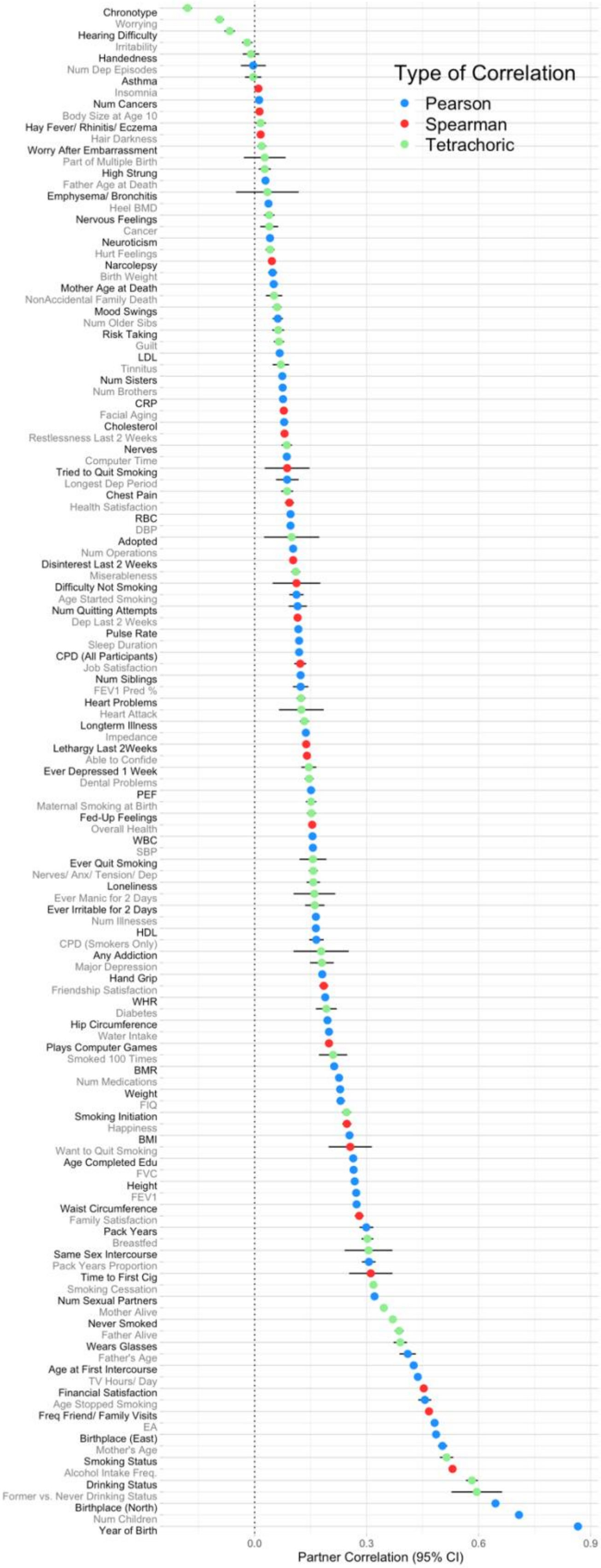

**Supplementary Figure 2a**

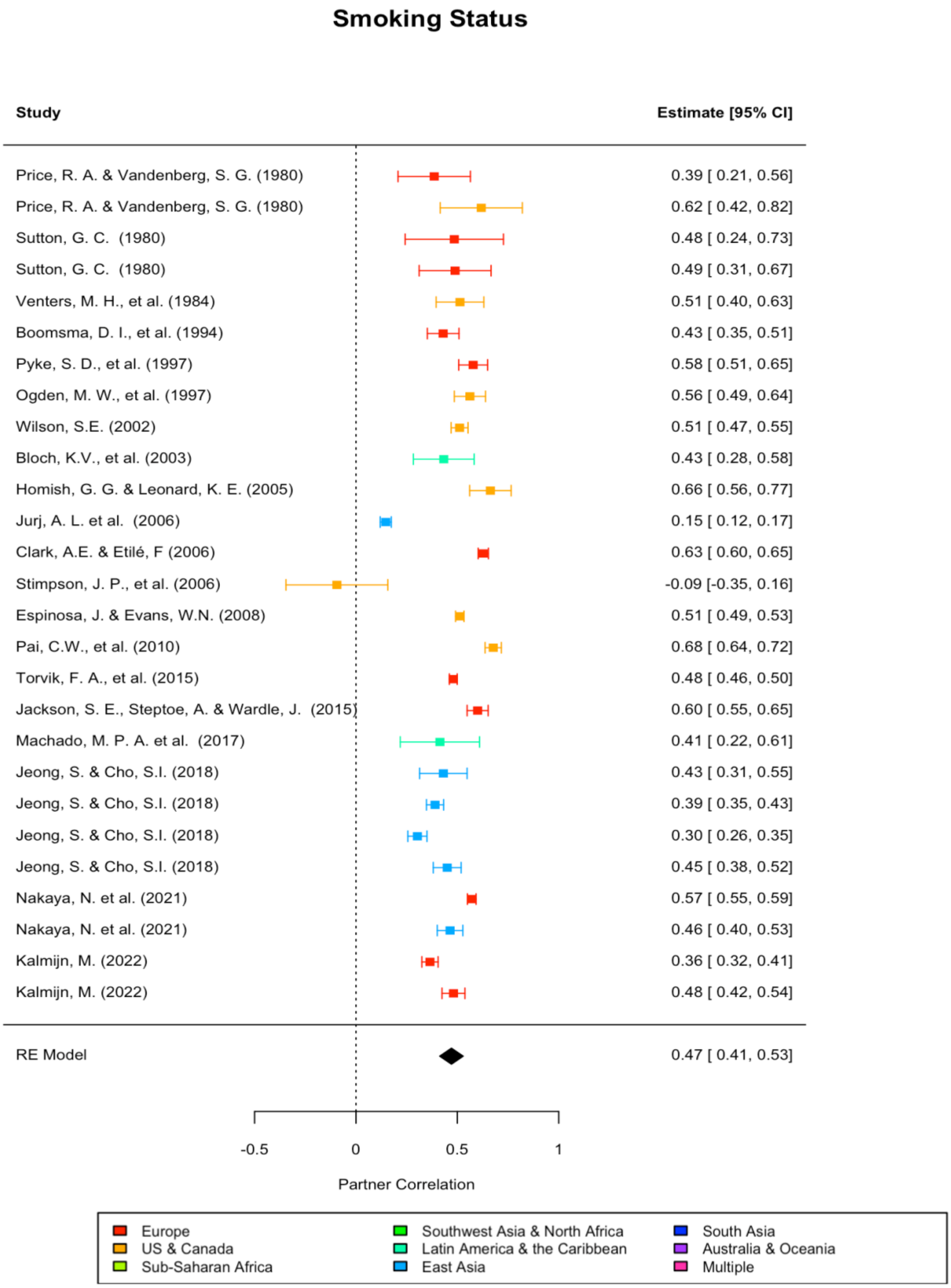

**Supplementary Figure 2b**

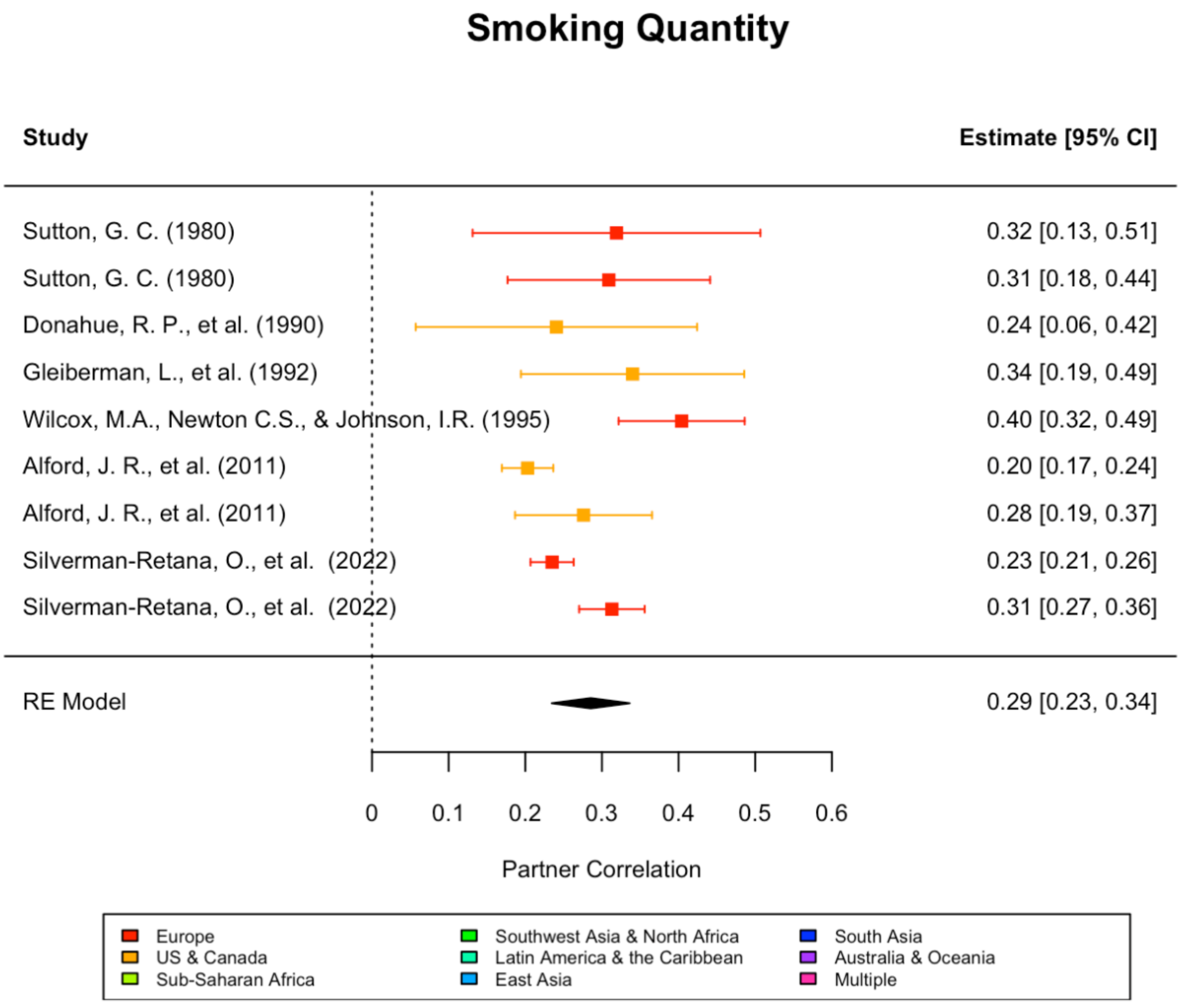

**Supplementary Figure 2c**

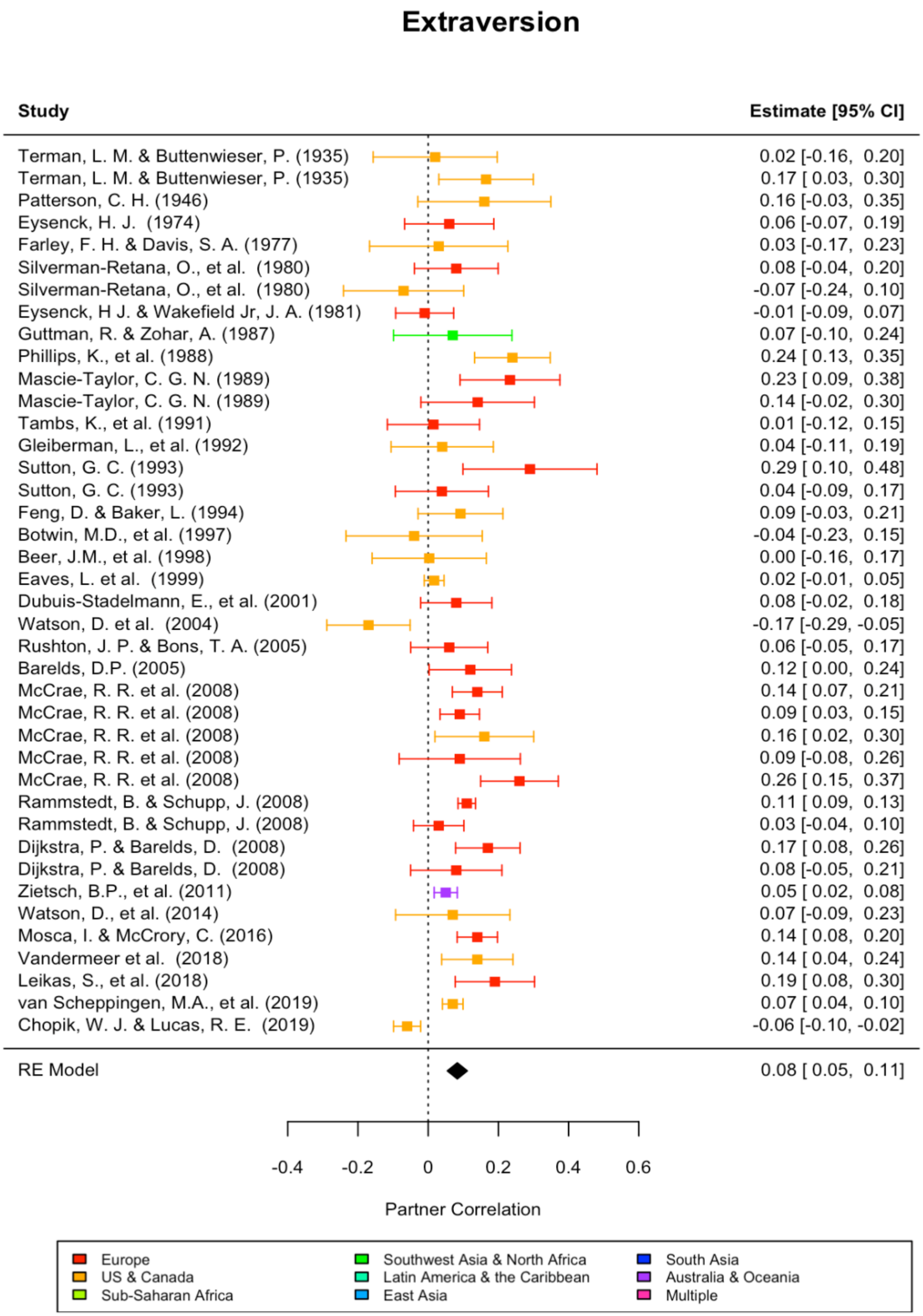

**Supplementary Figure 2d**

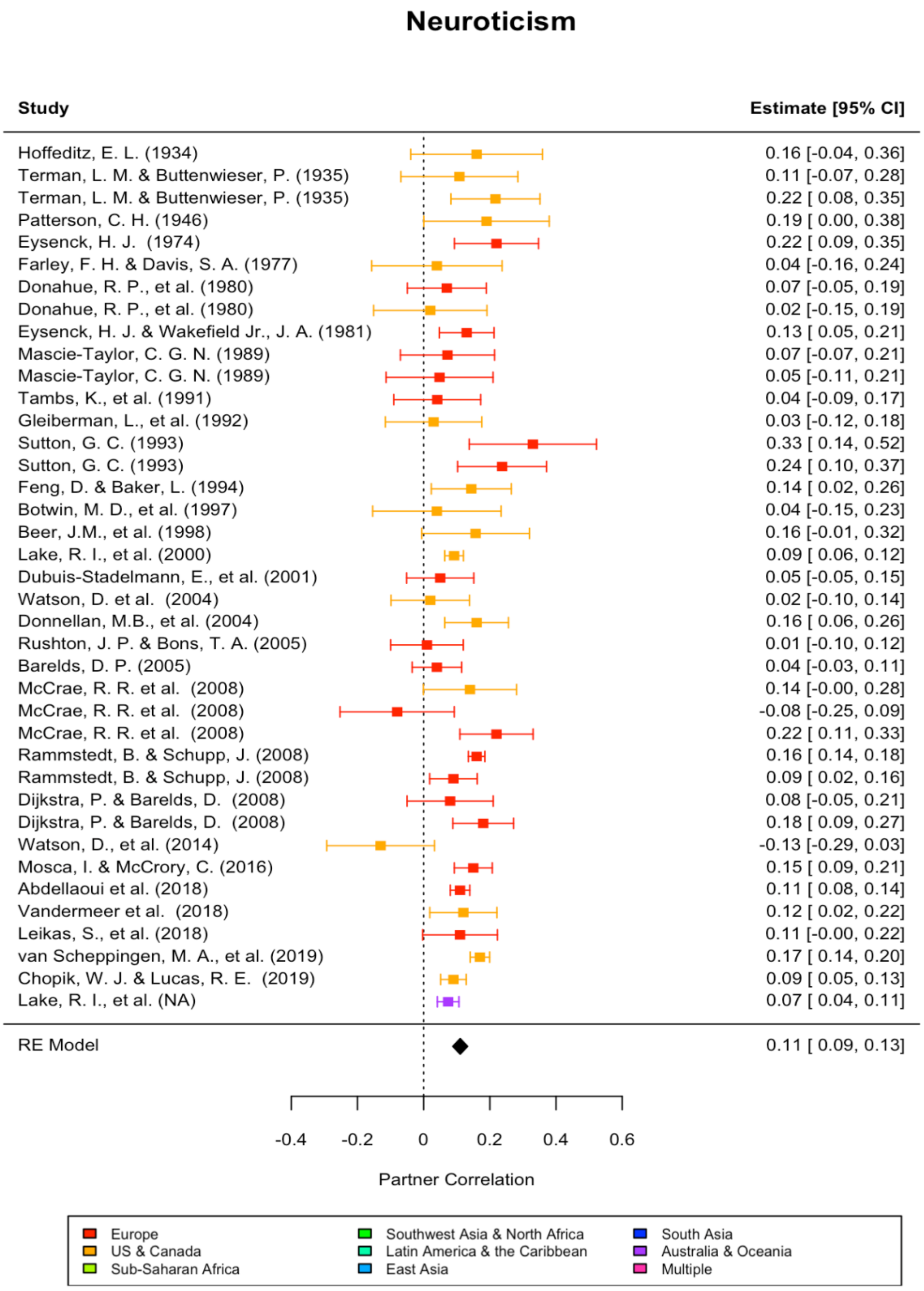

**Supplementary Figure 2e**

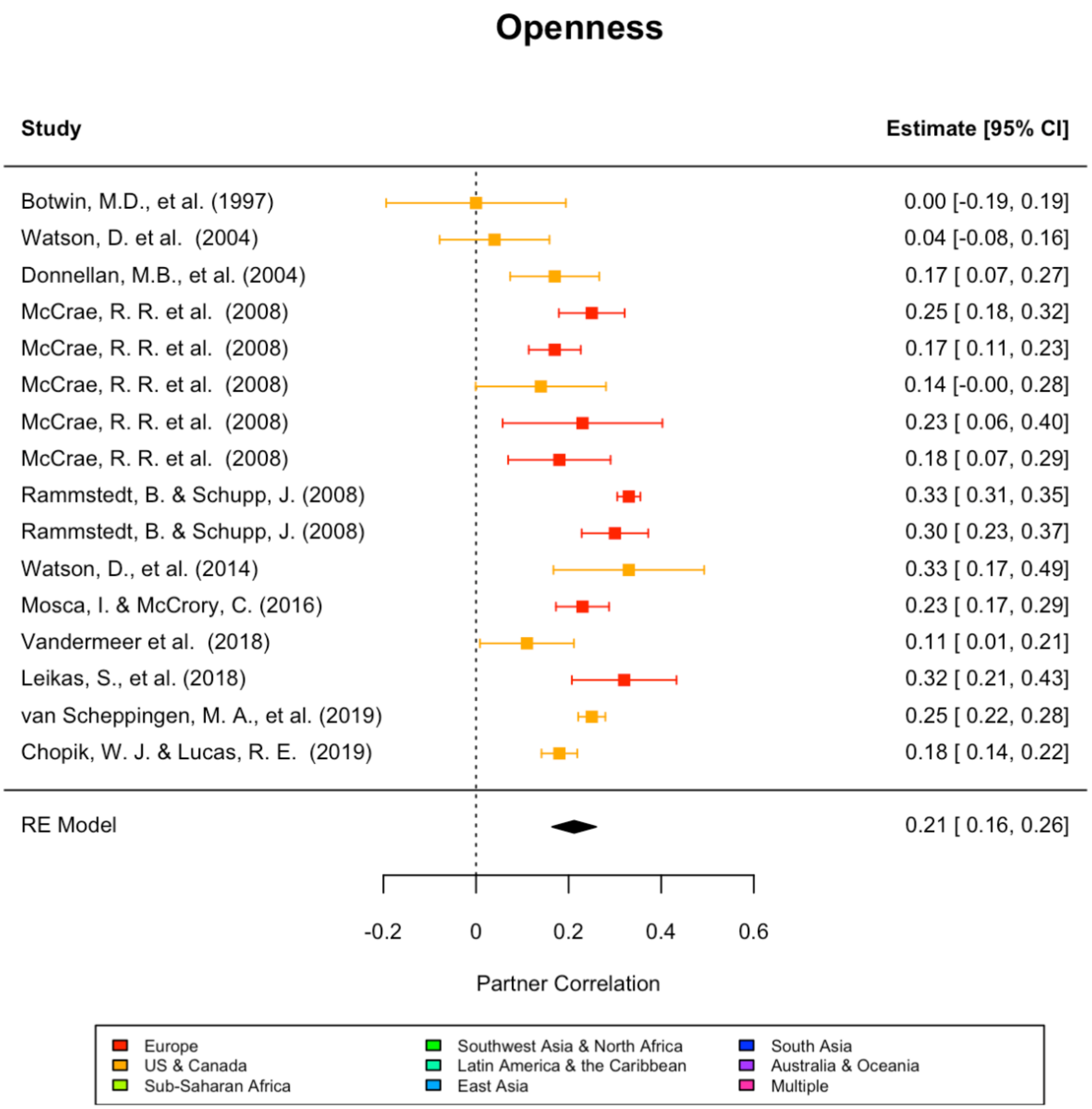

**Supplementary Figure 2f**

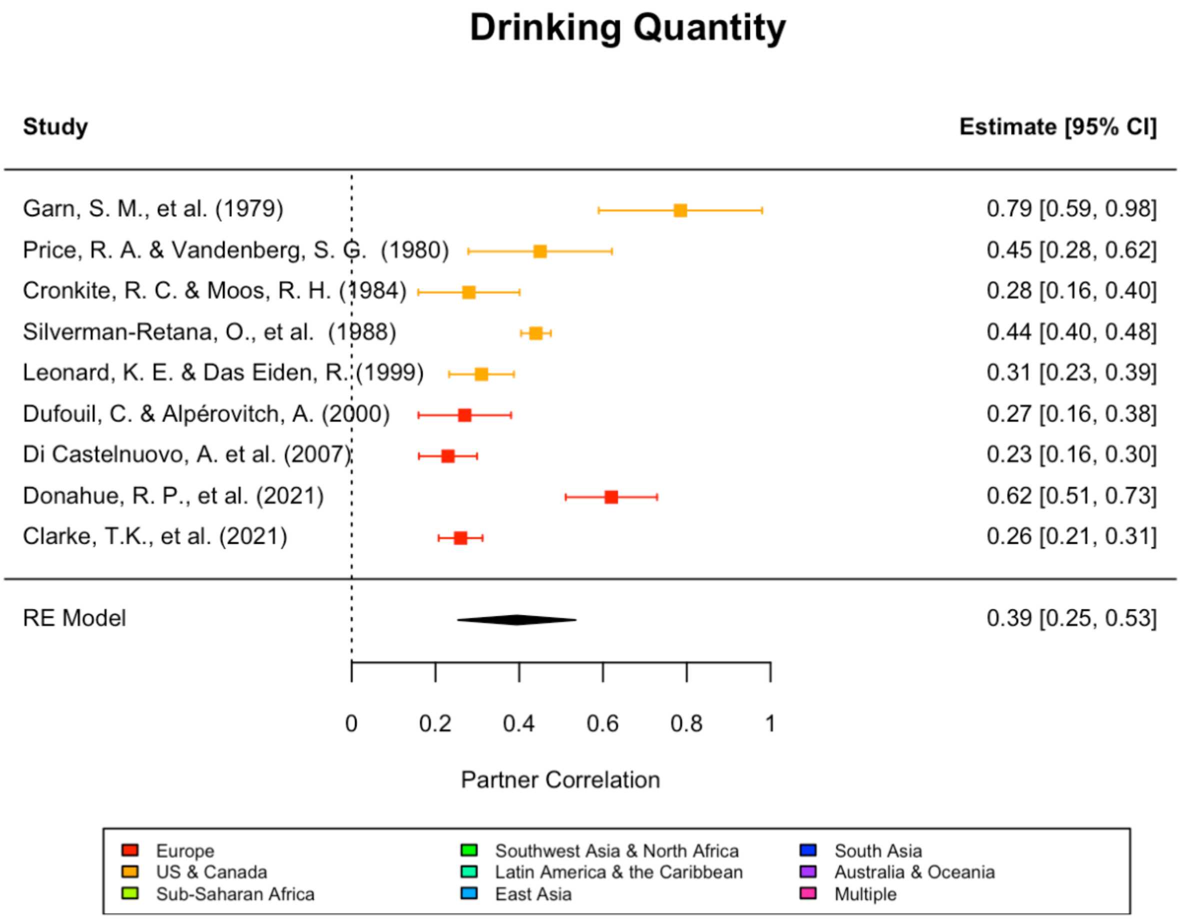

**Supplementary Figure 2g**

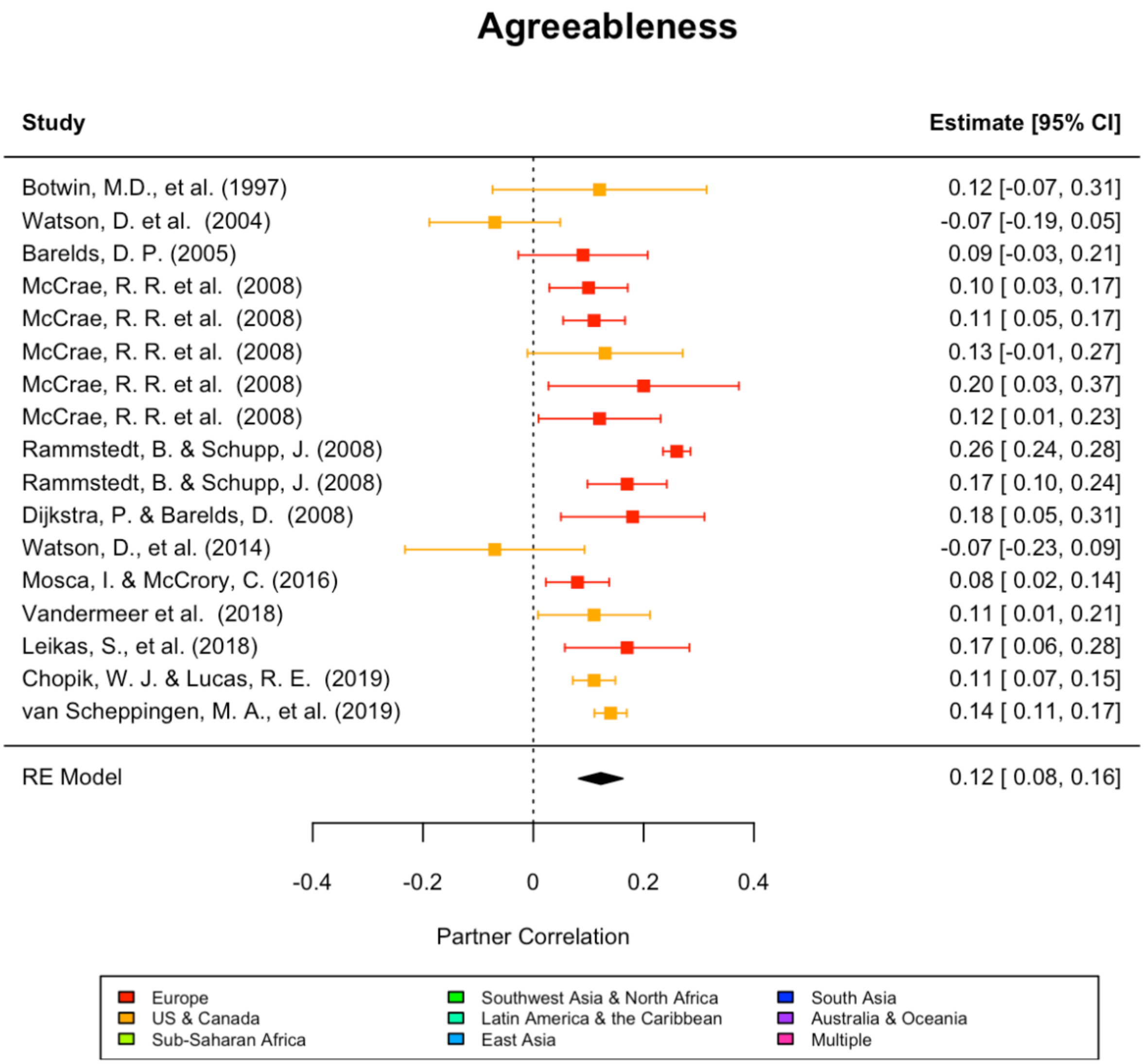

**Supplementary Figure 2h**

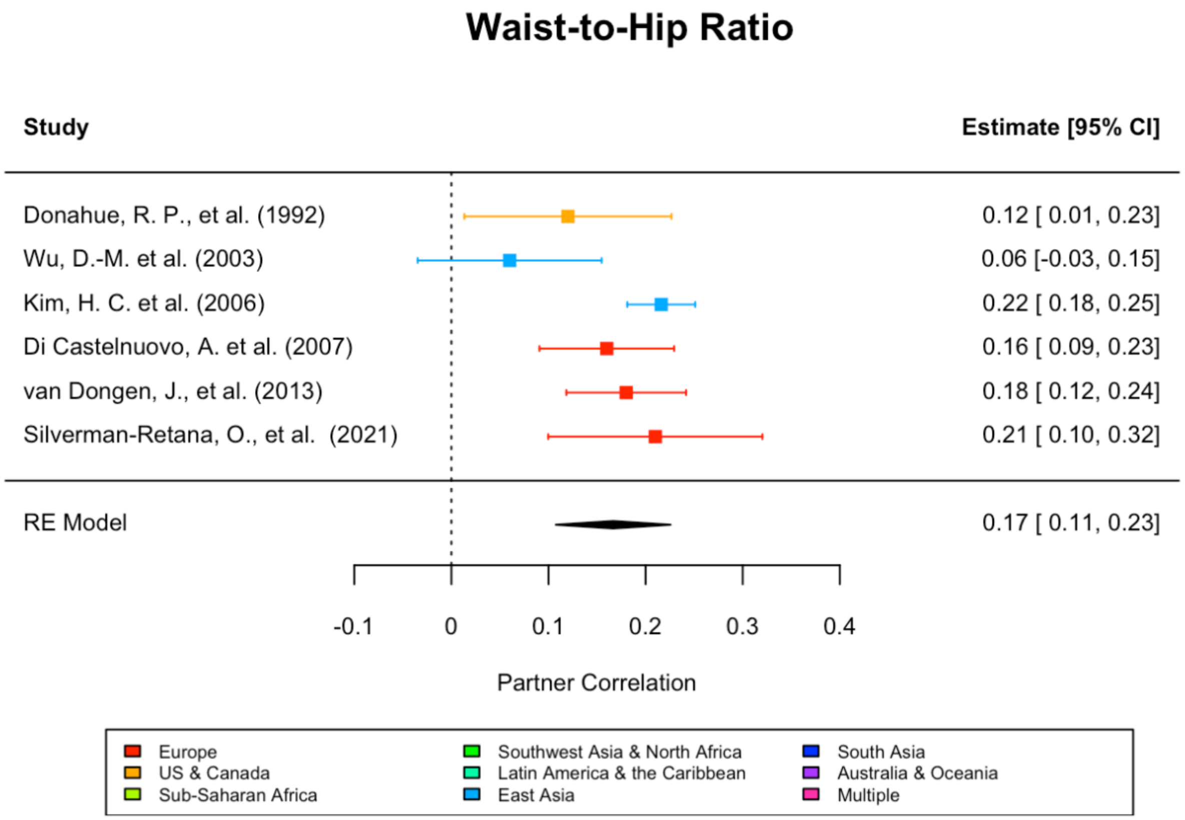

**Supplementary Figure 2i**

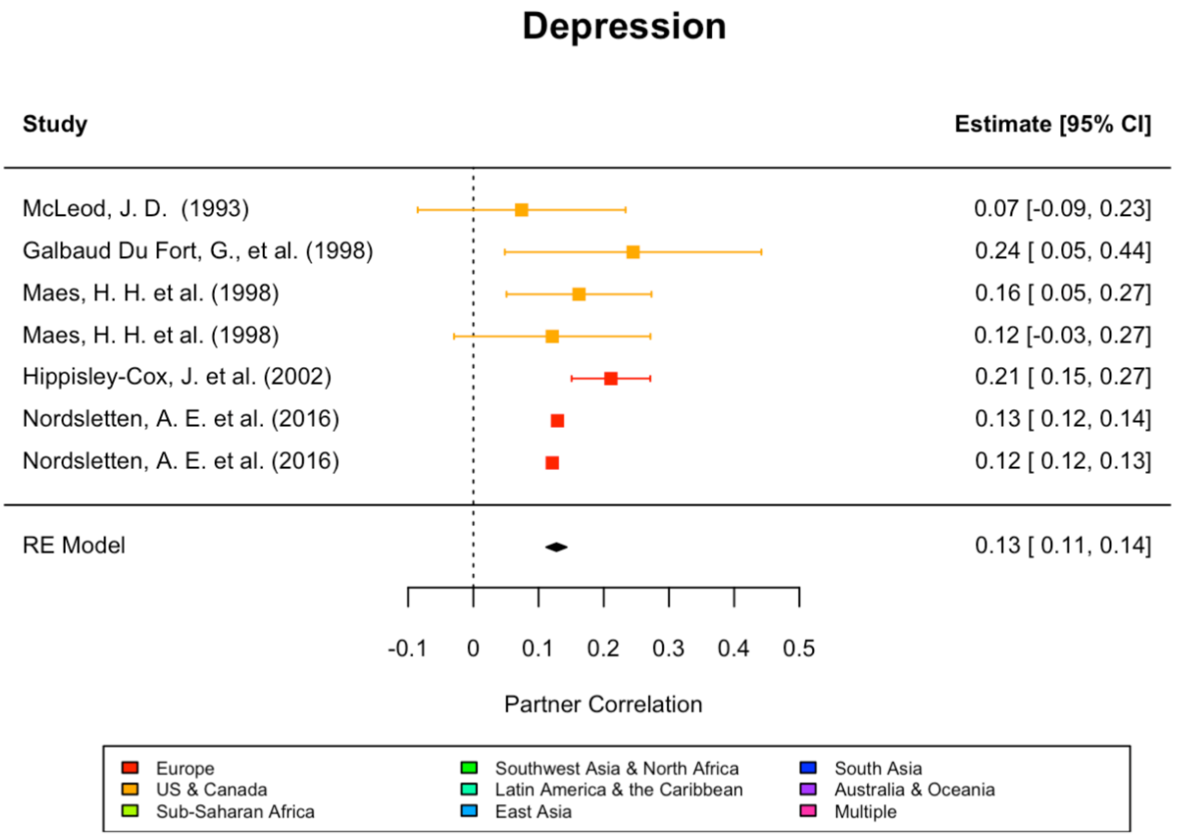

**Supplementary Figure 2j**

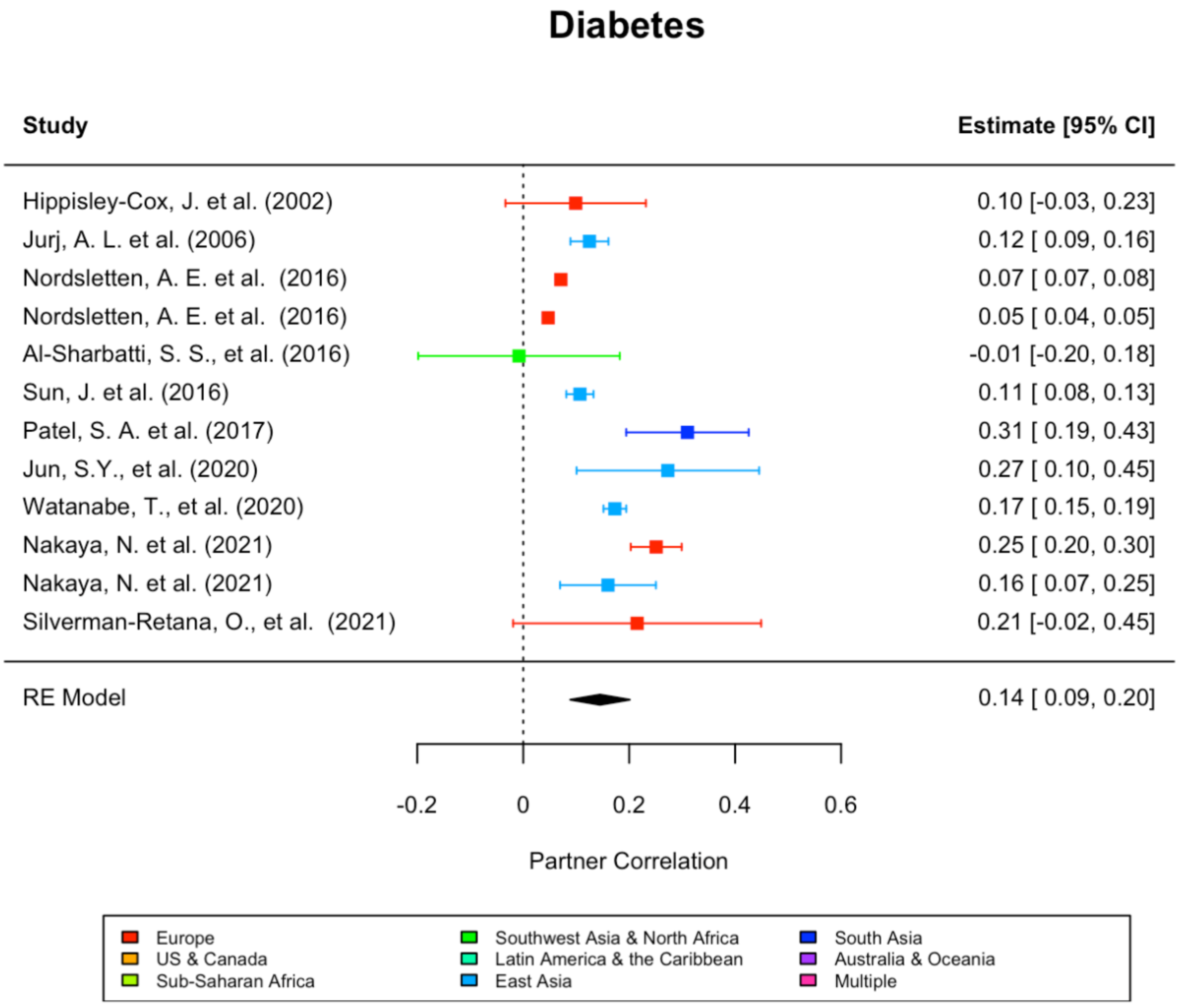

**Supplementary Figure 2k**

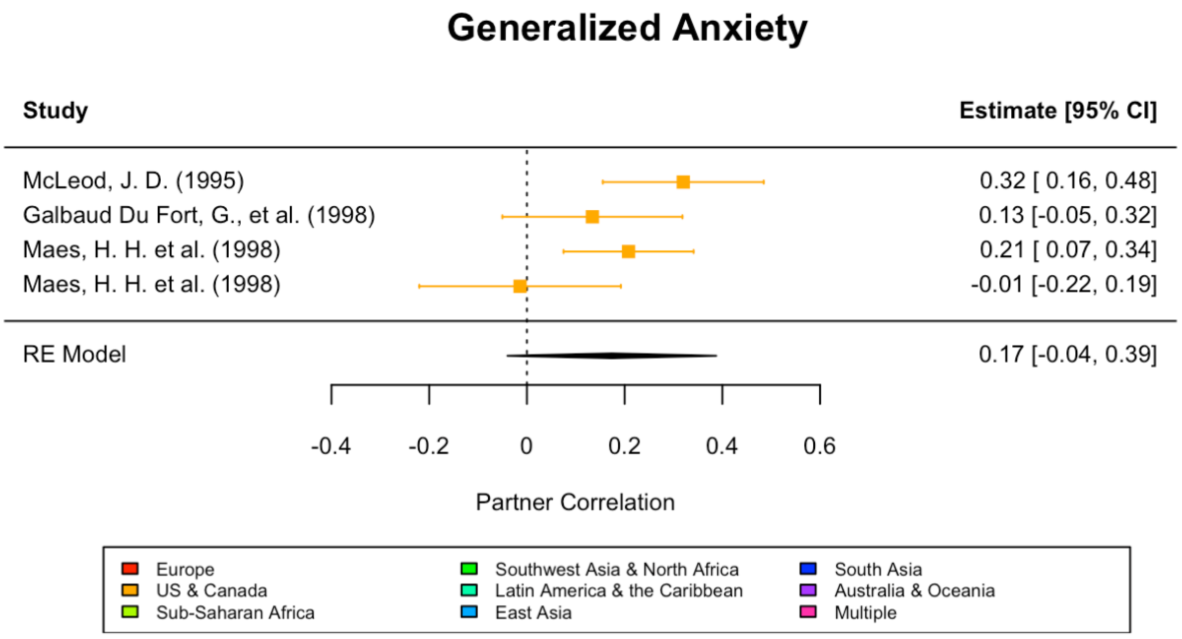

**Supplementary Figure 2l**

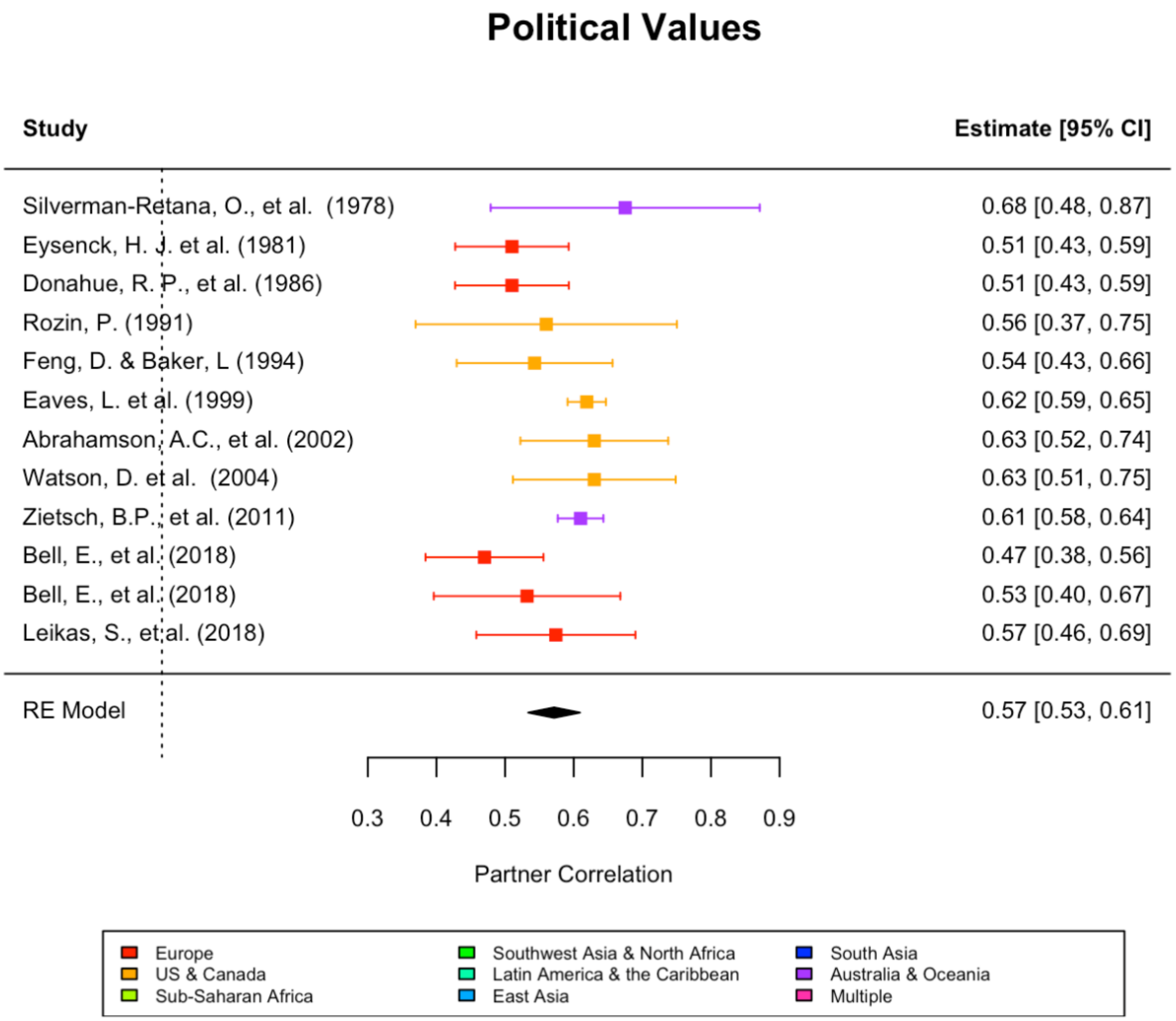

**Supplementary Figure 2m**

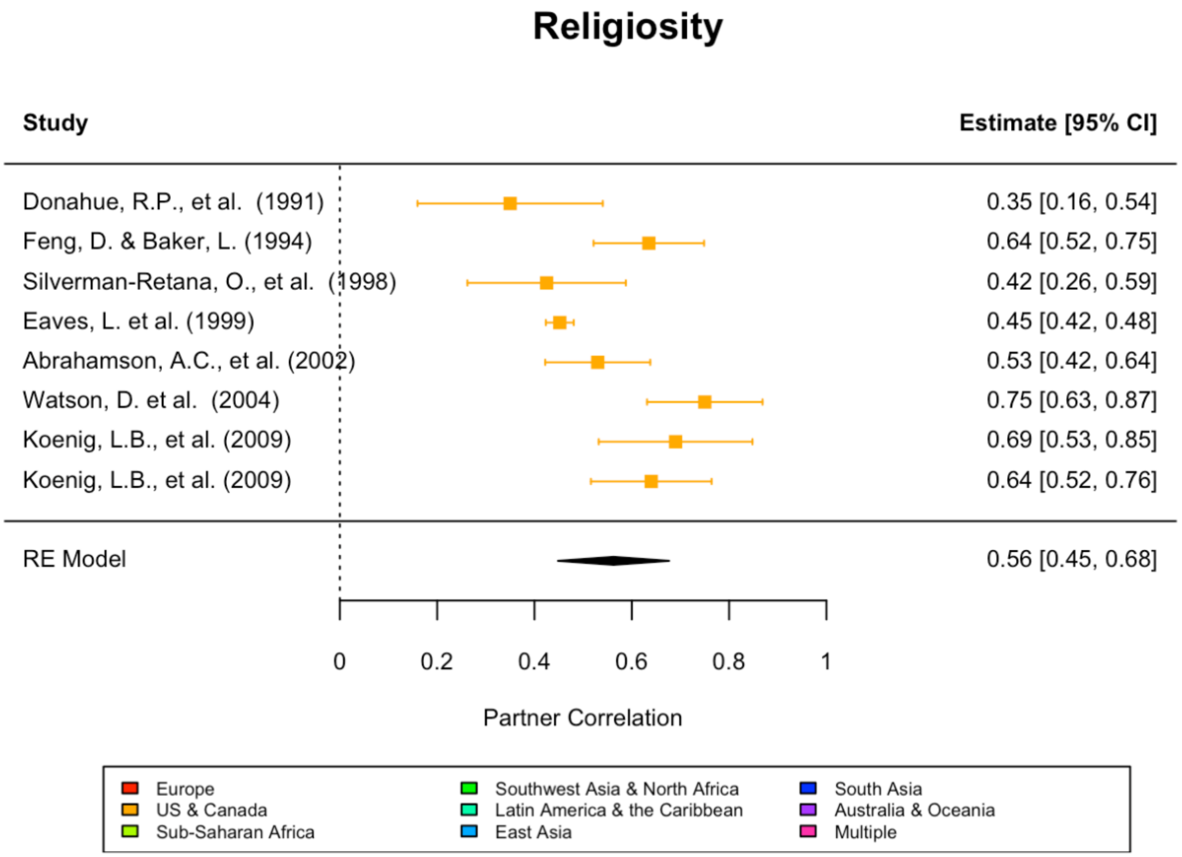

**Supplementary Figure 2n**

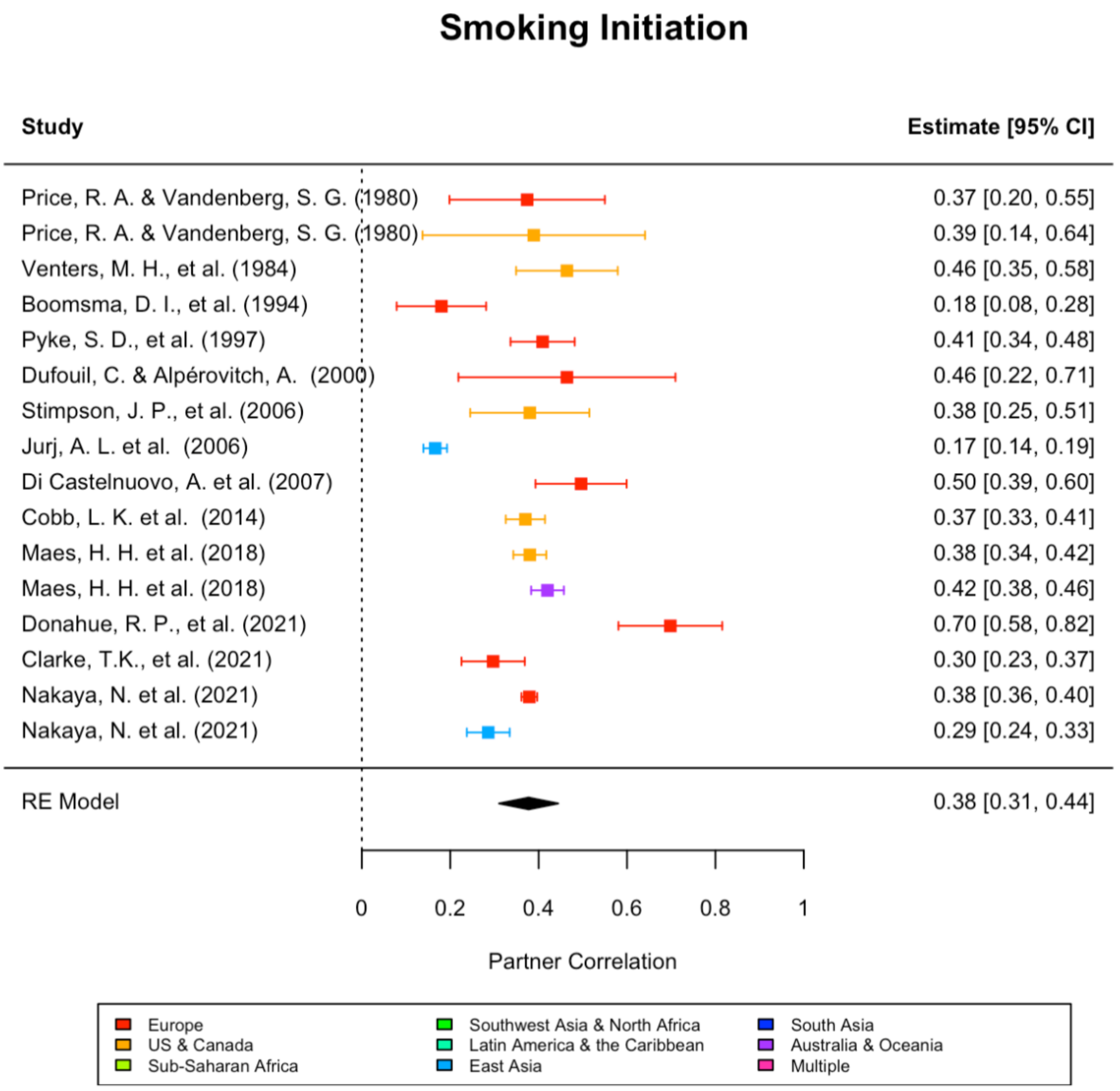

**Supplementary Figure 2o**

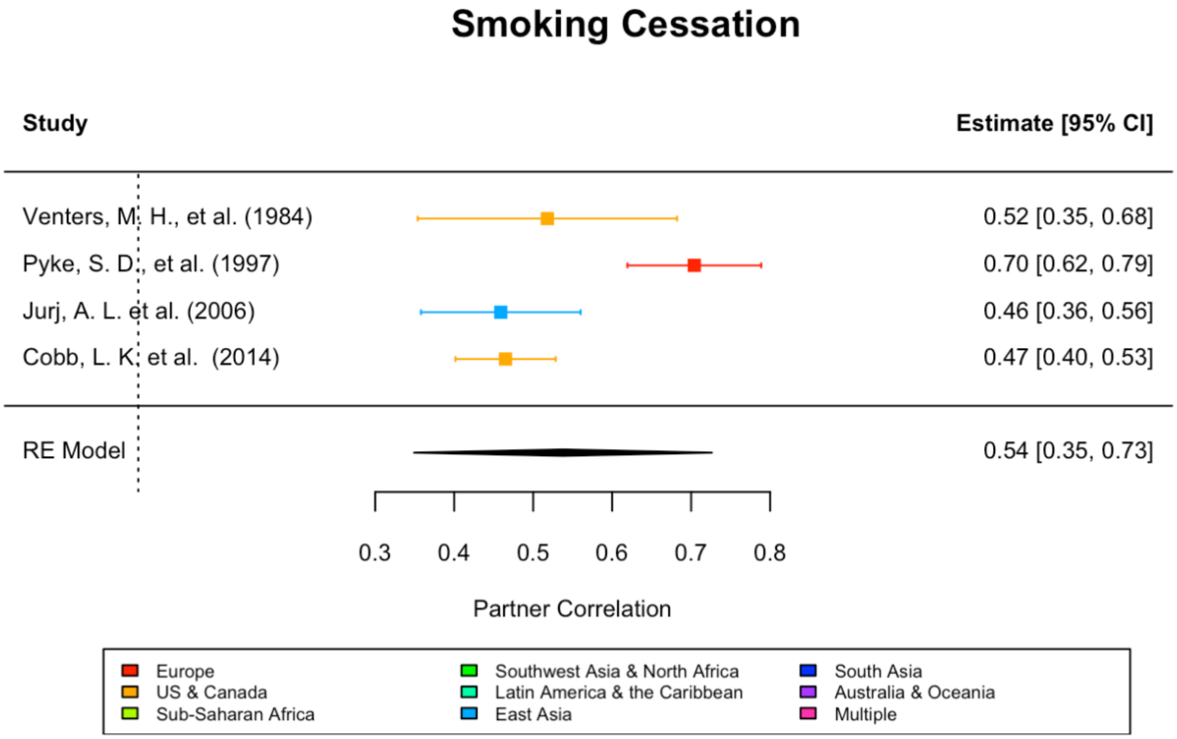

**Supplementary Figure 2p**

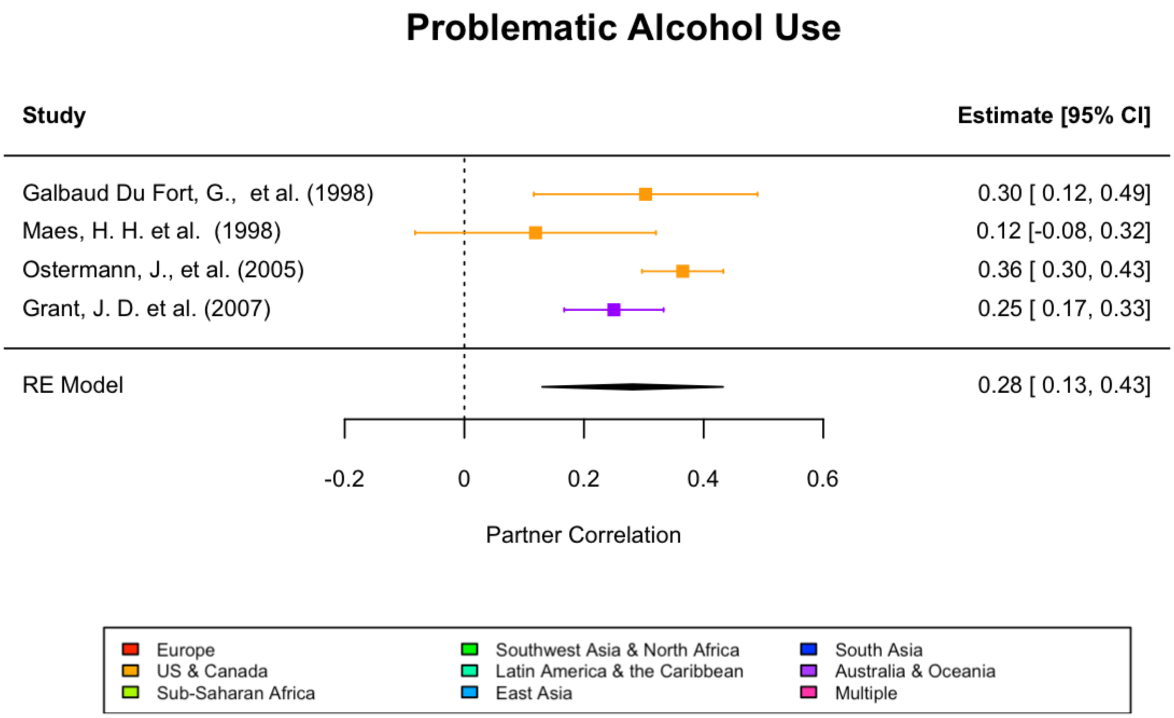

**Supplementary Figure 2q**

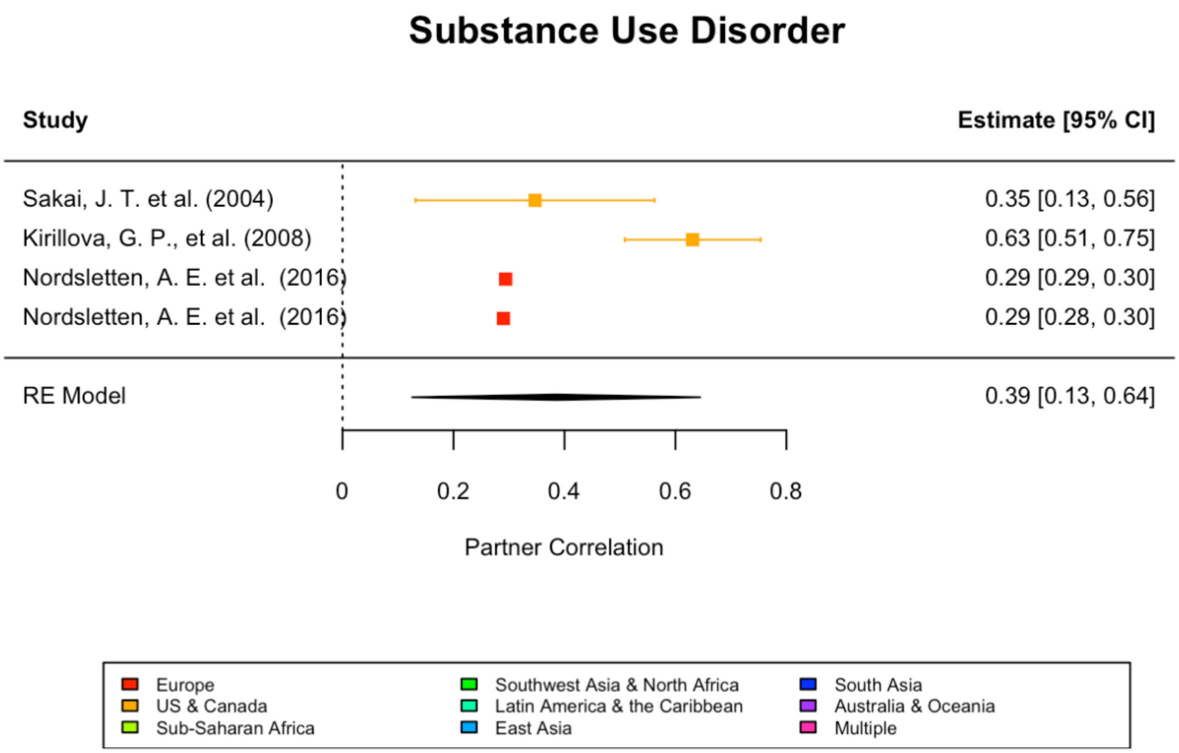

193 **Supplementary Figure 2r**

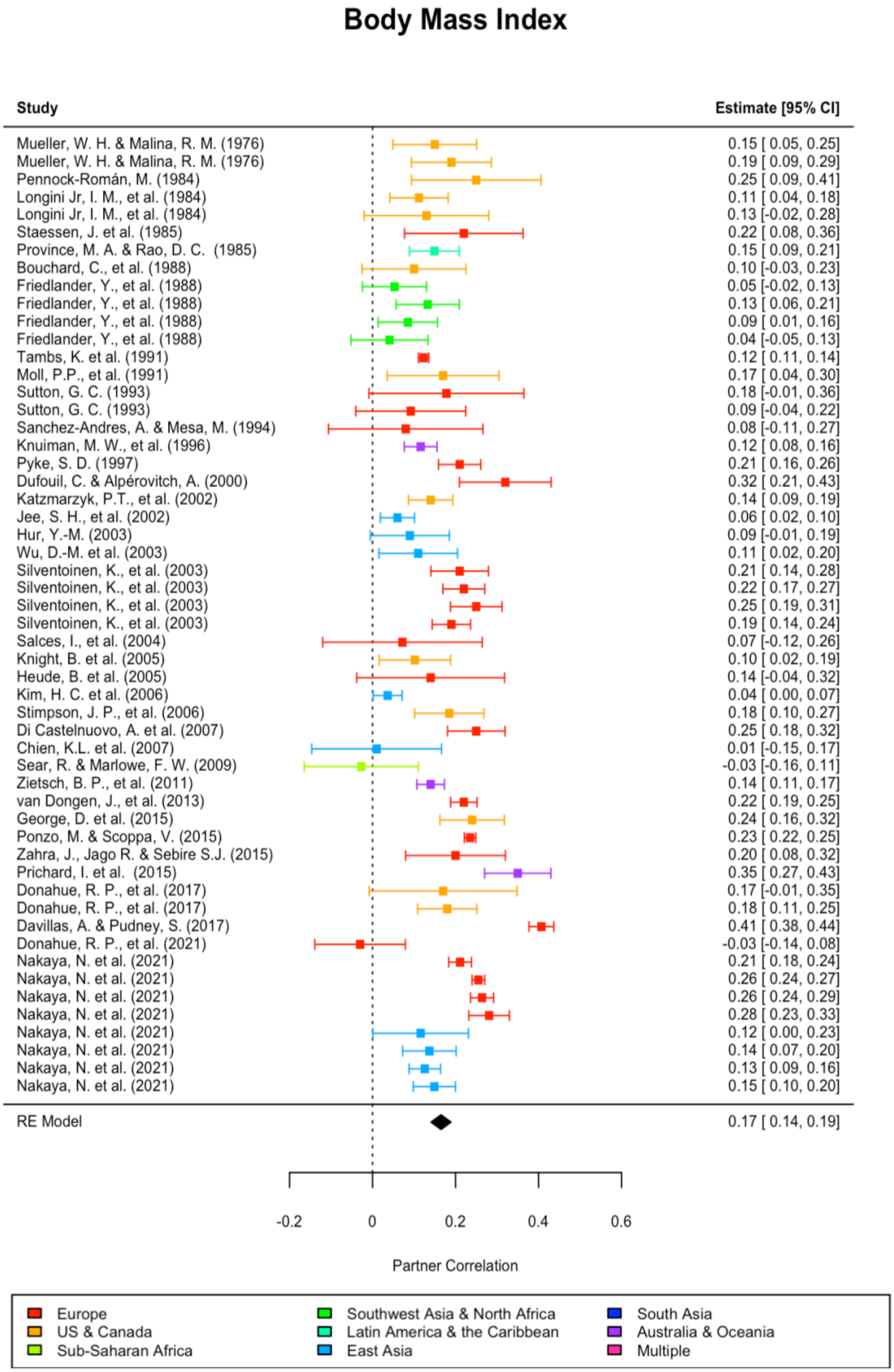


194

195

**Supplementary Figure 2s**

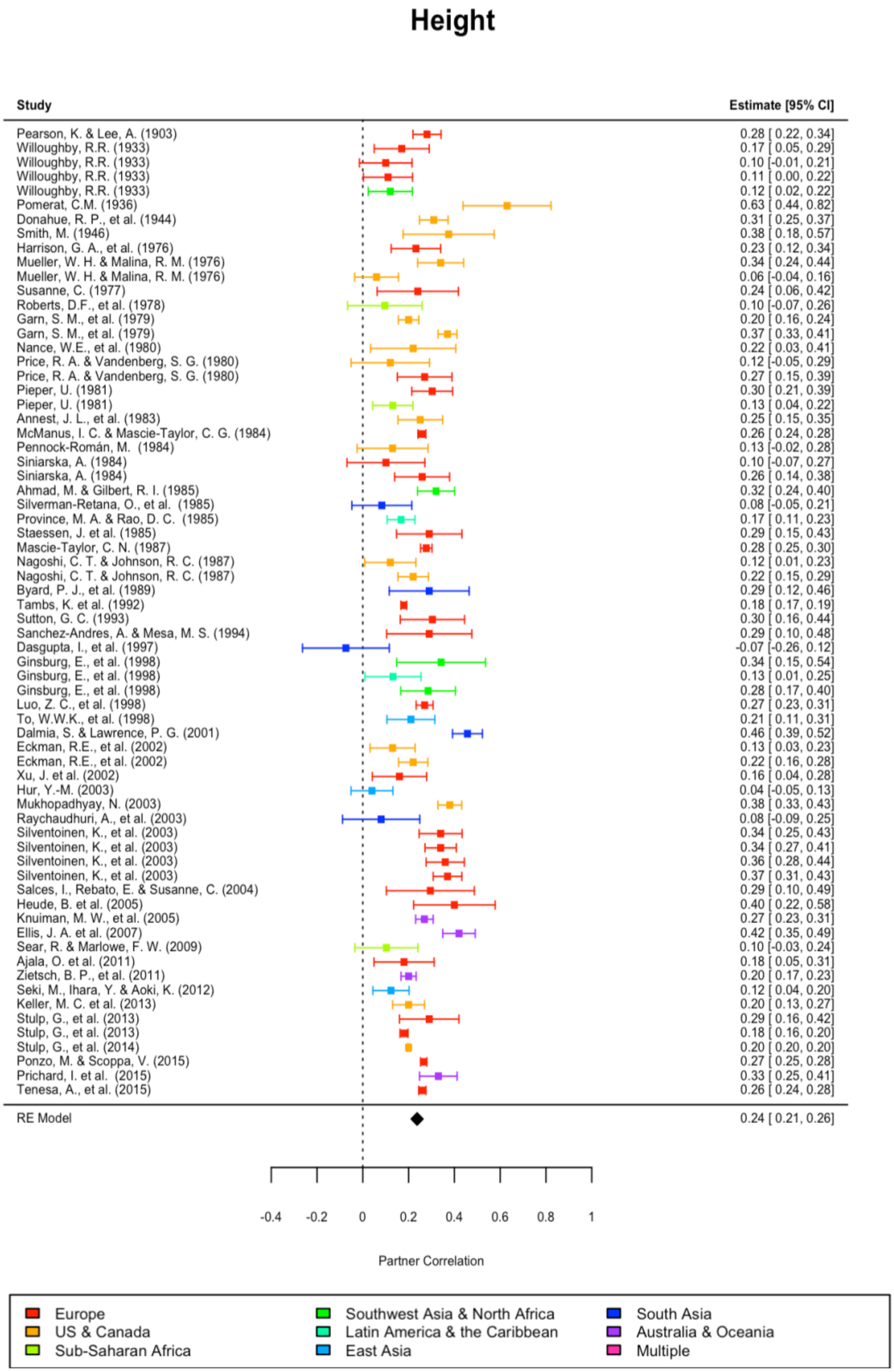

#### Supplementary Figure 2t

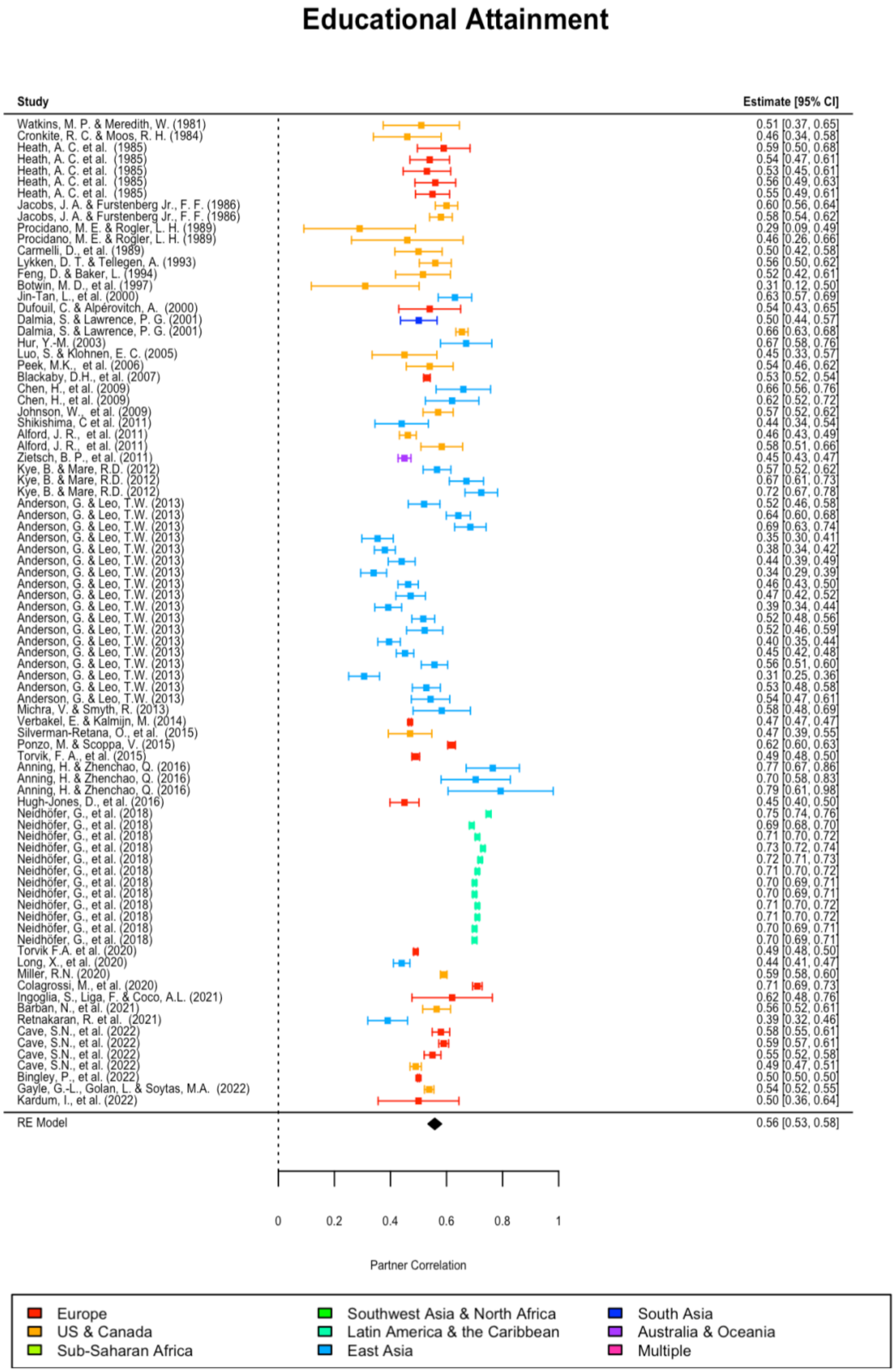

**Supplementary Figure 2u**

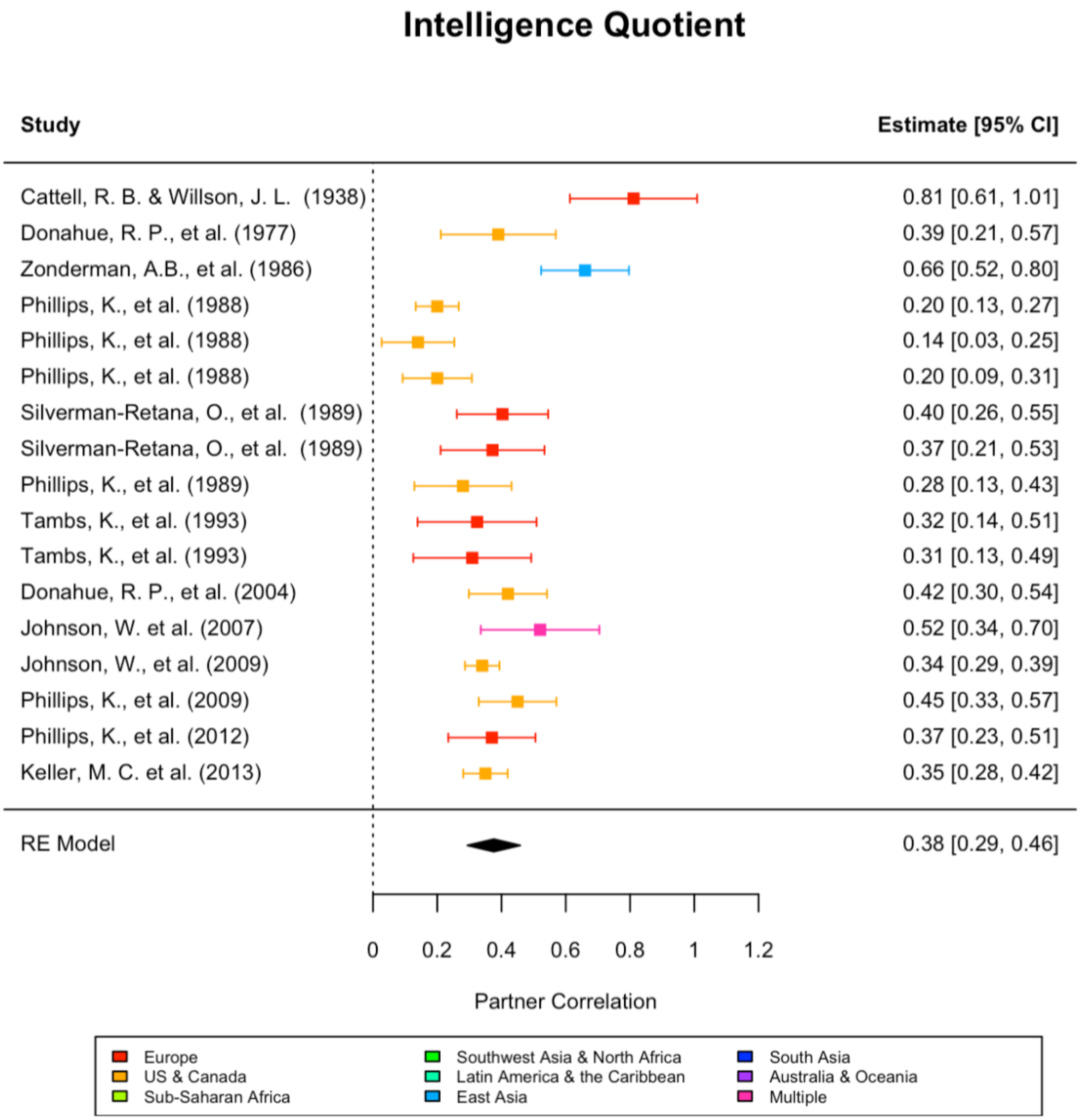

**Supplementary Figure 2v**

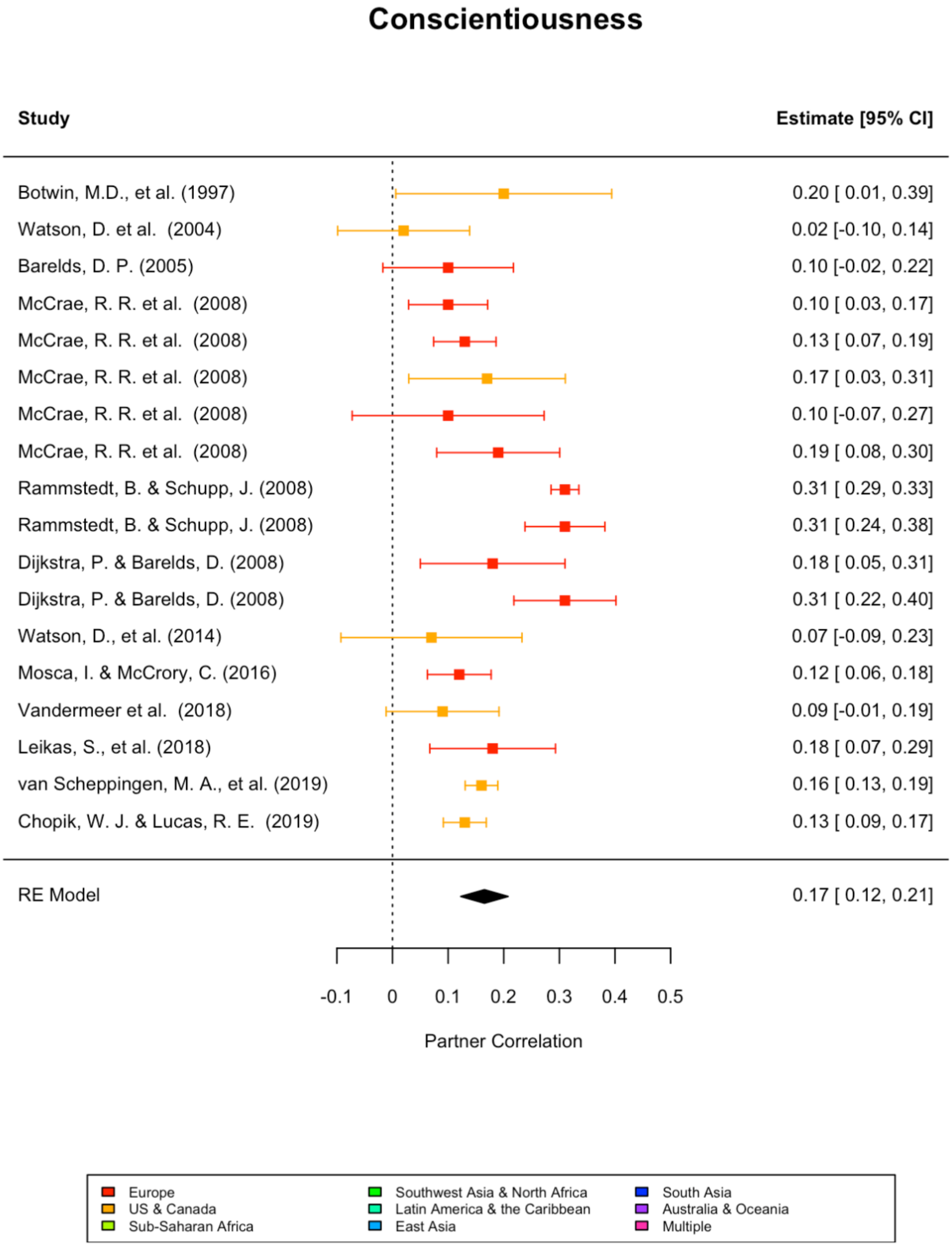


Estimates within each phenotype are sorted by year of publication (from top to bottom) and are color-coded to indicate the region from which the sample was taken, as indicated in the legend

**Supplementary Figures 3: Funnel plots of each meta-analyzed trait.**

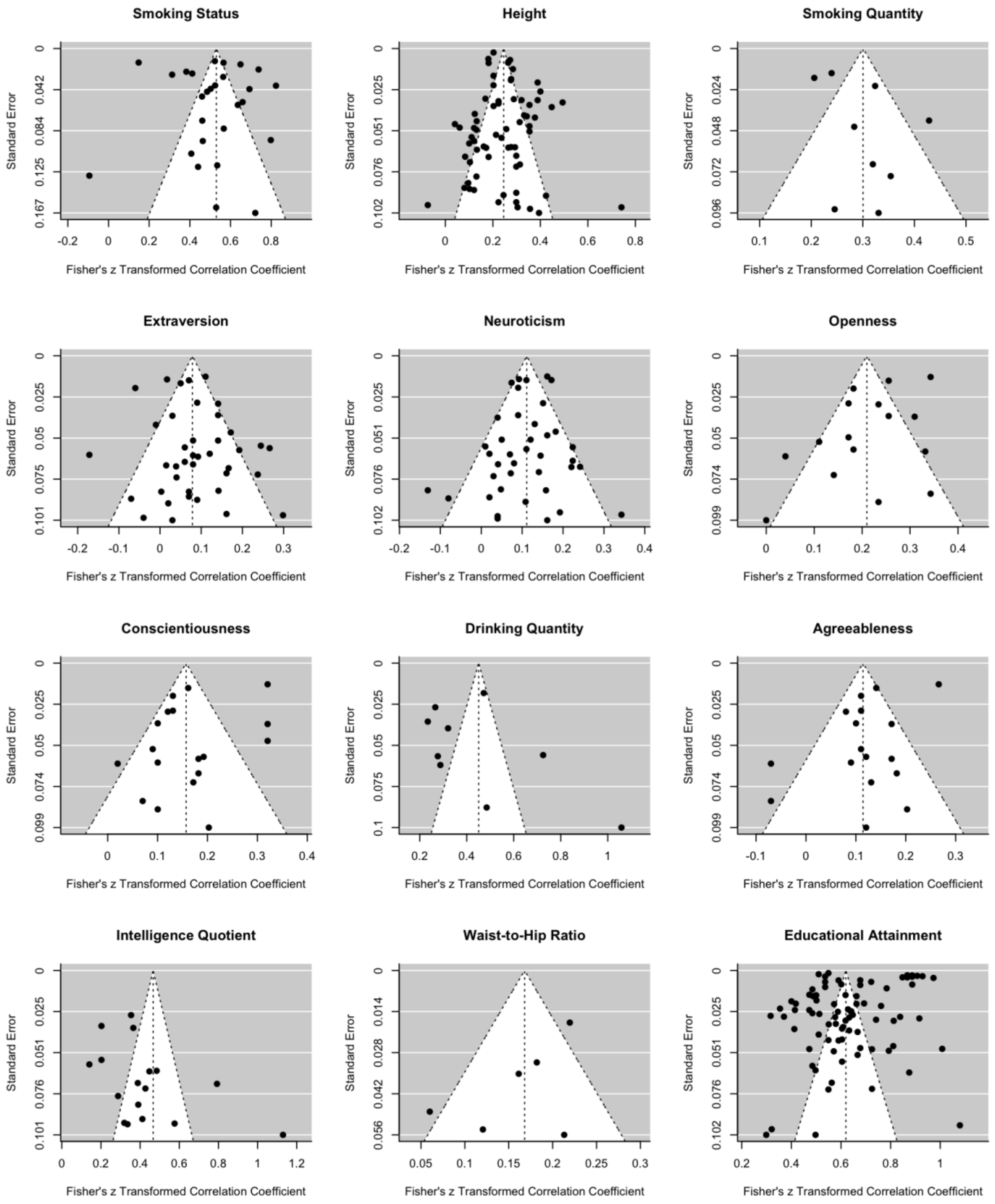

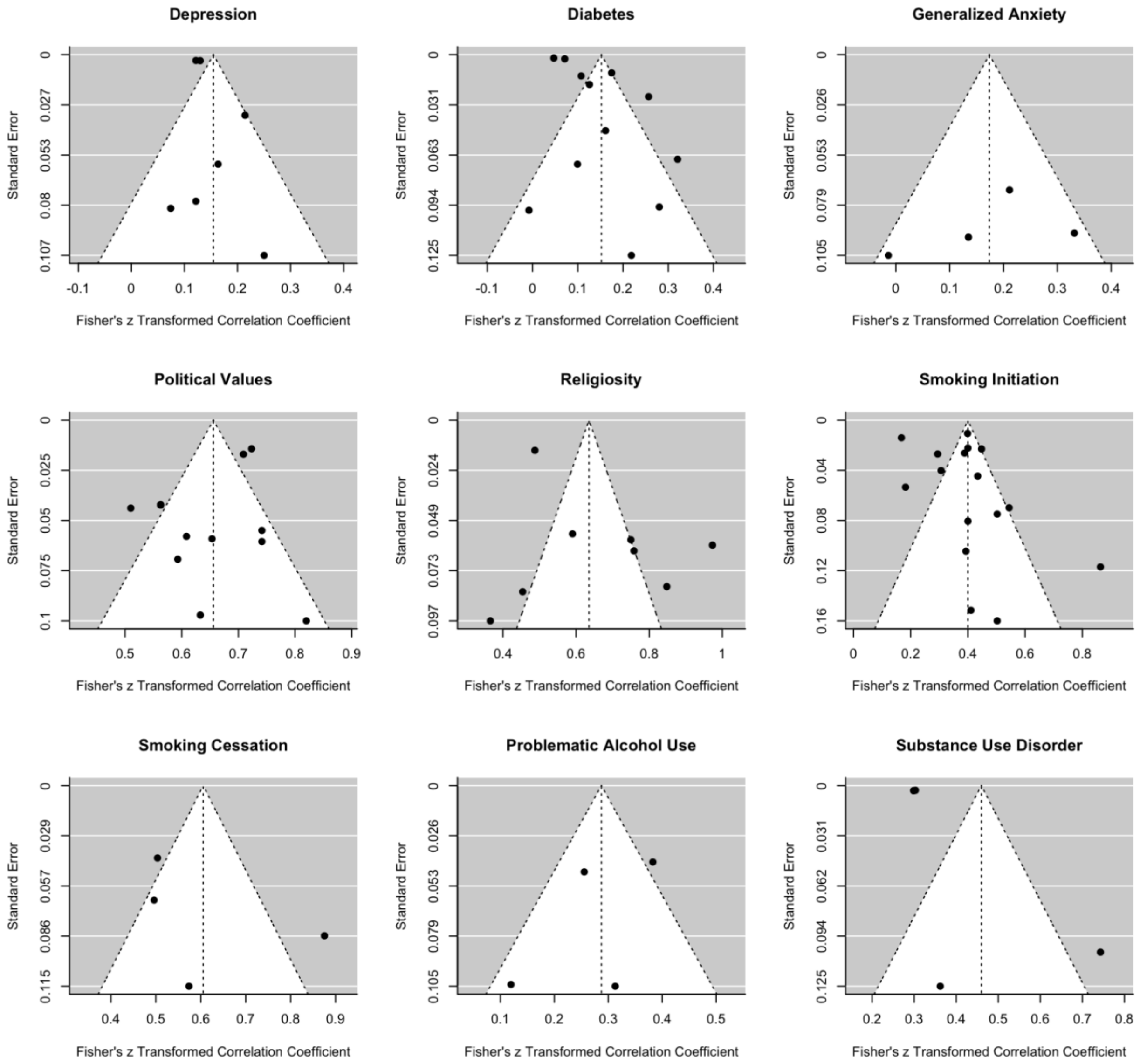


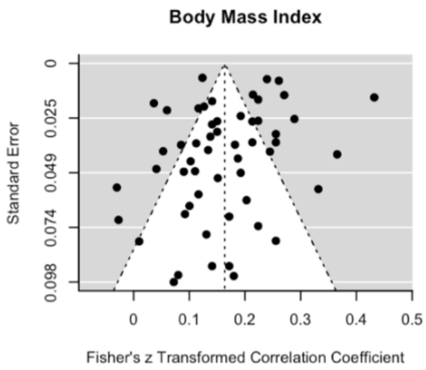


Funnel plots are designed to assess possible publication bias for each study. Here, the Fisher Z-transformed correlations are plotted against their respective transformed standard errors. For dichotomous traits, standard error was estimated using the delta method (see main text).

**Supplementary Figure 4: PRISMA flow diagrams.**

**Substance use disorder**

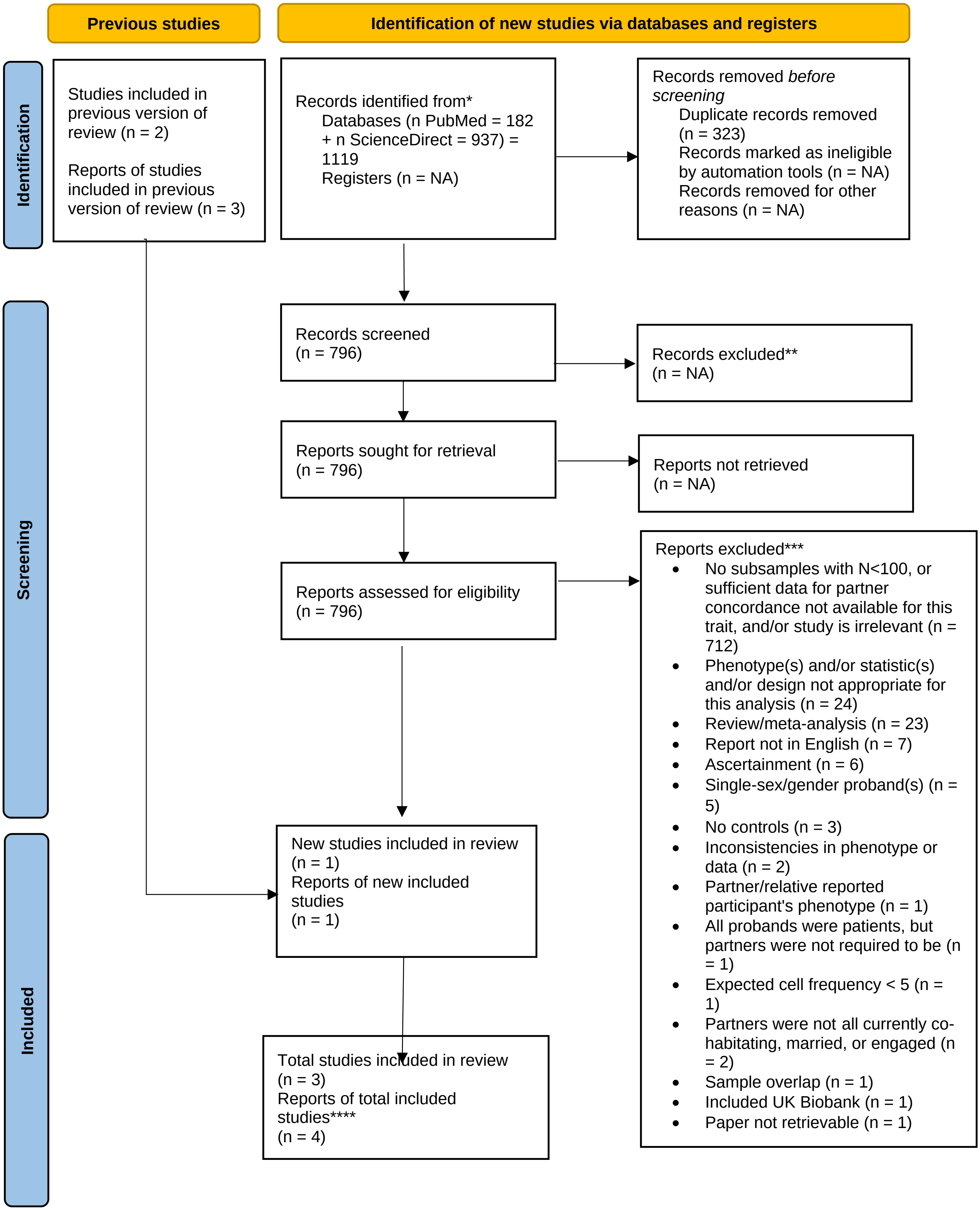

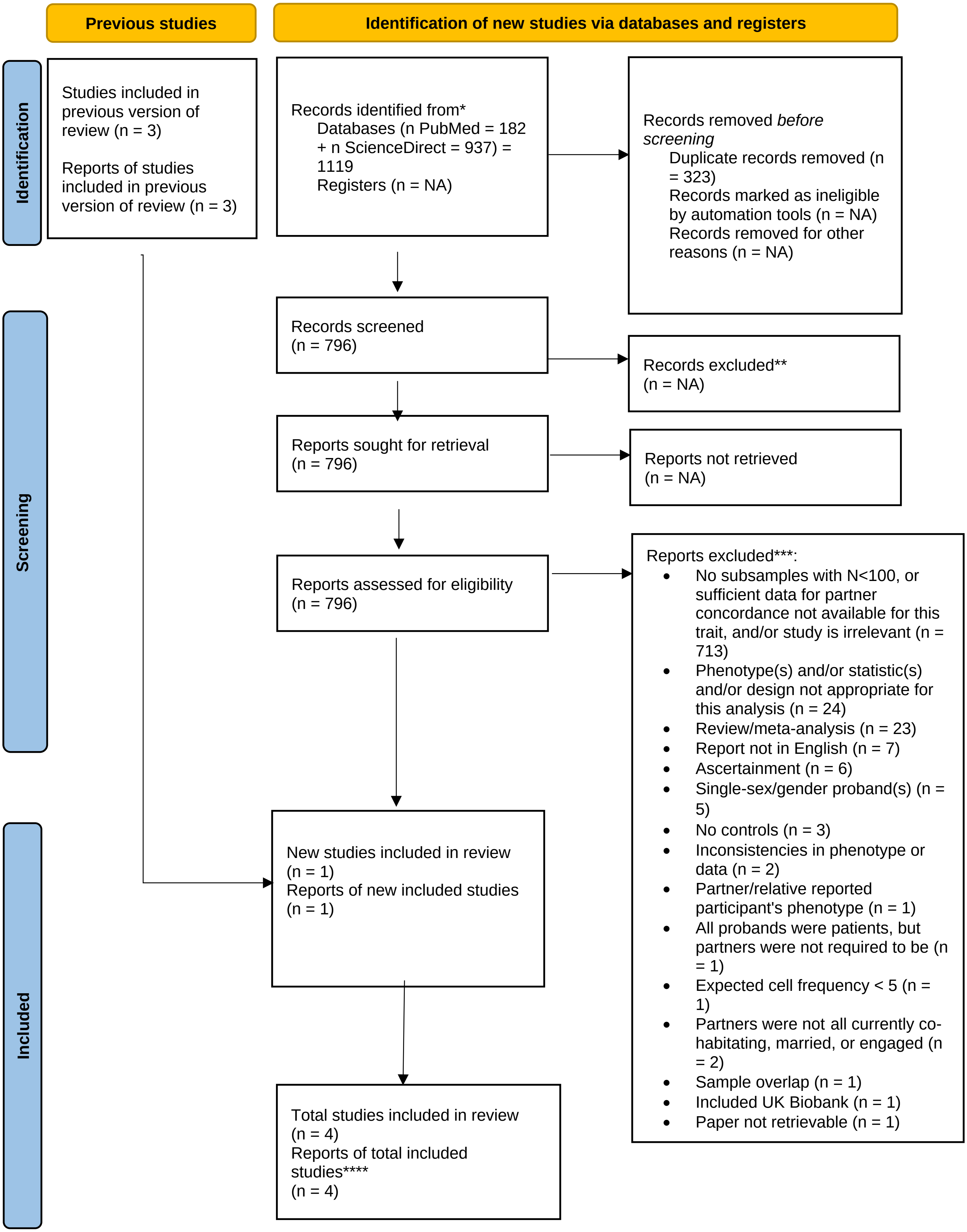
Previous studies
Identification of new studies via databases and registers
Identification
Studies included in previous version of review (n = 3)
Reports of studies included in previous version of review (n = 3)
Records identified from*
Databases (n PubMed = 182 + n ScienceDirect = 937) = 1119
Registers (n = NA)
Records removed before screening
Duplicate records removed (n = 323)
Records marked as ineligible by automation tools (n = NA)
Records removed for other reasons (n = NA)
Screening
Records screened (n = 796)
Records excluded** (n = NA)
Reports sought for retrieval (n = 796)
Reports not retrieved (n = NA)
Reports assessed for eligibility (n = 796)
Reports excluded***:
No subsamples with N<100, or sufficient data for partner concordance not available for this trait, and/or study is irrelevant (n = 713)
Phenotype(s) and/or statistic(s) and/or design not appropriate for this analysis (n = 24)
Review/meta-analysis (n = 23)
Report not in English (n = 7)
Ascertainment (n = 6)
Single-sex/gender proband(s) (n = 5)
No controls (n = 3)
Inconsistencies in phenotype or data (n = 2)
Partner/relative reported participant's phenotype (n = 1)
All probands were patients, but partners were not required to be (n = 1)
Expected cell frequency < 5 (n = 1)
Partners were not all currently co-habitating, married, or engaged (n = 2)
Sample overlap (n = 1)
Included UK Biobank (n = 1)
Paper not retrievable (n = 1)
Included
New studies included in review (n = 1)
Reports of new included studies (n = 1)
Total studies included in review (n = 4)
Reports of total included studies**** (n = 4)

**Drinking quantity**

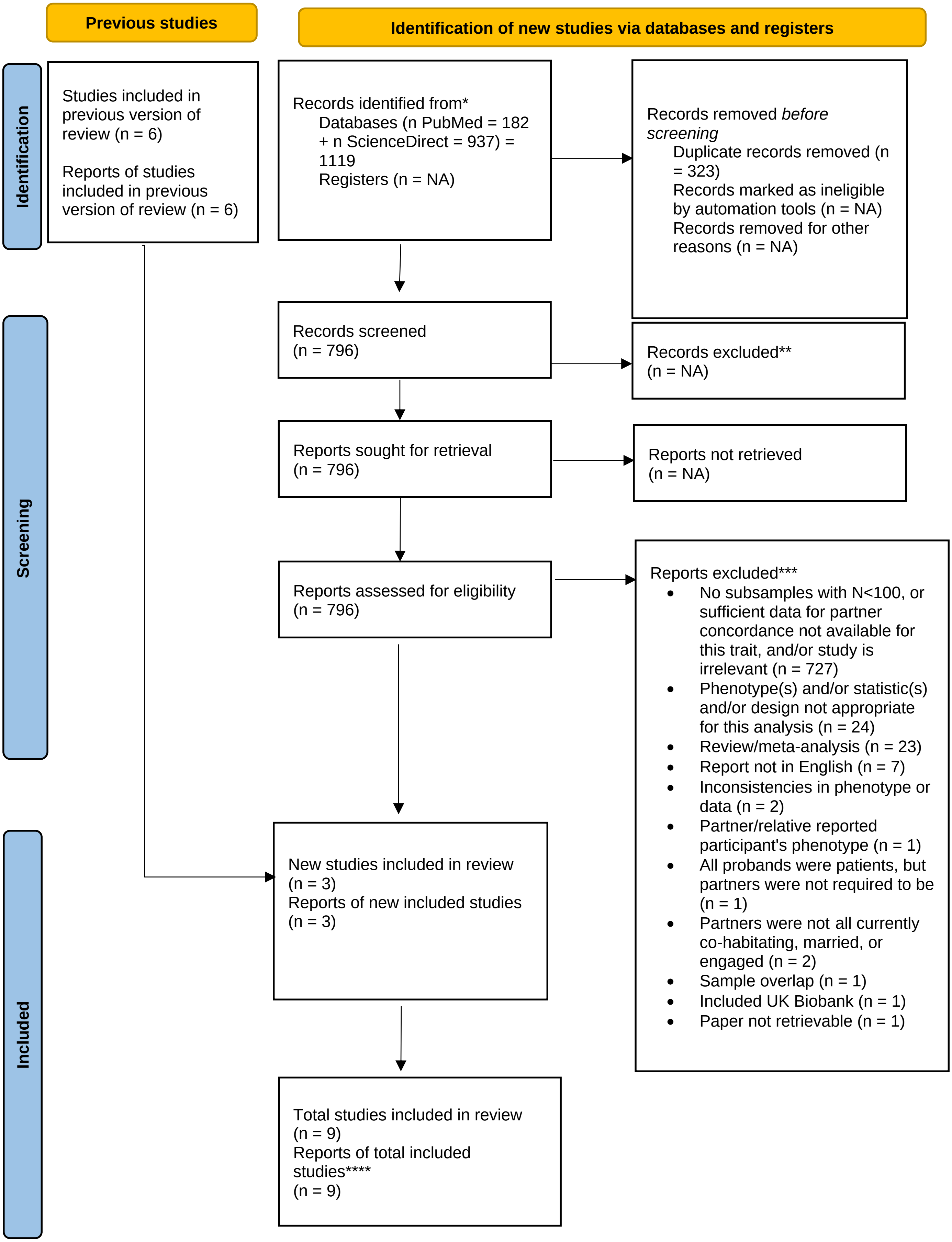

#### Waist-to-hip ratio

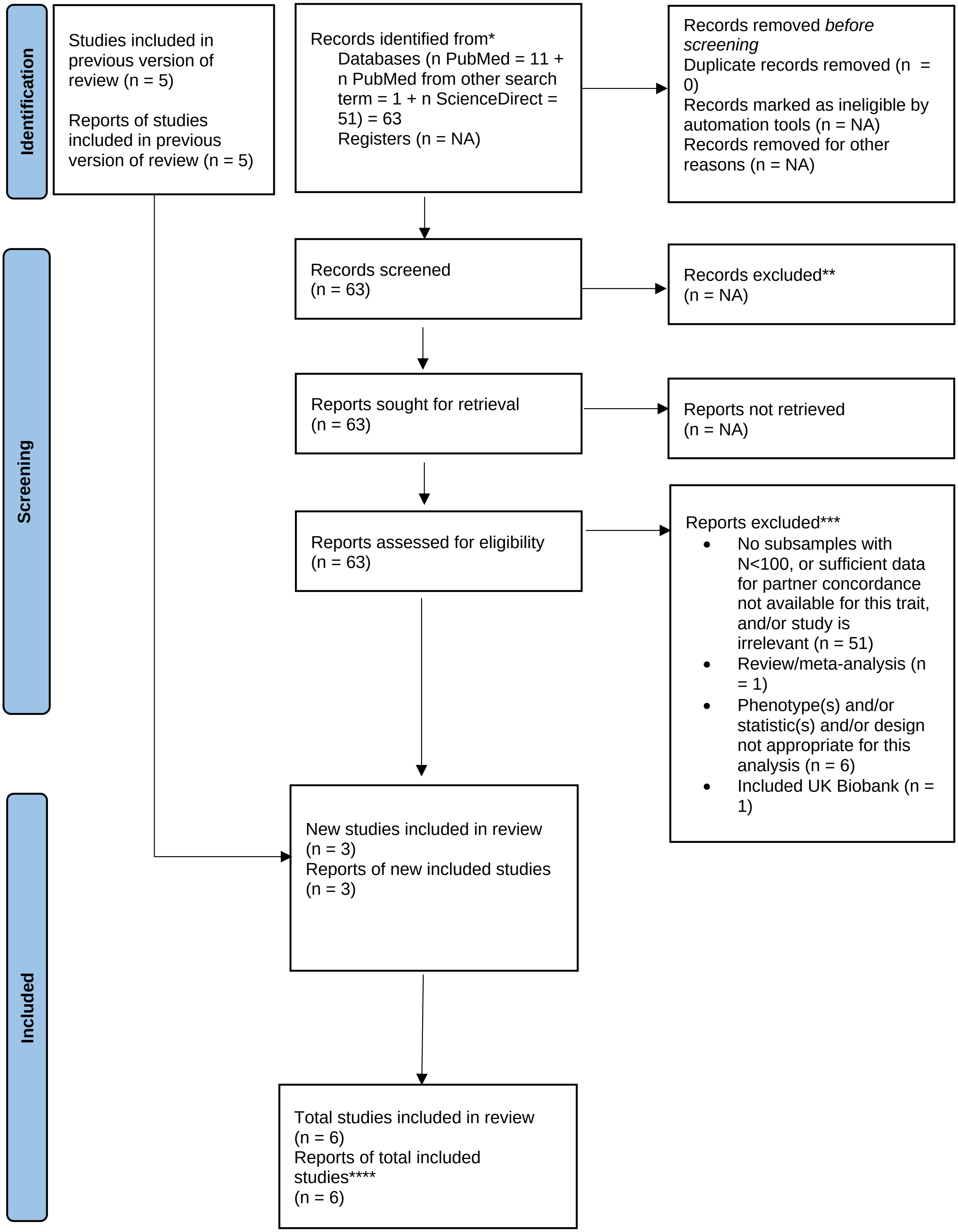

#### Height

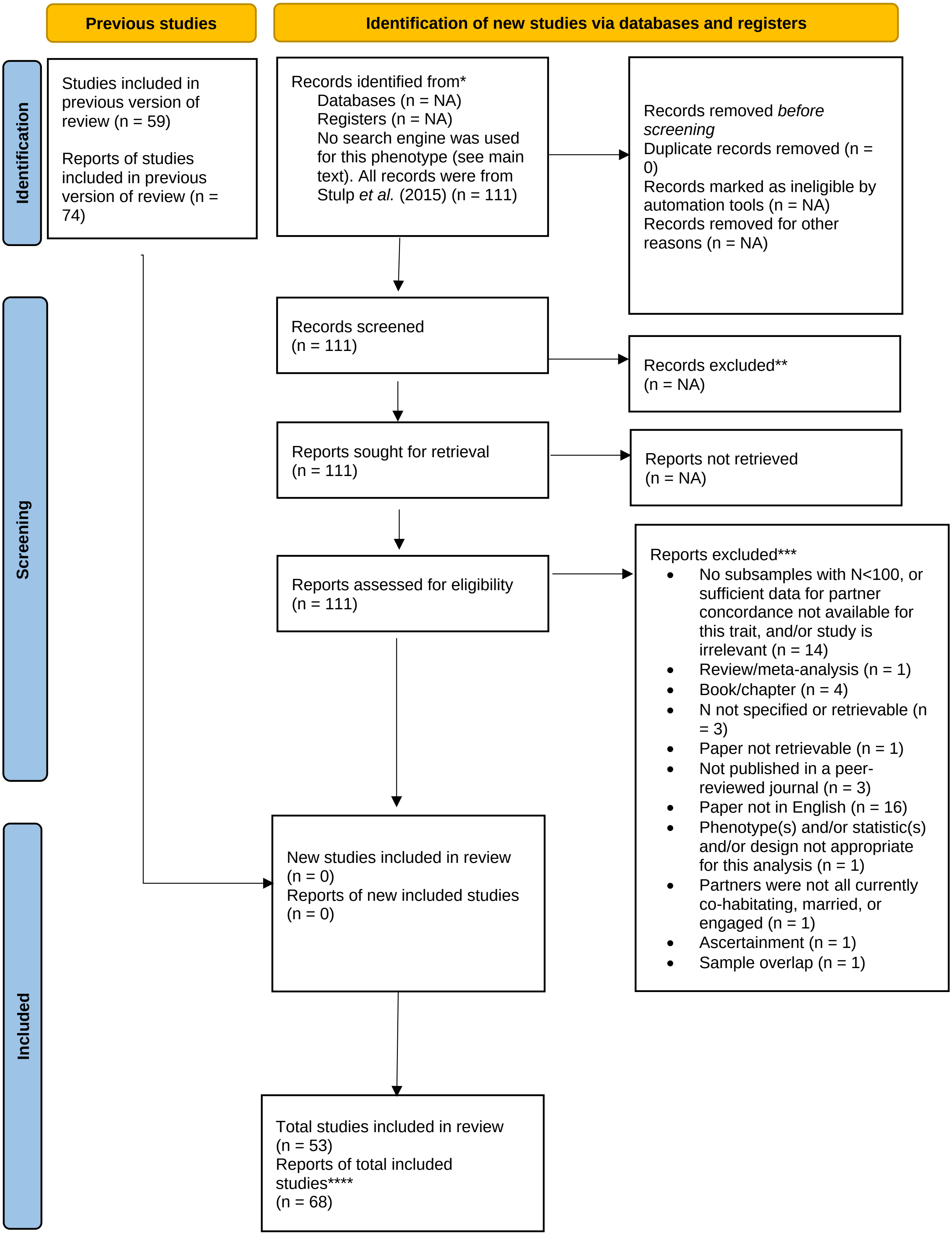
Previous studies
Identification of new studies via databases and registers
Identification
Screening
Included
Studies included in previous version of review (n = 59)
Reports of studies included in previous version of review (n = 74)
Records identified from*
Databases (n = NA)
Registers (n = NA)
No search engine was used for this phenotype (see main text). All records were from Stulp et al. (2015) (n = 111)
Records removed before screening
Duplicate records removed (n = 0)
Records marked as ineligible by automation tools (n = NA)
Records removed for other reasons (n = NA)
Records screened (n = 111)
Records excluded** (n = NA)
Reports sought for retrieval (n = 111)
Reports not retrieved (n = NA)
Reports assessed for eligibility (n = 111)
Reports excluded***
No subsamples with N<100, or sufficient data for partner concordance not available for this trait, and/or study is irrelevant (n = 14)
Review/meta-analysis (n = 1)
Book/chapter (n = 4)
N not specified or retrievable (n = 3)
Paper not retrievable (n = 1)
Not published in a peer-reviewed journal (n = 3)
Paper not in English (n = 16)
Phenotype(s) and/or statistic(s) and/or design not appropriate for this analysis (n = 1)
Partners were not all currently co-habitating, married, or engaged (n = 1)
Ascertainment (n = 1)
Sample overlap (n = 1)
New studies included in review (n = 0)
Reports of new included studies (n = 0)
Total studies included in review (n = 53)
Reports of total included studies**** (n = 68)

**Body mass index**

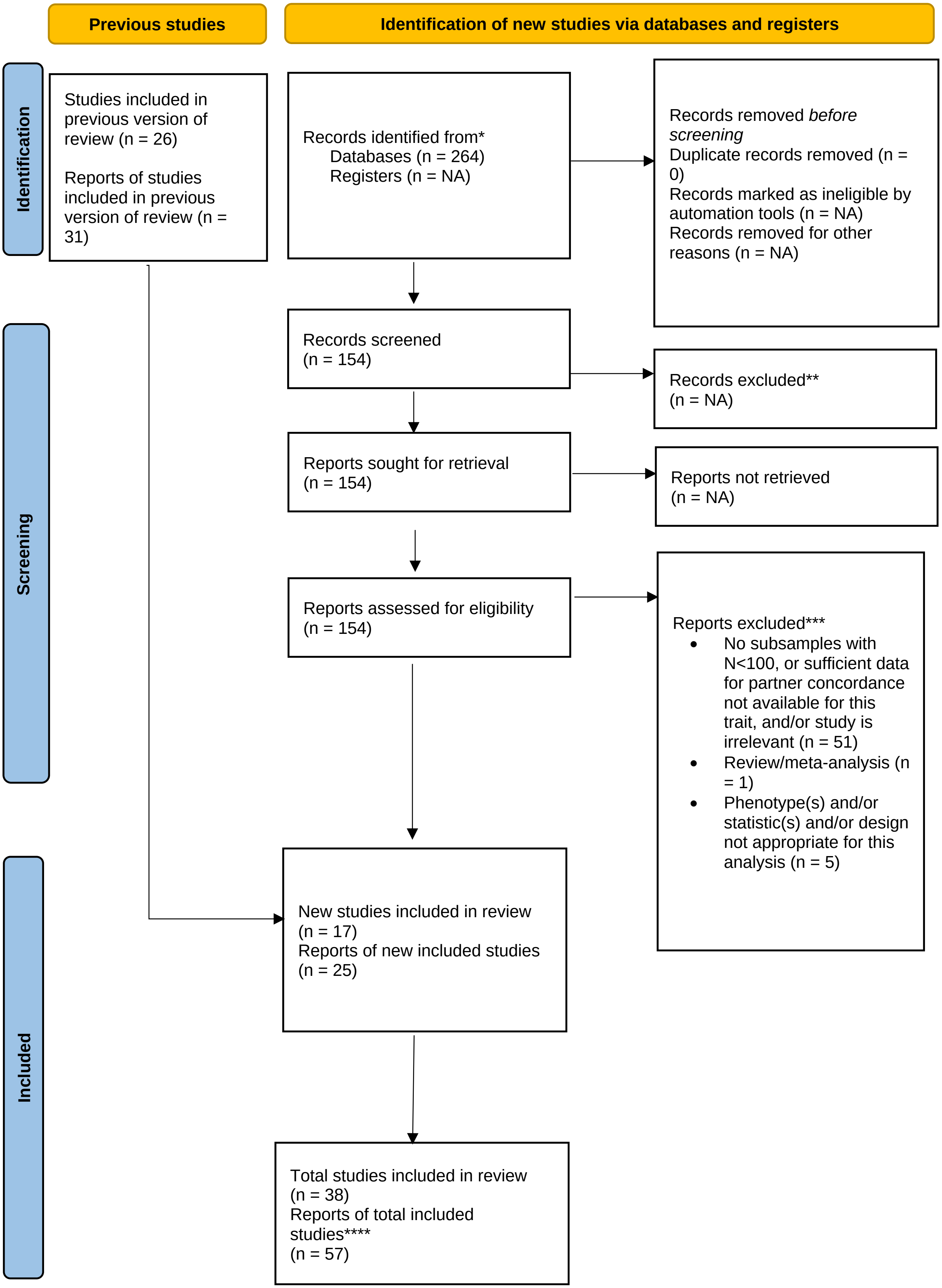

**Extraversion**

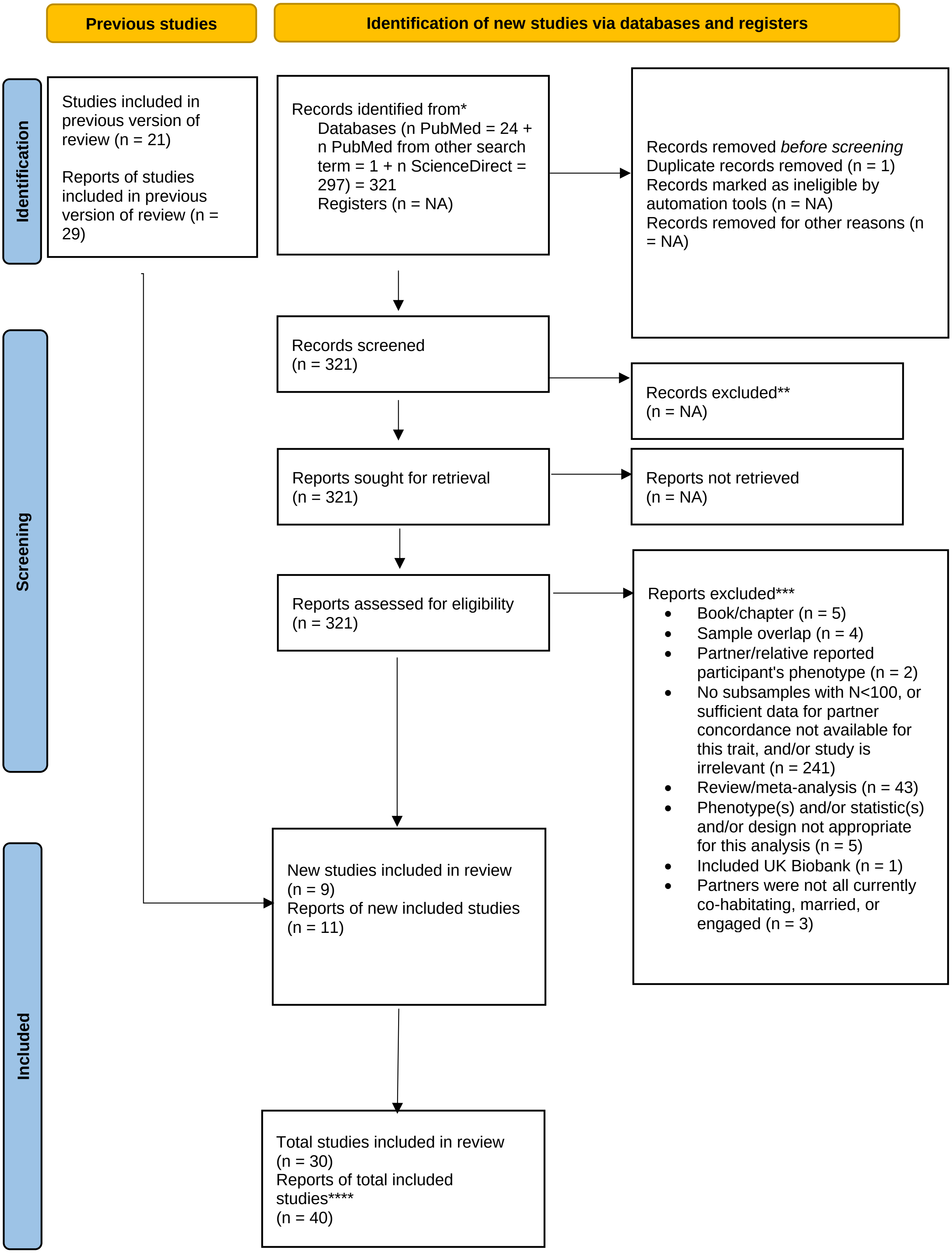

#### Neuroticism

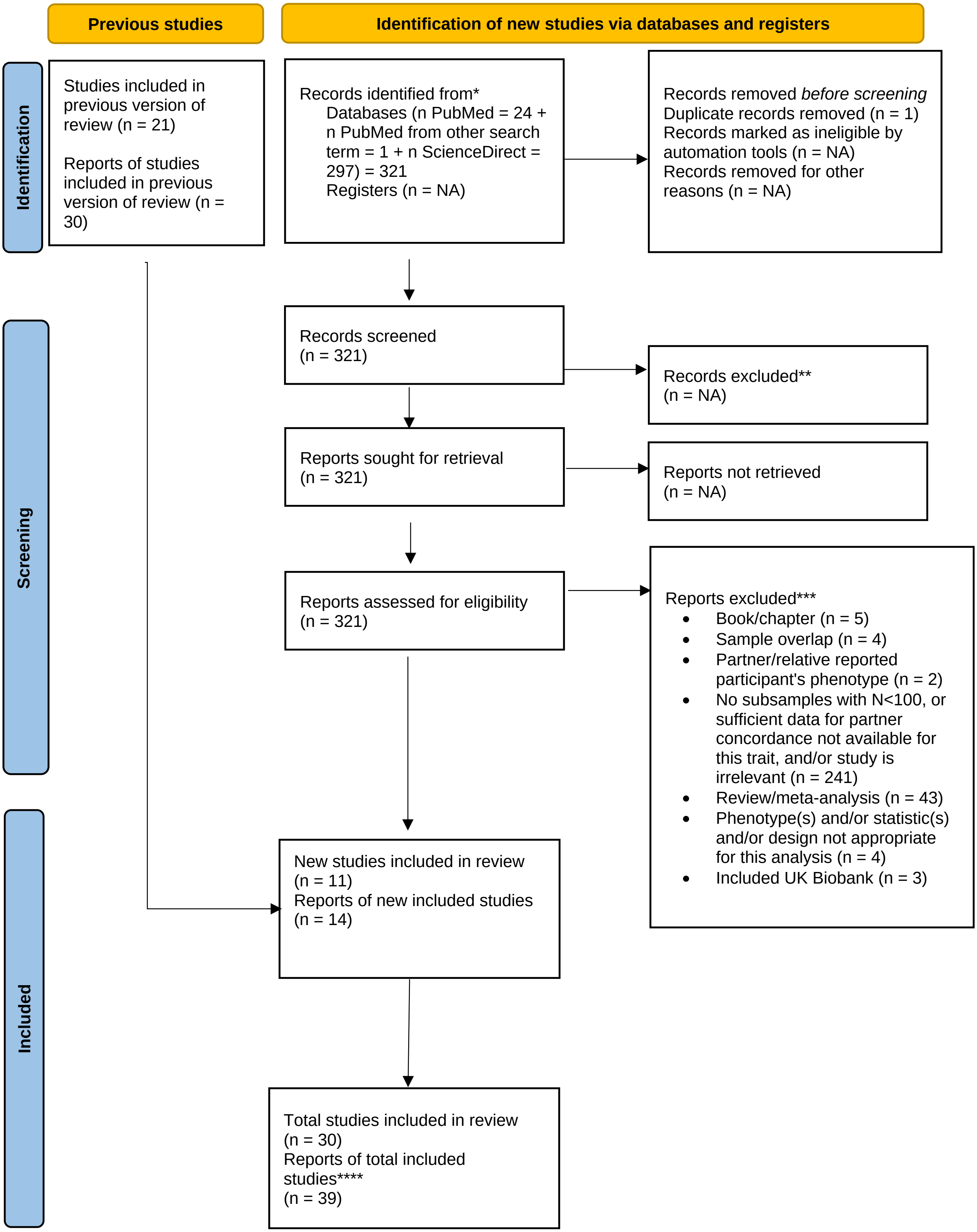

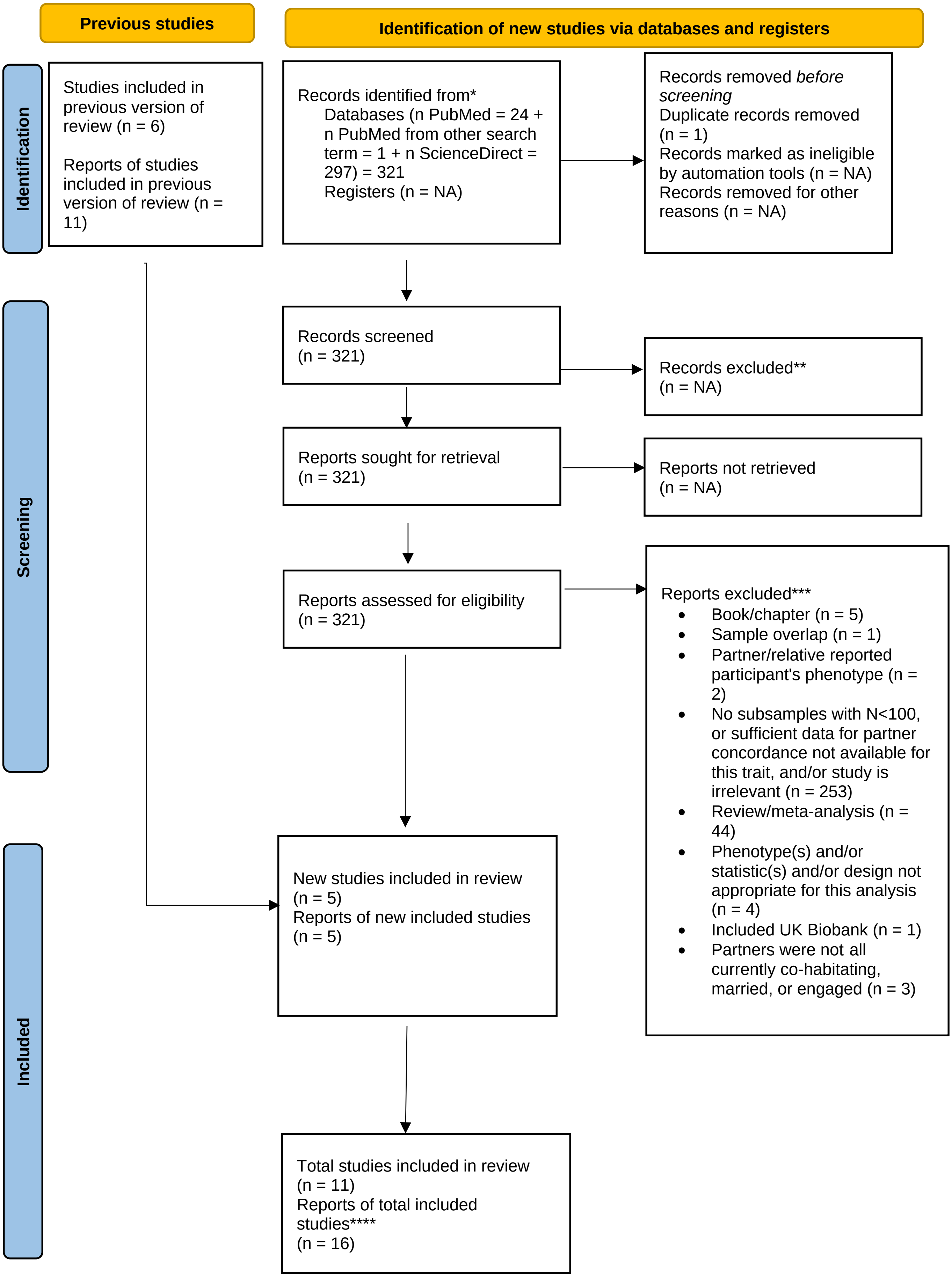
Previous studies
Identification of new studies via databases and registers
Identification
Screening
Included
Studies included in previous version of review (n = 6)
Reports of studies included in previous version of review (n = 11)
Records identified from*
Databases (n PubMed = 24 + n PubMed from other search term = 1 + n ScienceDirect = 297) = 321
Registers (n = NA)
Records removed *before screening*
Duplicate records removed (n = 1)
Records marked as ineligible by automation tools (n = NA)
Records removed for other reasons (n = NA)
Records screened (n = 321)
Records excluded** (n = NA)
Reports sought for retrieval (n = 321)
Reports not retrieved (n = NA)
Reports assessed for eligibility (n = 321)
Reports excluded***
• Book/chapter (n = 5)
• Sample overlap (n = 1)
• Partner/relative reported participant's phenotype (n = 2)
• No subsamples with N<100, or sufficient data for partner concordance not available for this trait, and/or study is irrelevant (n = 253)
• Review/meta-analysis (n = 44)
• Phenotype(s) and/or statistic(s) and/or design not appropriate for this analysis (n = 4)
• Included UK Biobank (n = 1)
• Partners were not all currently co-habitating, married, or engaged (n = 3)
New studies included in review (n = 5)
Reports of new included studies (n = 5)
Total studies included in review (n = 11)
Reports of total included studies**** (n = 16)

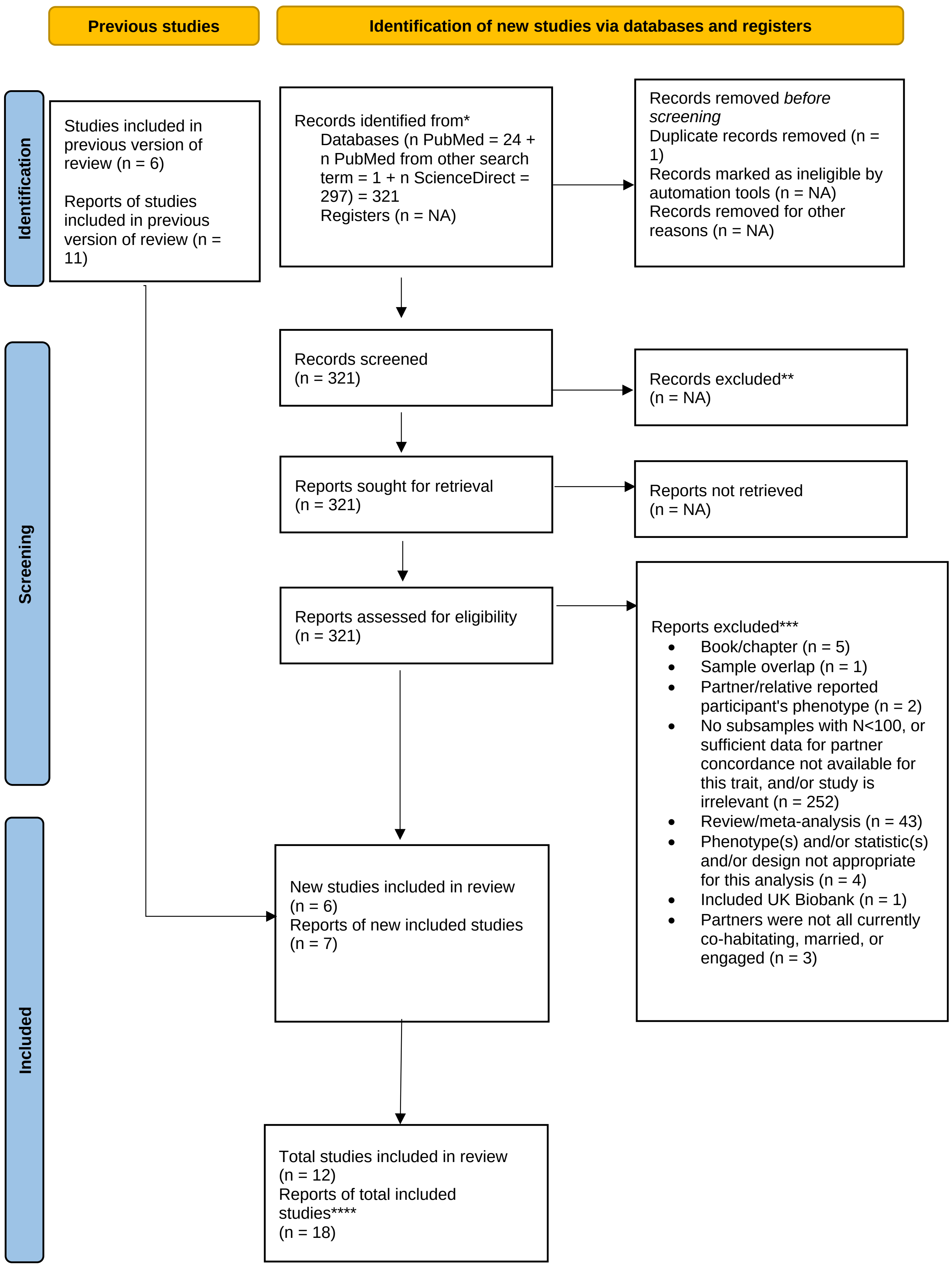
Previous studies
Identification of new studies via databases and registers
Identification
Studies included in previous version of review (n = 6)
Reports of studies included in previous version of review (n = 11)
Records identified from*
Databases (n PubMed = 24 + n PubMed from other search term = 1 + n ScienceDirect = 297) = 321
Registers (n = NA)
Records removed *before screening*
Duplicate records removed (n = 1)
Records marked as ineligible by automation tools (n = NA)
Records removed for other reasons (n = NA)
Screening
Records screened (n = 321)
Records excluded** (n = NA)
Reports sought for retrieval (n = 321)
Reports not retrieved (n = NA)
Reports assessed for eligibility (n = 321)
Reports excluded***
Book/chapter (n = 5)
Sample overlap (n = 1)
Partner/relative reported participant's phenotype (n = 2)
No subsamples with N<100, or sufficient data for partner concordance not available for this trait, and/or study is irrelevant (n = 252)
Review/meta-analysis (n = 43)
Phenotype(s) and/or statistic(s) and/or design not appropriate for this analysis (n = 4)
Included UK Biobank (n = 1)
Partners were not all currently co-habitating, married, or engaged (n = 3)
Included
New studies included in review (n = 6)
Reports of new included studies (n = 7)
Total studies included in review (n = 12)
Reports of total included studies**** (n = 18)

#### Agreeableness

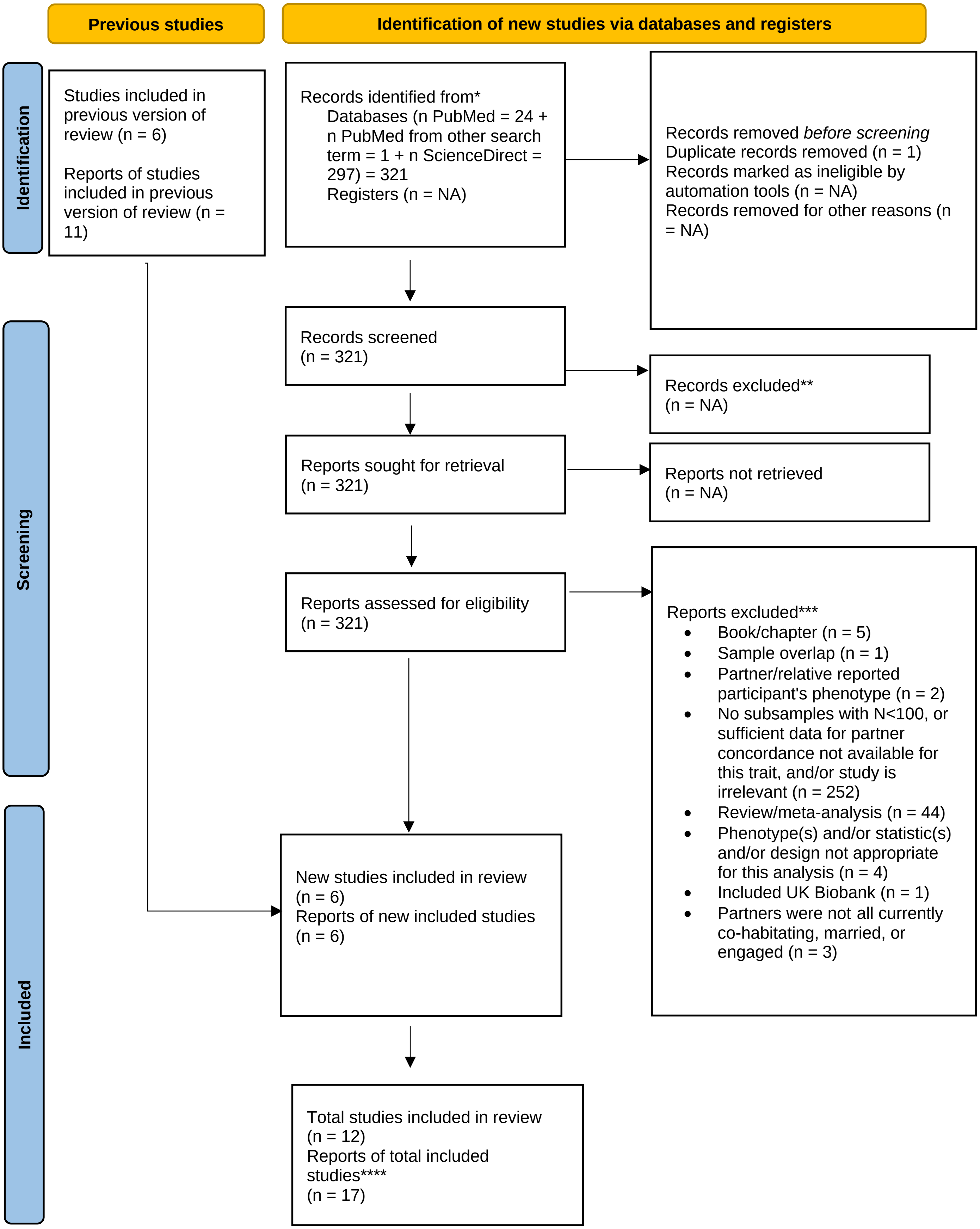

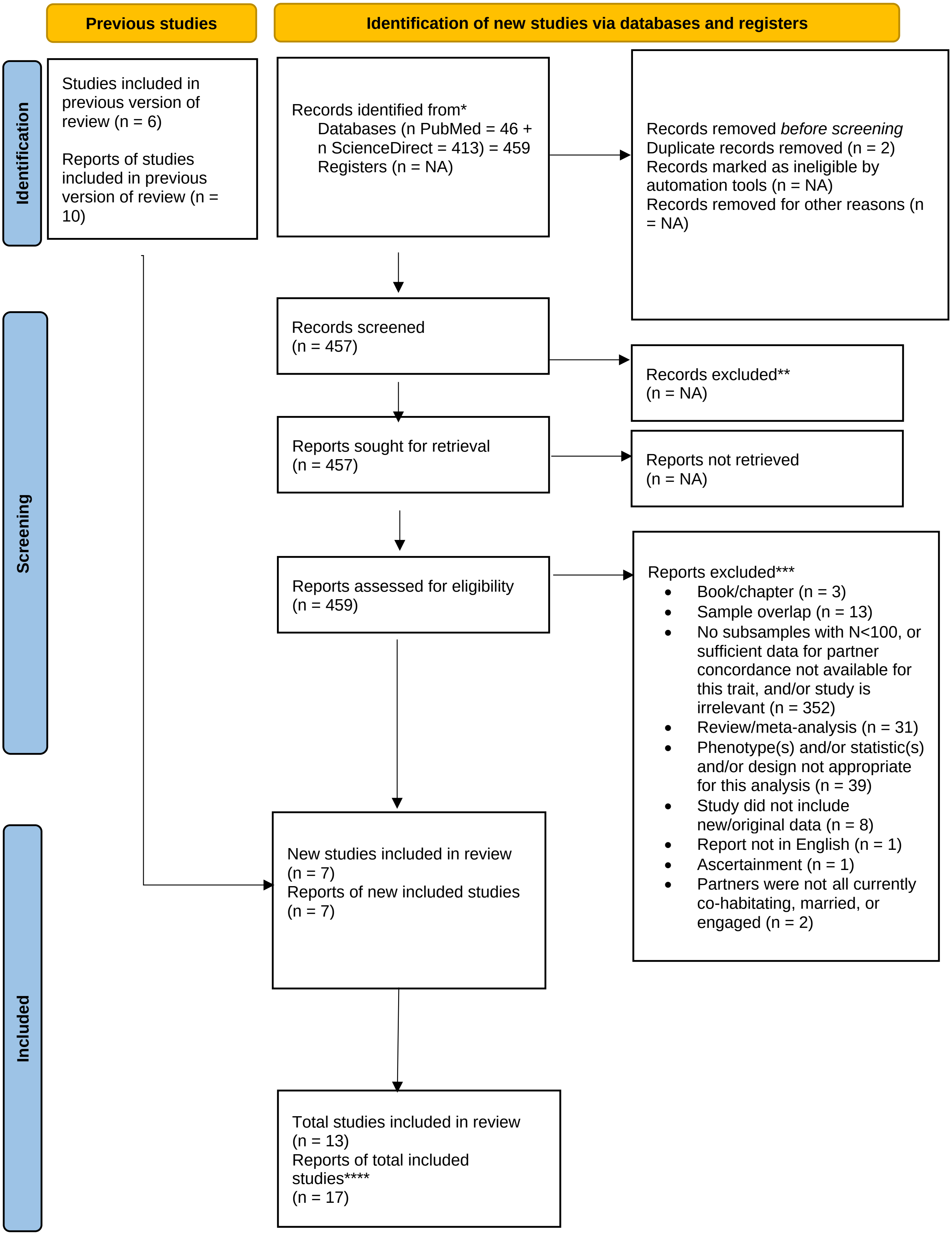
Previous studies
Identification of new studies via databases and registers
Identification
Studies included in previous version of review (n = 6)
Reports of studies included in previous version of review (n = 10)
Records identified from*
Databases (n PubMed = 46 + n ScienceDirect = 413) = 459
Registers (n = NA)
Records removed *before screening*
Duplicate records removed (n = 2)
Records marked as ineligible by automation tools (n = NA)
Records removed for other reasons (n = NA)
Screening
Records screened
(n = 457)
Records excluded**
(n = NA)
Reports sought for retrieval
(n = 457)
Reports not retrieved
(n = NA)
Reports assessed for eligibility
(n = 459)
Reports excluded***
• Book/chapter (n = 3)
• Sample overlap (n = 13)
• No subsamples with N<100, or sufficient data for partner concordance not available for this trait, and/or study is irrelevant (n = 352)
• Review/meta-analysis (n = 31)
• Phenotype(s) and/or statistic(s) and/or design not appropriate for this analysis (n = 39)
• Study did not include new/original data (n = 8)
• Report not in English (n = 1)
• Ascertainment (n = 1)
• Partners were not all currently co-habitating, married, or engaged (n = 2)
Included
New studies included in review
(n = 7)
Reports of new included studies
(n = 7)
Total studies included in review
(n = 13)
Reports of total included studies****
(n = 17)

#### Diabetes

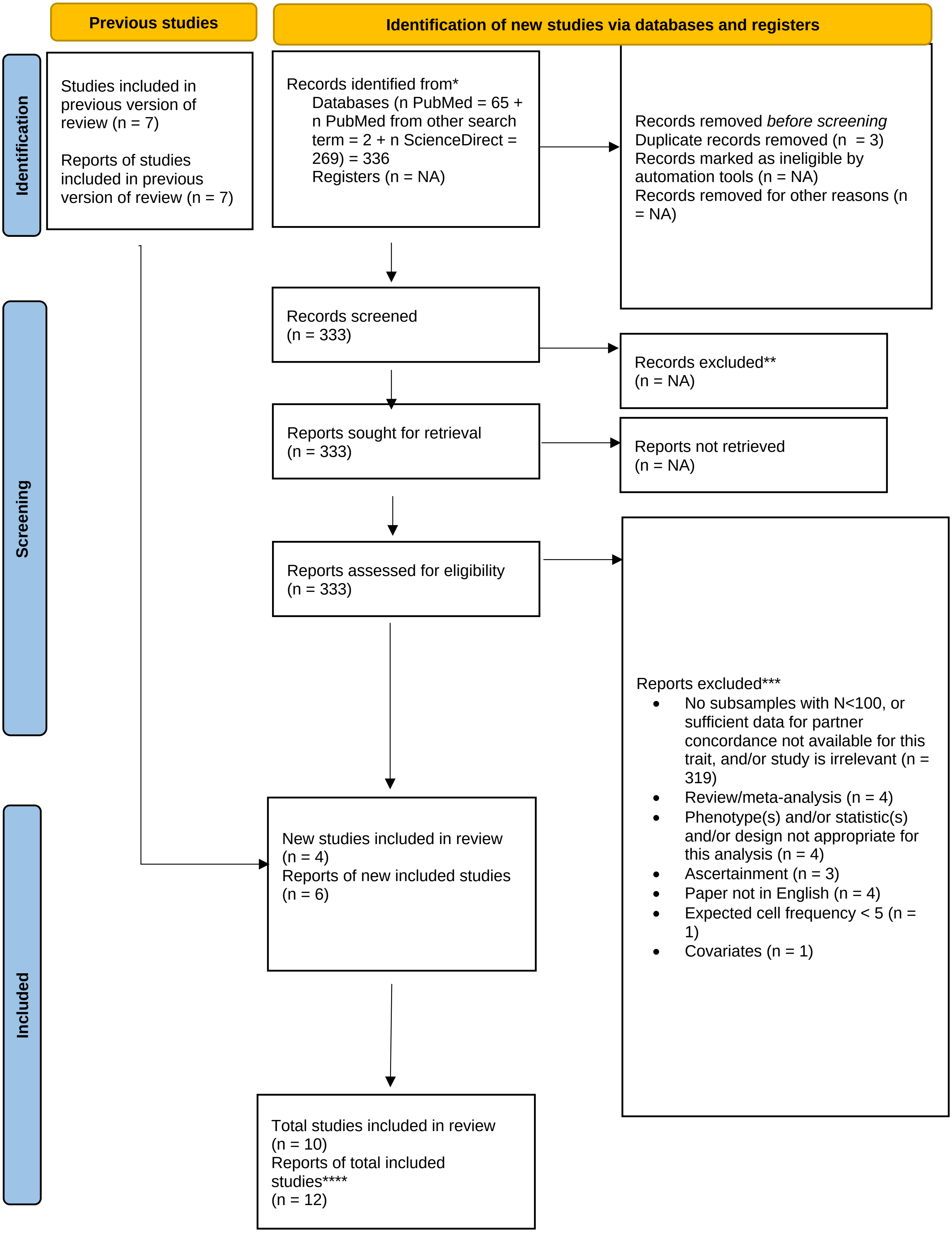
Previous studies
Identification of new studies via databases and registers
Identification
Studies included in previous version of review (n = 7)
Reports of studies included in previous version of review (n = 7)
Records identified from*
Databases (n PubMed = 65 + n PubMed from other search term = 2 + n ScienceDirect = 269) = 336
Registers (n = NA)
Records removed *before screening*
Duplicate records removed (n = 3)
Records marked as ineligible by automation tools (n = NA)
Records removed for other reasons (n = NA)
Screening
Records screened
(n = 333)
Records excluded**
(n = NA)
Reports sought for retrieval
(n = 333)
Reports not retrieved
(n = NA)
Reports assessed for eligibility
(n = 333)
Reports excluded***
No subsamples with N<100, or sufficient data for partner concordance not available for this trait, and/or study is irrelevant (n = 319)
Review/meta-analysis (n = 4)
Phenotype(s) and/or statistic(s) and/or design not appropriate for this analysis (n = 4)
Ascertainment (n = 3)
Paper not in English (n = 4)
Expected cell frequency < 5 (n = 1)
Covariates (n = 1)
Included
New studies included in review
(n = 4)
Reports of new included studies
(n = 6)
Total studies included in review
(n = 10)
Reports of total included studies****
(n = 12)

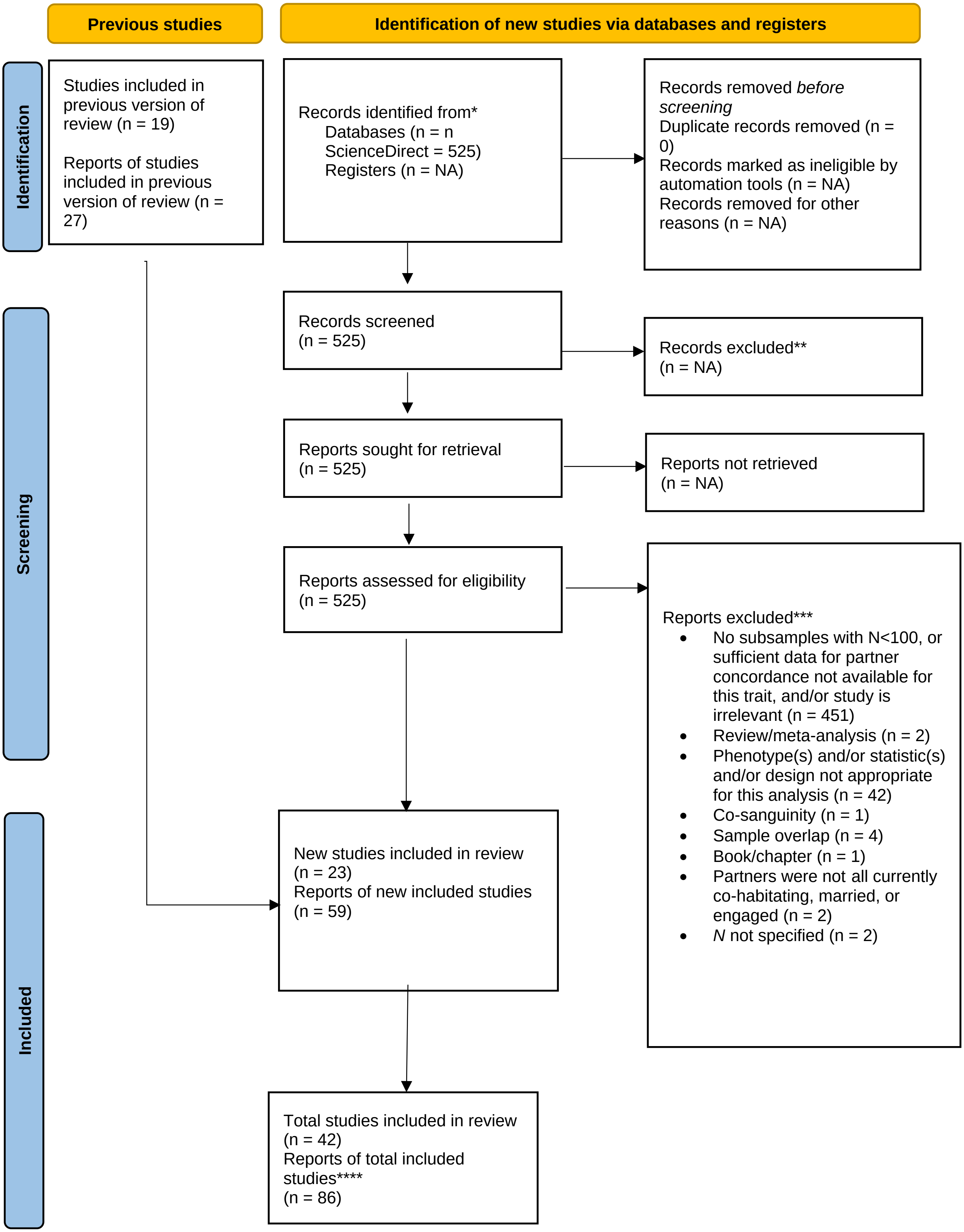
Previous studies
Identification of new studies via databases and registers
Identification
Screening
Included
Studies included in previous version of review (n = 19)
Reports of studies included in previous version of review (n = 27)
Records identified from*
Databases (n = n ScienceDirect = 525)
Registers (n = NA)
Records removed before screening
Duplicate records removed (n = 0)
Records marked as ineligible by automation tools (n = NA)
Records removed for other reasons (n = NA)
Records screened (n = 525)
Records excluded** (n = NA)
Reports sought for retrieval (n = 525)
Reports not retrieved (n = NA)
Reports assessed for eligibility (n = 525)
Reports excluded***
No subsamples with N<100, or sufficient data for partner concordance not available for this trait, and/or study is irrelevant (n = 451)
Review/meta-analysis (n = 2)
Phenotype(s) and/or statistic(s) and/or design not appropriate for this analysis (n = 42)
Co-sanguinity (n = 1)
Sample overlap (n = 4)
Book/chapter (n = 1)
Partners were not all currently co-habitating, married, or engaged (n = 2)
N not specified (n = 2)
New studies included in review (n = 23)
Reports of new included studies (n = 59)
Total studies included in review (n = 42)
Reports of total included studies**** (n = 86)

#### Smoking status

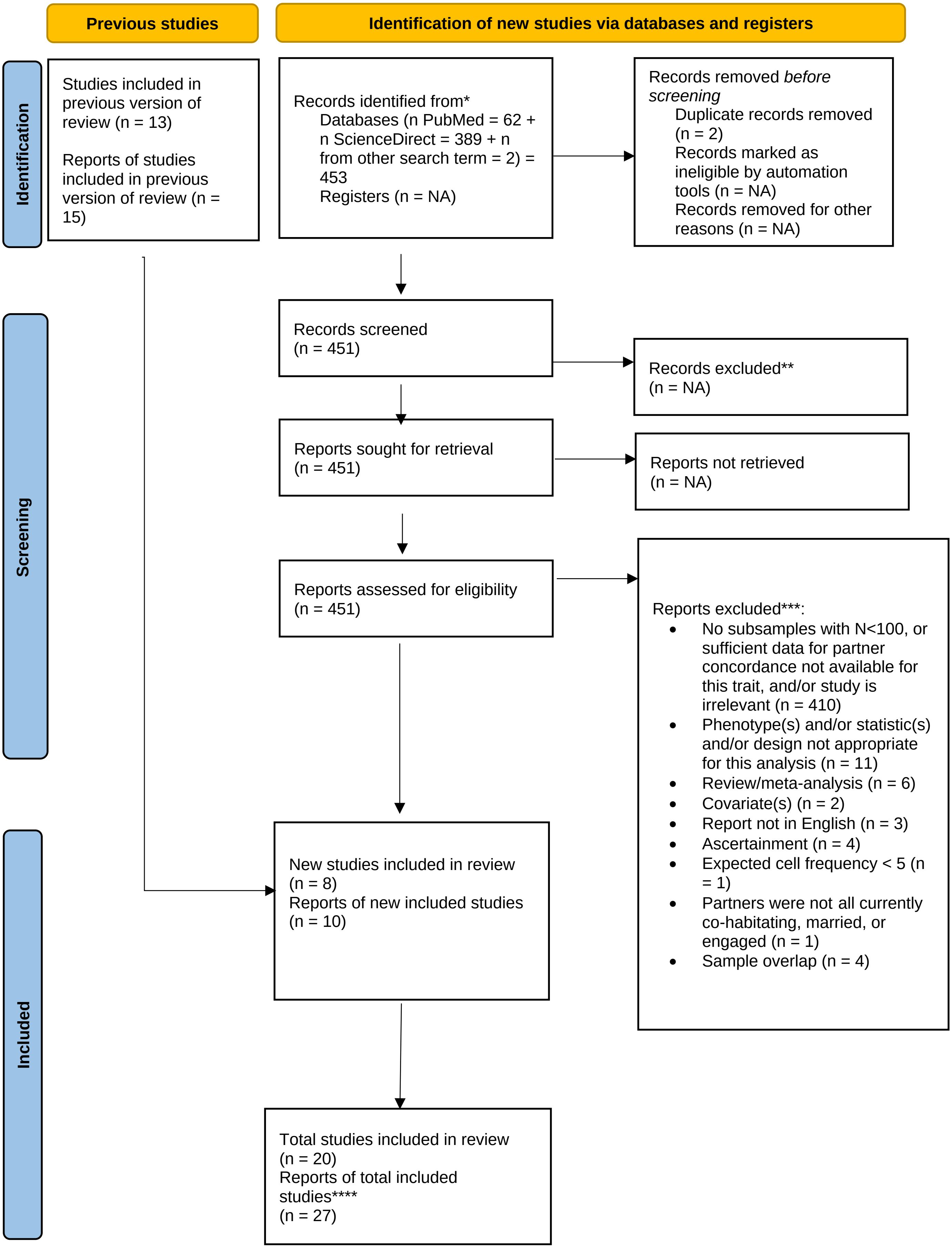

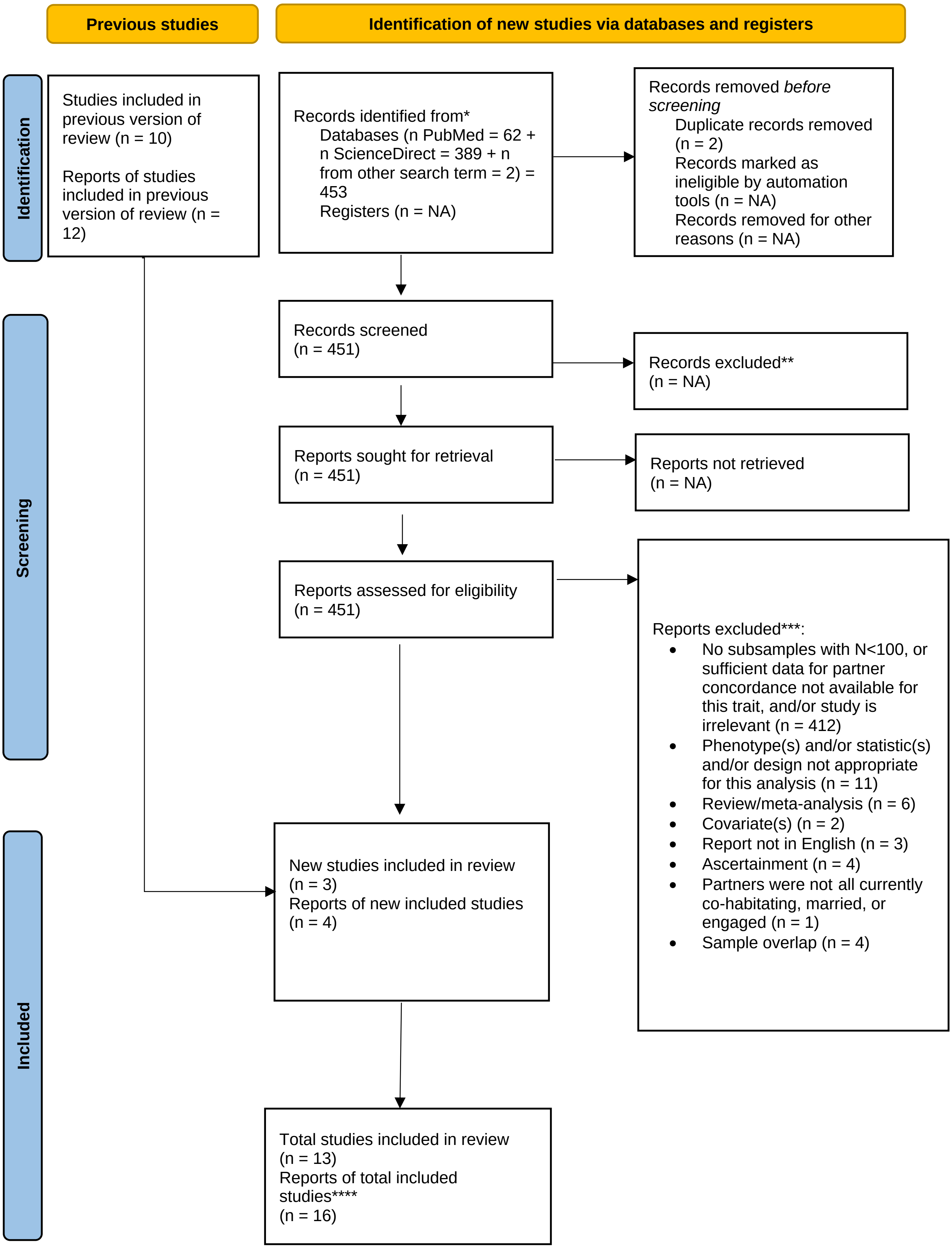
Previous studies
Identification of new studies via databases and registers
Identification
Screening
Included
Studies included in previous version of review (n = 10)
Reports of studies included in previous version of review (n = 12)
Records identified from*
Databases (n PubMed = 62 + n ScienceDirect = 389 + n from other search term = 2) = 453
Registers (n = NA)
Records removed *before screening*
Duplicate records removed (n = 2)
Records marked as ineligible by automation tools (n = NA)
Records removed for other reasons (n = NA)
Records screened (n = 451)
Records excluded** (n = NA)
Reports sought for retrieval (n = 451)
Reports not retrieved (n = NA)
Reports assessed for eligibility (n = 451)
Reports excluded***:
• No subsamples with N<100, or sufficient data for partner concordance not available for this trait, and/or study is irrelevant (n = 412)
• Phenotype(s) and/or statistic(s) and/or design not appropriate for this analysis (n = 11)
• Review/meta-analysis (n = 6)
• Covariate(s) (n = 2)
• Report not in English (n = 3)
• Ascertainment (n = 4)
• Partners were not all currently co-habitating, married, or engaged (n = 1)
• Sample overlap (n = 4)
New studies included in review (n = 3)
Reports of new included studies (n = 4)
Total studies included in review (n = 13)
Reports of total included studies**** (n = 16)

**Smoking cessation**

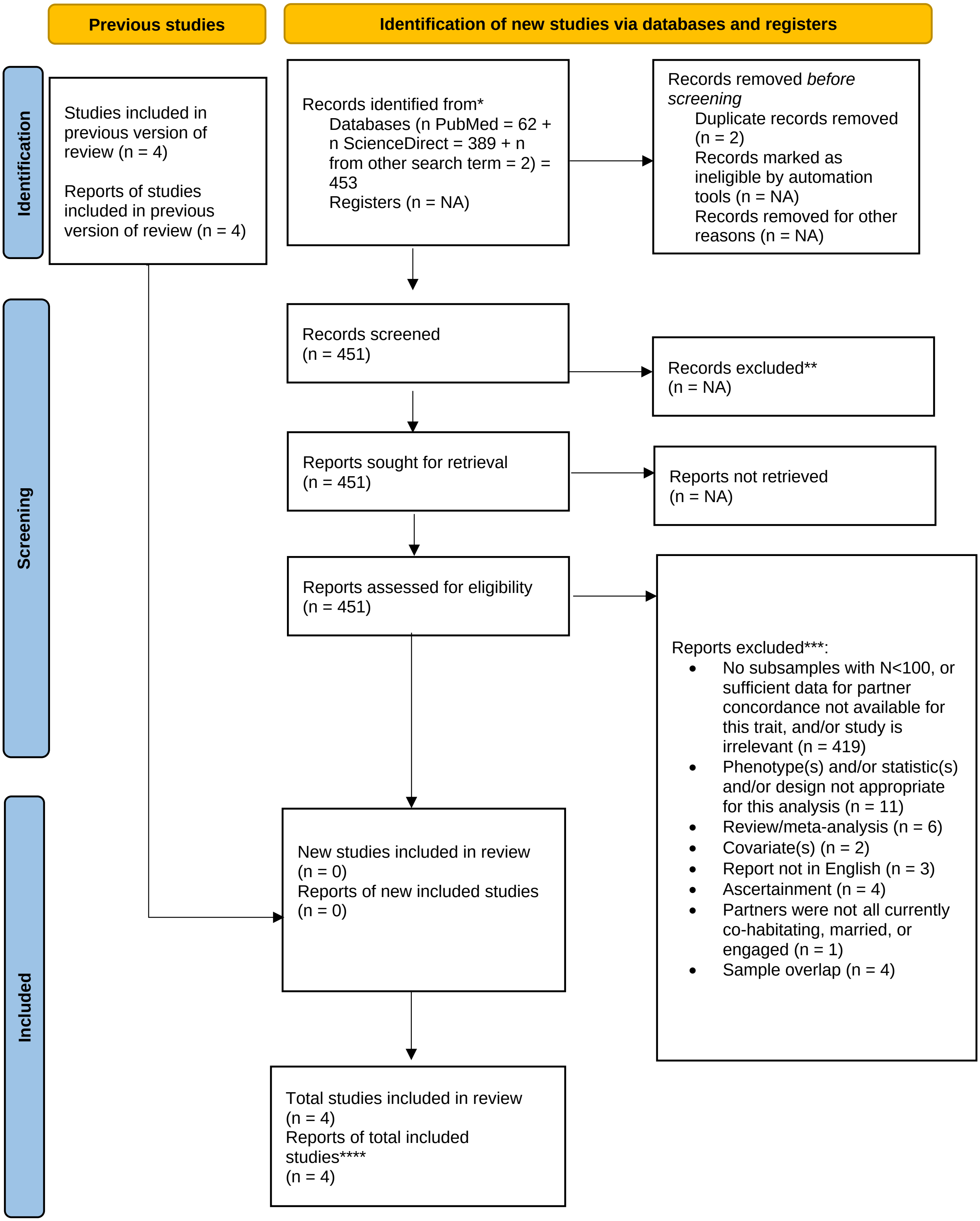

**Smoking quantity**

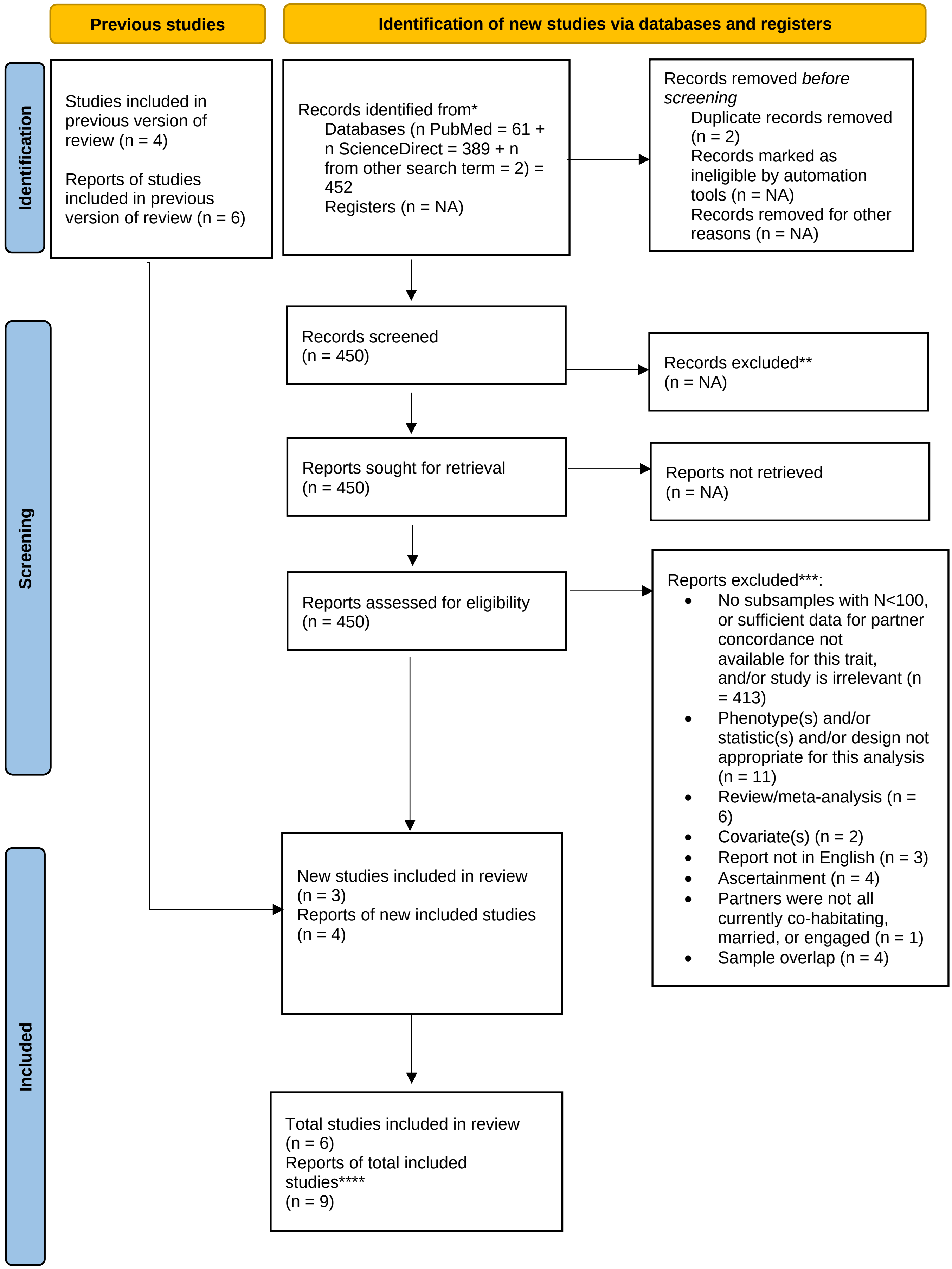

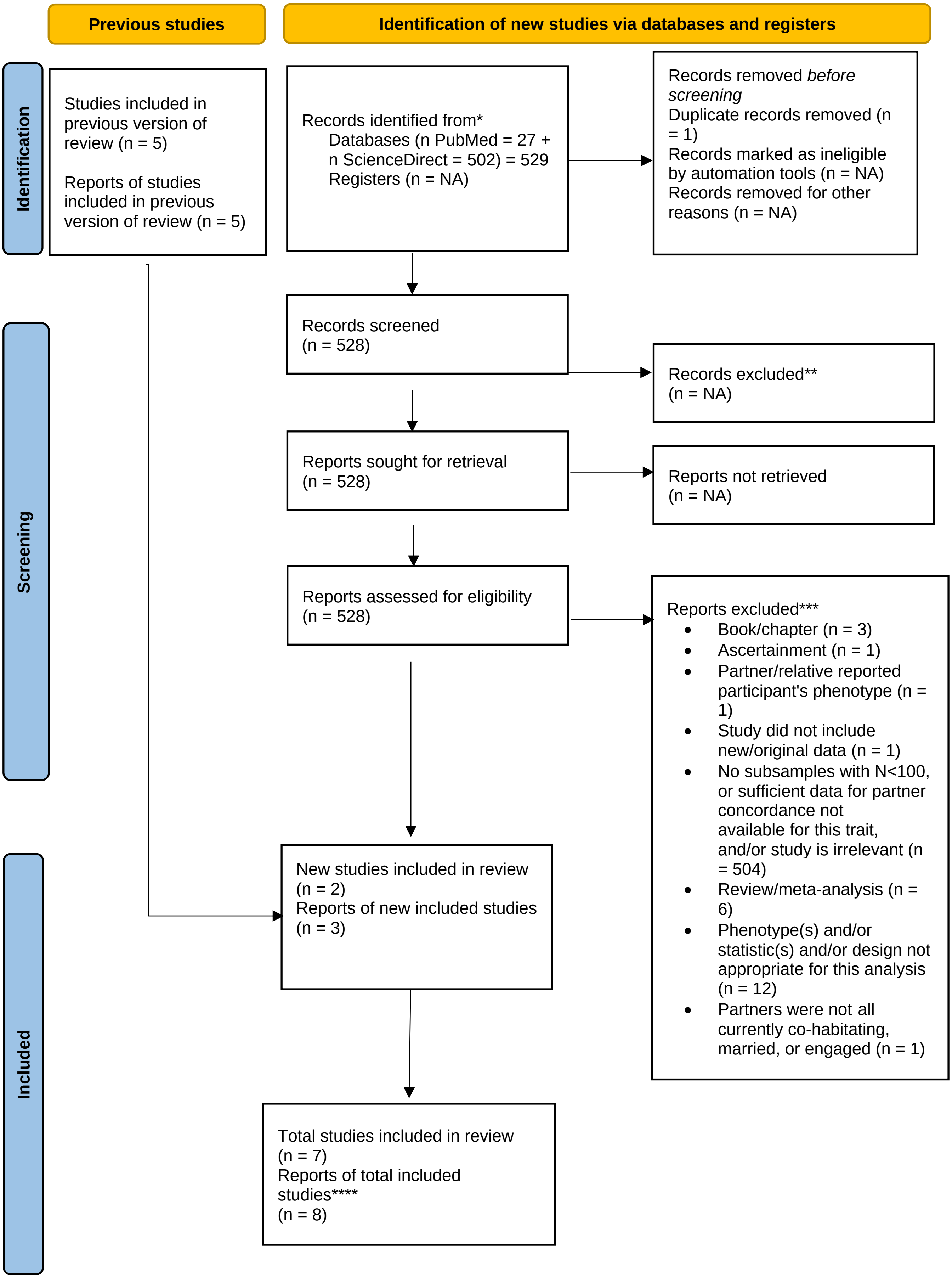
Previous studies
Identification of new studies via databases and registers
Identification
Screening
Included
Studies included in previous version of review (n = 5)
Reports of studies included in previous version of review (n = 5)
Records identified from*
Databases (n PubMed = 27 + n ScienceDirect = 502) = 529
Registers (n = NA)
Records removed *before screening*
Duplicate records removed (n = 1)
Records marked as ineligible by automation tools (n = NA)
Records removed for other reasons (n = NA)
Records screened
(n = 528)
Records excluded**
(n = NA)
Reports sought for retrieval
(n = 528)
Reports not retrieved
(n = NA)
Reports assessed for eligibility
(n = 528)
Reports excluded***
• Book/chapter (n = 3)
• Ascertainment (n = 1)
• Partner/relative reported participant's phenotype (n = 1)
• Study did not include new/original data (n = 1)
• No subsamples with N<100, or sufficient data for partner concordance not available for this trait, and/or study is irrelevant (n = 504)
• Review/meta-analysis (n = 6)
• Phenotype(s) and/or statistic(s) and/or design not appropriate for this analysis (n = 12)
• Partners were not all currently co-habitating, married, or engaged (n = 1)
New studies included in review
(n = 2)
Reports of new included studies
(n = 3)
Total studies included in review
(n = 7)
Reports of total included studies****
(n = 8)

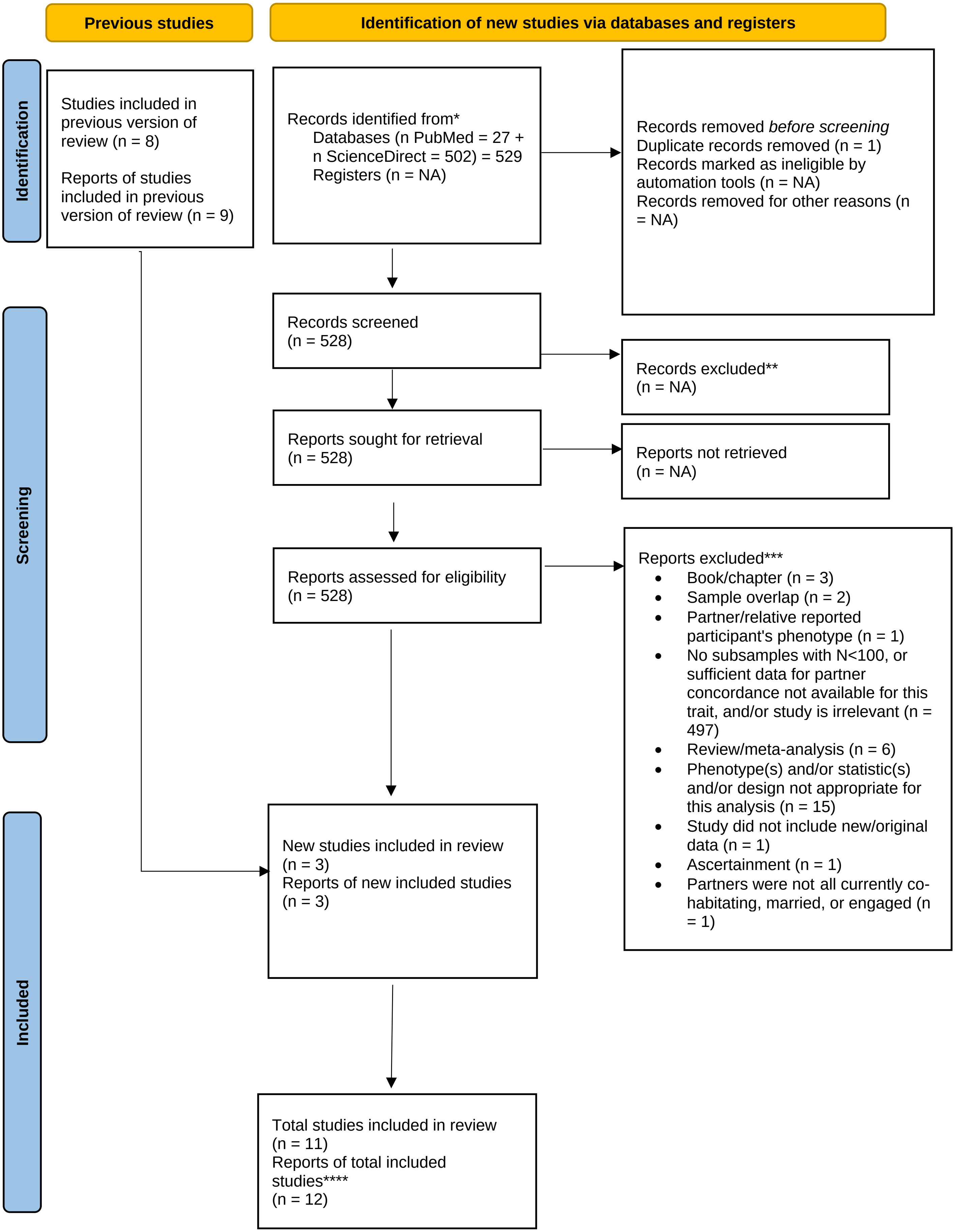
Previous studies
Identification of new studies via databases and registers
Identification
Studies included in previous version of review (n = 8)
Reports of studies included in previous version of review (n = 9)
Records identified from*
Databases (n PubMed = 27 + n ScienceDirect = 502) = 529
Registers (n = NA)
Records removed before screening
Duplicate records removed (n = 1)
Records marked as ineligible by automation tools (n = NA)
Records removed for other reasons (n = NA)
Screening
Records screened
(n = 528)
Records excluded**
(n = NA)
Reports sought for retrieval
(n = 528)
Reports not retrieved
(n = NA)
Reports assessed for eligibility
(n = 528)
Reports excluded***
Book/chapter (n = 3)
Sample overlap (n = 2)
Partner/relative reported participant's phenotype (n = 1)
No subsamples with N<100, or sufficient data for partner concordance not available for this trait, and/or study is irrelevant (n = 497)
Review/meta-analysis (n = 6)
Phenotype(s) and/or statistic(s) and/or design not appropriate for this analysis (n = 15)
Study did not include new/original data (n = 1)
Ascertainment (n = 1)
Partners were not all currently co-habitating, married, or engaged (n = 1)
Included
New studies included in review
(n = 3)
Reports of new included studies
(n = 3)
Total studies included in review
(n = 11)
Reports of total included studies****
(n = 12)

**Depression**

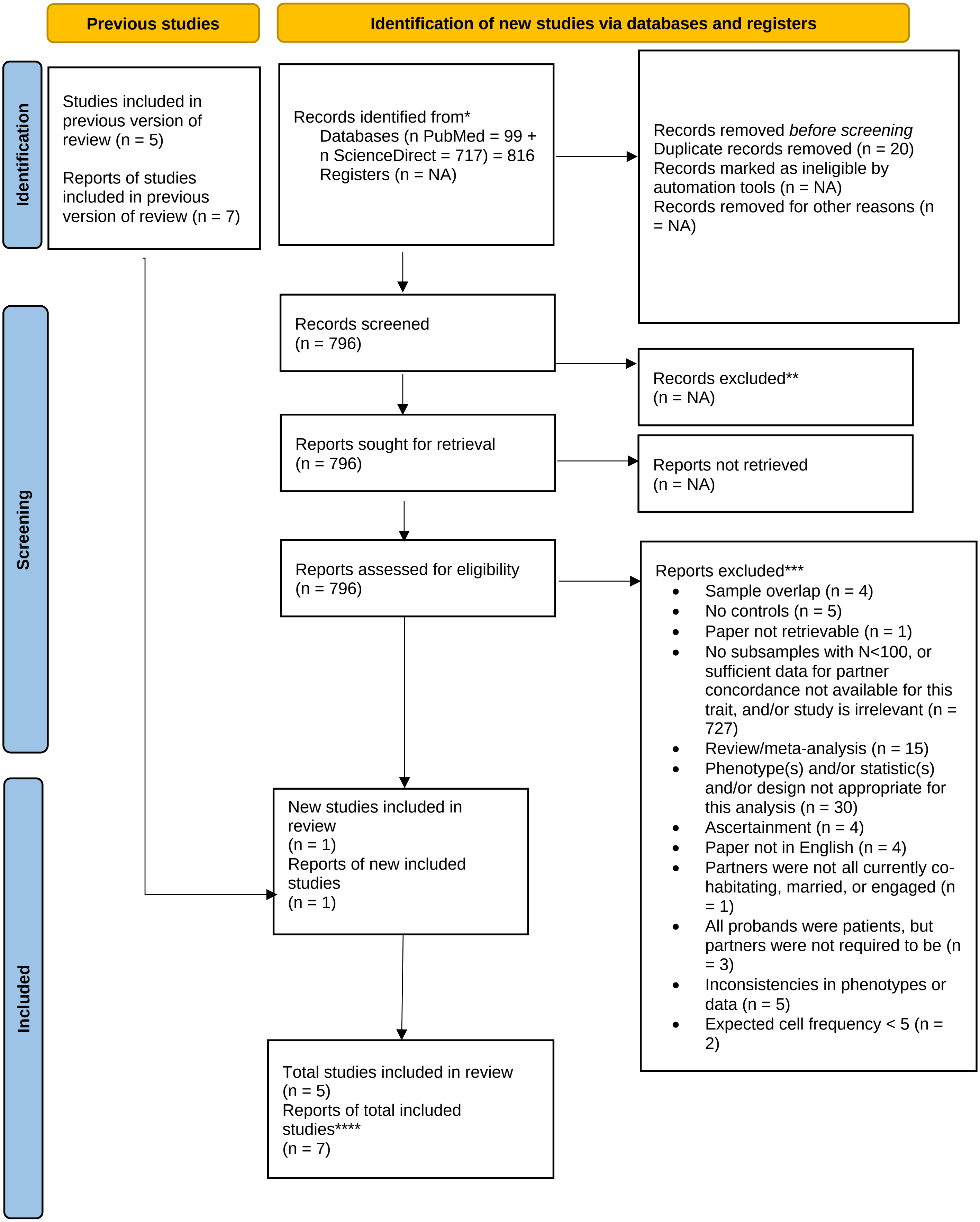
Previous studies
Identification of new studies via databases and registers
Identification
Studies included in previous version of review (n = 5)
Reports of studies included in previous version of review (n = 7)
Records identified from*
Databases (n PubMed = 99 + n ScienceDirect = 717) = 816
Registers (n = NA)
Records removed *before screening*
Duplicate records removed (n = 20)
Records marked as ineligible by automation tools (n = NA)
Records removed for other reasons (n = NA)
Screening
Records screened
(n = 796)
Records excluded**
(n = NA)
Reports sought for retrieval
(n = 796)
Reports not retrieved
(n = NA)
Reports assessed for eligibility
(n = 796)
Reports excluded***
Sample overlap (n = 4)
No controls (n = 5)
Paper not retrievable (n = 1)
No subsamples with N<100, or sufficient data for partner concordance not available for this trait, and/or study is irrelevant (n = 727)
Review/meta-analysis (n = 15)
Phenotype(s) and/or statistic(s) and/or design not appropriate for this analysis (n = 30)
Ascertainment (n = 4)
Paper not in English (n = 4)
Partners were not all currently co-habitating, married, or engaged (n = 1)
All probands were patients, but partners were not required to be (n = 3)
Inconsistencies in phenotypes or data (n = 5)
Expected cell frequency < 5 (n = 2)
Included
New studies included in review
(n = 1)
Reports of new included studies
(n = 1)
Total studies included in review
(n = 5)
Reports of total included studies****
(n = 7)

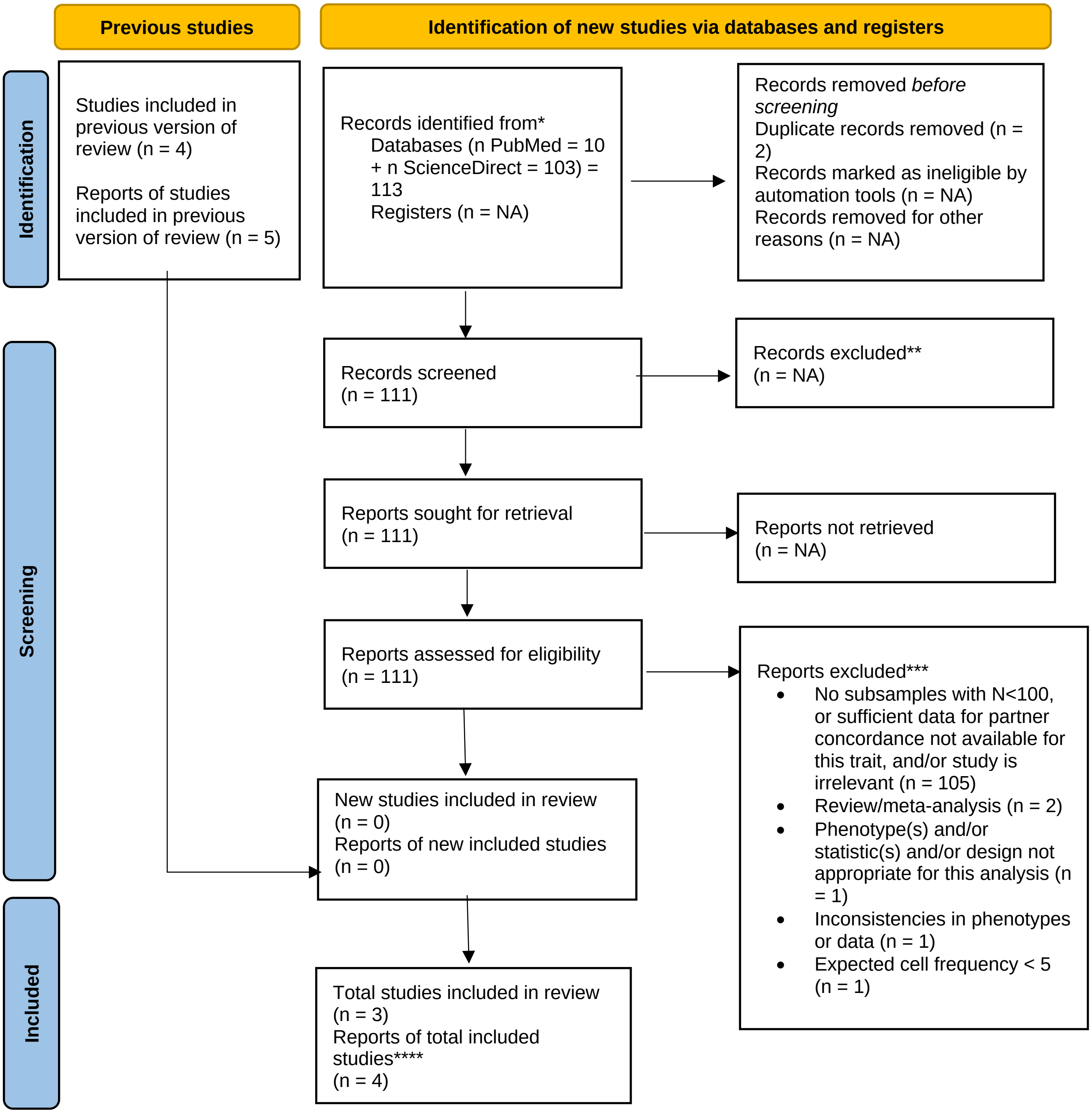


*Consider, if feasible to do so, reporting the number of records identified from each database or register searched (rather than the total number across all databases/registers).

**If automation tools were used, indicate how many records were excluded by a human and how many were excluded by automation tools.

***Total may exceed number of reports assessed for eligibility minus new studies included because some reports were excluded for more than one reason.

****Total may differ from number of reports in previous version plus number of new reports if any reports from the original version were since excluded

The PRISMA diagrams depict the number of records each search engine returned for each phenotype and the number of reports excluded for various reasons.

**Supplementary Note**

*The following lists the traits we meta-analyzed and citations corresponding to each study that was meta-analyzed for that trait, followed by descriptions of the more specific construct(s) that we measured for that trait. For the specific measure(s) that each study used for less standardized traits, see Supplementary Tables 1 and 2:*

**Educational attainment**[1–42]: Defined on an ordinal scale (e.g. range of years/schooling level completed), or by number of years of schooling. All ordinal measures of educational attainment with fewer than four points were excluded.

**Intelligence quotient score**[25,43–54]: Any measure of *general* intelligence quotient.

**Political values**[7,8,44,55–62]: We considered continuous or ordinal measures of attitudes toward political issues instead of specific party affiliation/support.

**Religiosity**[7,44,55,59,61,63,64]: We considered continuous measures of religious ideals/practices, rather than specific religious affiliation or only attendance at a place of worship.

**Problematic alcohol use**[65–68]: This was measured as a dichotomous trait encompassing alcohol abuse, alcoholism, alcohol dependence, and problem drinking.

**Drinking quantity**[9,28,69–75]: Amount of alcohol (e.g. milliliters, grams, ounces, or kilocalories) consumed per day or week**.**

**Smoking cessation**[76–79]: This was a dichotomous measure of whether subjects were former or current smokers among couples in which both partners had a history of smoking. Because the phenotype was only examined in couples in which both partners had ever smoked, whenever “never smoked” was included as an option for participants, this cell was excluded from the contingency table such that for this measure, the odds ratio reflected the odds of a partner being a former smoker if their partner was a former smoker divided by the odds of a partner being a former smoker if their partner was a current smoker.

**Smoking initiation**[9,69–71,74,76–83]: This was a dichotomous measure of whether participants had ever smoked tobacco, had ever smoked regularly, or had ever smoked 100 or more cigarettes, depending on the study. When options for participants included "former," "current," and "never," cells from those who responded "former" and "current" were combined in the contingency table such that for this measure, the odds ratio reflected the odds of a partner having ever smoked (formerly *or* currently) if their partner ever smoked (formerly *or* currently) divided by the odds of a partner having ever smoked if their partner never smoked.

**Smoking quantity**[29,84–88]: Number of cigarettes (or equivalent in tobacco) consumed per day (not restricted to those with smoking histories).

**Smoking status**[6,69,76–78,80,82,83,86,88–98]: This was a dichotomous measure of whether participants currently smoked tobacco. When options for participants included "former," "current," and "never," cells from those who responded "former" and "never" were combined in the contingency table such that for this measure, the odds ratio reflected the odds of a partner being a current smoker if their partner was a current smoker divided by the odds of a partner being a current smoker if their partner did not currently smoke (never smoked *or* formerly smoked).

**Substance use disorder**[99–101]: This was measured as a dichotomous trait also encompassing substance dependence and substance abuse.

**Agreeableness**[12,44,56,102–110]: A "Big Five" personality trait characterized by an individual's level of kindness, sympathy, cooperativeness, warmth, politeness, etc.[111] Scores for this trait were typically based on the degree to which participants agreed with statements pertaining to agreeableness or where they identified themselves on a set of spectra of bipolar adjectives pertaining to agreeableness. Ratings were typically given on a 5-point or 7-point scale (see Supplementary Table 1 for the measure used in each study).

**Conscientiousness**[12,44,56,102–110]: A “Big Five” personality trait characterized by an individual’s level of efficiency, organization, carefulness, tidiness, practicality, etc.[111] Scores for this trait were typically based on the degree to which participants agreed with statements pertaining to conscientiousness or where they identified themselves on a set of spectra of bipolar adjectives pertaining to conscientiousness. Ratings were typically given on a 5-point or 7-point scale (see Supplementary Table 1 for the measure used in each study).

**Extraversion**[7,8,12,43–45,55,56,62,63,69,85,102–110,112–120]: A “Big Five” personality trait characterized by an individual’s level of energy, talkativeness, outgoingness, etc.[111] Scores for this trait were typically based on the degree to which participants agreed with statements pertaining to extraversion or where they identified themselves on a set of spectra of bipolar adjectives pertaining to extraversion. Ratings were typically given on a 5-point or 7-point scale (see Supplementary Table 1 for the measure used in each study).

**Neuroticism**[7,12,44,45,56,62,63,69,85,102–110,112,113,115–124] (otherwise characterized by its diametric opposite, emotional stability[125]): A “Big Five” personality trait characterized by an individual’s level of enviousness, jealousy, (in)security, anxiety, moodiness, emotionality, etc.[111] Scores for this trait were typically based on the degree to which participants agreed with statements pertaining to neuroticism or where they identified themselves on a set of spectra of bipolar adjectives pertaining to neuroticism. Ratings were typically given on a 5-point or 7-point scale (see Supplementary Table 1 for the measure used in each study).

**Openness**[12,44,56,102,103,105,106,108,110,123,126] (otherwise characterized as Intellect or Openness to experiencer): A “Big Five” personality trait characterized by an individual’s level of creativeness, intelligence, imagination, etc.[111] Scores for this trait were typically based on the degree to which participants agreed with statements pertaining to openness or where they identified themselves on a set of spectra of bipolar adjectives pertaining to openness. Ratings were typically given on a 5-point or 7-point scale (see Supplementary Table 1 for the measure used in each study).

**Body mass index**[1,8,9,36,39,41,70,71,76,80,83,93,120,127–153]: Kilograms/meters$^2$ or any measure of weight/height$^2$.

**Height**[4,8,36,41,49,69,75,120,129,130,141,142,148–188]: We re-analyzed 68 samples from 53 studies from a single meta-analysis of partner correlations for standing height[189] (see main text).

**Waist-to-Hip ratio**[70,128,132,140,147,190]: Waist circumference/hip circumference**.**

**Depression**[65,66,99,191,192]: This was measured as a dichotomous trait. In addition to a classic diagnosis of major depression, we considered studies that presented partner concordance based on whether participants lay below or above a cut-off score indicating a depressive disorder/episode or a probable depressive disorder/episode. However, we excluded studies that only presented partner concordance in the form of correlations for symptom count.

**Diabetes**[77,83,99,147,191,193–197]: This was measured as a dichotomous trait. We only included studies that examined prevalent diabetes, excluding studies that only measured participants with incident diabetes (individuals who developed diabetes over the course of the study period). One study[194] reported concordance for type 2 diabetes, but we included it because 90-95% of people with diabetes have type 2[198].

**Generalized anxiety**[65,66,199]: This was measured as a dichotomous trait. In addition to a classic diagnosis of generalized anxiety disorder, we considered studies that presented partner concordance based on whether participants lay below or above a cut-off score, though we excluded studies that only presented partner concordance in the form of correlation for symptom count. We also excluded analyses measuring concordance for *any* anxiety disorder.

#### References

1. Retnakaran, R. *et al.* Spousal concordance of cardiovascular risk factors in newly married couples in china. *JAMA Netw. Open* **4**, e2140578 (2021).
2. Blackaby, D. H., Carlin, P. S. & Murphy, P. D. A change in the earnings penalty for British men with working wives: Evidence from the 1980's and 1990's. *Labour Econ.* **14**, 119–134 (2007).
3. Neidhöfer, G., Serrano, J. & Gasparini, L. Educational inequality and intergenerational mobility in Latin America: A new database. *J. Dev. Econ.* **134**, 329–349 (2018).
4. Dalmia, S. & Lawrence, P. G. An empirical analysis of assortative mating in India and the US. *Int. Adv. Econ. Res.* **7**, 443–458 (2001).
5. Kardum, I., Hudek-Knezevic, J. & Mehic, N. Similarity indices of the Dark Triad traits based on self and partner-reports: Evidence from variable-centered and couple-centered approaches. *Personal. Individ. Differ.* **193**, 111626 (2022).
6. Torvik, F. A., Gustavson, K., Røysamb, E. & Tambs, K. Health, health behaviors, and health dissimilarities predict divorce: results from the HUNT study. *BMC Psychol.* **3**, 13 (2015).
7. Feng, D. & Baker, L. Spouse similarity in attitudes, personality, and psychological well-being. *Behav. Genet.* **24**, 357–364 (1994).
8. Zietsch, B. P., Verweij, K. J., Heath, A. C. & Martin, N. G. Variation in human mate choice: simultaneously investigating heritability, parental influence, sexual imprinting, and assortative mating. *Am. Nat.* **177**, 605–616 (2011).
9. Dufouil, C. & Alpérovitch, A. Couple similarities for cognitive functions and psychological health. *J. Clin. Epidemiol.* **53**, 589–593 (2000).

10. Luo, S. & Klohnen, E. C. Assortative Mating and Marital Quality in Newlyweds: A Couple-Centered Approach. *J. Pers. Soc. Psychol.* **88**, 304–326 (2005).

11. Watkins, M. P. & Meredith, W. Spouse similarity in newlyweds with respect to specific cognitive abilities, socioeconomic status, and education. *Behav. Genet.* **11**, 1–21 (1981).

12. Botwin, M. D., Buss, D. M. & Shackelford, T. K. Personality and mate preferences: five factors in mate selection and marital satisfaction. *J. Pers.* **65**, 107–136 (1997).

13. Hugh-Jones, D., Verweij, K. J. H., St. Pourcain, B. & Abdellaoui, A. Assortative mating on educational attainment leads to genetic spousal resemblance for polygenic scores. *Intelligence* **59**, 103–108 (2016).

14. Gayle, G.-L., Golan, L. & Soytas, M. A. What is the source of the intergenerational correlation in earnings? *J. Monet. Econ.* **129**, 24–45 (2022).

15. Barban, N., De Cao, E., Oreffice, S. & Quintana-Domeque, C. The effect of education on spousal education: A genetic approach. *Labour Econ.* **71**, 102023 (2021).

16. Colagrossi, M., d'Hombres, B. & Schnepf, S. V. Like (grand)parent, like child? Multigenerational mobility across the EU. *Eur. Econ. Rev.* **130**, 103600 (2020).

17. Bingley, P., Cappellari, L. & Tatsiramos, K. Parental assortative mating and the intergenerational transmission of human capital. *Eur. Assoc. Labour Econ. World Conf. EALESOLEAASLE Berl. Ger. 25 – 27 June 2020* **77**, 102047 (2022).

18. Cave, S. N., Wright, M. & von Stumm, S. Change and stability in the association of parents' education with children's intelligence. *Intelligence* **90**, 101597 (2022).

19. Carmelli, D., Swan, G. E., Hunt, S. C. & Williams, R. R. Cross-spouse correlates of blood pressure in hypertension-prone families in Utah. *J. Psychosom. Res.* **33**, 75–84 (1989).

20. Miller, R. N. Educational assortative mating and time use in the home. *Soc. Sci. Res.* **90**, 102440 (2020).

21. Mishra, V. & Smyth, R. Economic returns to schooling for China's Korean minority. *J. Asian Econ.* **24**, 89–102 (2013).

22. Anderson, G. & Leo, T. W. An empirical examination of matching theories: The one child policy, partner choice and matching intensity in urban China. *Law Finance* **41**, 468–489 (2013).

23. Liu, J.-T., Hammitt, J. K. & Jeng Lin, C. Family background and returns to schooling in Taiwan. *Econ. Educ. Rev.* **19**, 113–125 (2000).

24. Hu, A. & Qian, Z. Does higher education expansion promote educational homogamy? Evidence from married couples of the post-80s generation in Shanghai, China. *Soc. Sci. Res.* **60**, 148–162 (2016).

25. Johnson, W., Deary, I. J. & Iacono, W. G. Genetic and environmental transactions underlying educational attainment. *Intelligence* **37**, 466–478 (2009).

26. Xing, L., Campbell, C., Li, X., Noellert, M. & Lee, J. Education, class and assortative marriage in rural Shanxi, China in the mid-twentieth century. *Res. Soc. Stratif. Mobil.* **66**, 100460 (2020).

27. Peek, M. K., Stimpson, J. P., Townsend, A. L. & Markides, K. S. Well-being in older Mexican American spouses. *The Gerontologist* **46**, 258–265 (2006).

28. Cronkite, R. C. & Moos, R. H. The role of predisposing and moderating factors in the stress-illness relationship. *J. Health Soc. Behav.* 372–393 (1984).

29. Alford, J. R., Hatemi, P. K., Hibbing, J. R., Martin, N. G. & Eaves, L. J. The politics of mate choice. *J. Polit.* **73**, 362–379 (2011).

30. Heath, A. C. *et al.* No decline in assortative mating for educational level. *Behav. Genet.* **15**, 349–369 (1985).

31. Lykken, D. T. & Tellegen, A. Is human mating adventitious or the result of lawful choice? A twin study of mate selection. *J. Pers. Soc. Psychol.* **65**, 56 (1993).

32. Kye, B. & Mare, R. D. Intergenerational effects of shifts in women's educational distribution in South Korea: Transmission, differential fertility, and assortative mating. *Soc. Sci. Res.* **41**, 1495–1514 (2012).

33. Ingoglia, S., Liga, F., Coco, A. L. & Inguglia, C. Informant discrepancies in perceived parental psychological control, adolescent autonomy, and relatedness psychological needs. *J. Appl. Dev. Psychol.* **77**, 101333 (2021).

34. Torvik, F. A. *et al.* Mechanisms linking parental educational attainment with child ADHD, depression, and academic problems: a study of extended families in The Norwegian Mother, Father and Child Cohort Study. *J. Child Psychol. Psychiatry* **61**, 1009–1018 (2020).

35. Shikishima, C. *et al.* A simple syllogism-solving test: Empirical findings and implications for g research. *Intelligence* **39**, 89–99 (2011).

36. Ponzo, M. & Scoppa, V. Trading height for education in the marriage market. *Am. J. Hum. Biol.* **27**, 164–174 (2015).

37. Procidano, M. E. & Rogler, L. H. Homogamous assortative mating among Puerto Rican families: Intergenerational processes and the migration experience. *Behav. Genet.* **19**, 343–354 (1989).

38. Chen, H., Luo, S., Yue, G., Xu, D. & Zhaoyang, R. Do birds of a feather flock together in China? *Pers. Relatsh.* **16**, 167–186 (2009).

39. George, D. *et al.* Couple similarity on stimulus characteristics and marital satisfaction. *Personal. Individ. Differ.* **86**, 126–131 (2015).

40. Jacobs, J. A. & Furstenberg Jr, F. F. Changing places: conjugal careers and women's marital mobility. *Soc. Forces* **64**, 714–732 (1986).

41. Hur, Y.-M. Assortative mating for personality traits, educational level, religious affiliation, height, weight, and body mass index in parents of a Korean twin sample. *Twin Res.* **6**, 467–470 (2003).

42. Verbakel, E. & Kalmijn, M. Assortative Mating Among Dutch Married and Cohabiting Same-Sex and Different-Sex Couples: Homogamy in Same- and Different-Sex Couples. *J. Marriage Fam.* **76**, 1–12 (2014).

43. Phillips, K., Fulker, D. W., Carey, G. & Nagoshi, C. T. Direct marital assortment for cognitive and personality variables. *Behav. Genet.* **18**, 347–356 (1988).

44. Watson, D. *et al.* Match makers and deal breakers: analyses of assortative mating in newlywed couples. *J. Pers.* **72**, 1029–1068 (2004).

45. Mascie-Taylor, C. G. N. Spouse similarity for IQ and personality and convergence. *Behav. Genet.* **19**, 223–227 (1989).

46. Tambs, K., Sundet, J. M. & Berg, K. Correlations between identical twins and their spouses suggest social homogamy for intelligence in Norway. *Personal. Individ. Differ.* **14**, 279–281 (1993).

47. Cattell, R. B. & Willson, J. L. Contributions concerning mental inheritance: I. of intelligence. *Br. J. Educ. Psychol.* **8**, 129–149 (1938).

48. Johnson, W. *et al.* Genetic and environmental influences on the Verbal-Perceptual-Image Rotation (VPR) model of the structure of mental abilities in the Minnesota study of twins reared apart. *Intelligence* **35**, 542–562 (2007).

49. Keller, M. C. *et al.* The genetic correlation between height and IQ: shared genes or assortative mating? *PLoS Genet.* **9**, e1003451 (2013).

50. Nagoshi, C. T. & Johnson, R. C. The ubiquity of g. *Personal. Individ. Differ.* **7**, 201–207 (1986).

51. Jester, J. M. *et al.* Intergenerational transmission of neuropsychological executive functioning. *Brain Cogn.* **70**, 145–153 (2009).

52. Vinkhuyzen, A. A. E., van der Sluis, S., Maes, H. H. M. & Posthuma, D. Reconsidering the Heritability of Intelligence in Adulthood: Taking Assortative Mating and Cultural Transmission into Account. *Behav. Genet.* **42**, 187–198 (2012).

53. Loehlin, J. C., Horn, J. M. & Willerman, L. Modeling IQ change: evidence from the Texas Adoption Project. *Child Dev.* 993–1004 (1989).

54. Zonderman, A. B., Vandenberg, S. G., Spuhler, K. P. & Fain, P. R. Assortative marriage for cognitive abilities. *Behav. Genet.* **7**, 261–271 (1977).

55. Eaves, L. *et al.* Comparing the biological and cultural inheritance of personality and social attitudes in the Virginia 30,000 study of twins and their relatives. *Twin Res.* **2**, 62–80 (1999).

56. Leikas, S., Ilmarinen, V.-J., Verkasalo, M., Vartiainen, H.-L. & Lönnqvist, J.-E. Relationship satisfaction and similarity of personality traits, personal values, and attitudes. *Personal. Individ. Differ.* **123**, 191–198 (2018).

57. Martin, N. G. *et al.* Transmission of social attitudes. *Proc. Natl. Acad. Sci.* **83**, 4364–4368 (1986).

58. Feather, N. T. Family resemblances in conservatism: Are daughters more similar to parents than sons are? *J. Pers.* **46**, 260–278 (1978).

59. Rozin, P. Family resemblance in food and other domains: The family paradox and the role of parental congruence. *Appetite* **16**, 93–102 (1991).

60. Bell, E., Kandler, C. & Riemann, R. Genetic and environmental influences on sociopolitical attitudes: addressing some gaps in the new paradigm. *Polit. Life Sci.* **37**, 236–249 (2018).

61. Abrahamson, A. C., Baker, L. A. & Caspi, A. Rebellious teens? Genetic and environmental influences on the social attitudes of adolescents. *J. Pers. Soc. Psychol.* **83**, 1392–1408 (2002).

62. Eysenck, H. J. & Wakefield Jr, J. A. Psychological factors as predictors of marital satisfaction. *Adv. Behav. Res. Ther.* **3**, 151–192 (1981).

63. Beer, J. M., Arnold, R. D. & Loehlin, J. C. Genetic and environmental influences on MMPI factor scales: joint model fitting to twin and adoption data. *J. Pers. Soc. Psychol.* **74**, 818 (1998).

64. Koenig, L. B., McGue, M. & Iacono, W. G. Rearing environmental influences on religiousness: An investigation of adolescent adoptees. *Personal. Individ. Differ.* **47**, 652–656 (2009).

65. Galbaud Du Fort, G., Bland, R. C., Newman, S. C. & Boothroyd, L. J. Spouse similarity for lifetime psychiatric history in the general population. *Psychol. Med.* **28**, 789–802 (1998).

66. Maes, H. H. *et al.* Assortative mating for major psychiatric diagnoses in two population-based samples. *Psychol. Med.* **28**, 1389–1401 (1998).

67. Ostermann, J., Sloan, F. A. & Taylor, D. H. Heavy alcohol use and marital dissolution in the USA. *Soc. Sci. Med.* **61**, 2304–2316 (2005).

68. Grant, J. D. *et al.* Spousal concordance for alcohol dependence: evidence for assortative mating or spousal interaction effects? *Alcohol. Clin. Exp. Res.* **31**, 717–728 (2007).

69. Price, R. A. & Vandenberg, S. G. Spouse similarity in American and Swedish couples. *Behav. Genet.* **10**, 59–71 (1980).

70. Di Castelnuovo, A. *et al.* Cardiovascular risk factors and global risk of fatal cardiovascular disease are positively correlated between partners of 802 married couples from different European countries. *Thromb. Haemost.* **98**, 648–655 (2007).

71. Al Rashid, K. *et al.* Spousal associations of serum metabolomic profiles by nuclear magnetic resonance spectroscopy. *Sci. Rep.* **11**, 21587 (2021).

72. Pérusse, L., Leblanc, C. & Bouchard, C. Familial resemblance in lifestyle components: results from the Canada Fitness Survey. *Can. J. Public Health Rev. Can. Sante Publique* **79**, 201–205 (1988).

73. Leonard, K. E. & Das Eiden, R. Husband’s and wife’s drinking: unilateral or bilateral influences among newlyweds in a general population sample. *J. Stud. Alcohol. Suppl.* 130–138 (1999).

74. Clarke, T.-K. *et al.* Genetic and shared couple environmental contributions to smoking and alcohol use in the UK population. *Mol. Psychiatry* **26**, 4344–4354 (2021).

75. Garn, S. M., Cole, P. E. & Bailey, S. M. Living together as a factor in family-line resemblances. *Hum. Biol.* **51**, 565–587 (1979).

76. Pyke, S. D., Wood, D. A., Kinmonth, A. L. & Thompson, S. G. Change in coronary risk and coronary risk factor levels in couples following lifestyle intervention. The British Family Heart Study. *Arch. Fam. Med.* **6**, 354–360 (1997).

77. Jurj, A. L. *et al.* Spousal correlations for lifestyle factors and selected diseases in Chinese couples. *Ann. Epidemiol.* **16**, 285–291 (2006).

78. Venters, M. H., Jacobs Jr, D. R., Luepker, R. V., Maimaw, L. A. & Gillum, R. F. Spouse concordance of smoking patterns: the Minnesota Heart Survey. *Am. J. Epidemiol.* **120**, 608–616 (1984).

79. Cobb, L. K. *et al.* The association of spousal smoking status with the ability to quit smoking: the Atherosclerosis Risk in Communities Study. *Am. J. Epidemiol.* **179**, 1182–1187 (2014).

80. Stimpson, J. P., Masel, M. C., Rudkin, L. & Peek, M. K. Shared health behaviors among older Mexican American spouses. *Am. J. Health Behav.* **30**, 495–502 (2006).

81. Maes, H. H. *et al.* Cross-cultural comparison of genetic and cultural transmission of smoking initiation using an extended twin kinship model. *Twin Res. Hum. Genet.* **21**, 179–190 (2018).

82. Boomsma, D. I., Koopmans, J. R., Van Doornen, L. J. & Orlebeke, J. F. Genetic and social influences on starting to smoke: a study of Dutch adolescent twins and their parents. *Addiction* **89**, 219–226 (1994).

83. Nakaya, N. *et al.* Spousal similarities in cardiometabolic risk factors: A cross-sectional comparison between Dutch and Japanese data from two large biobank studies. *Atherosclerosis* **334**, 85–92 (2021).

84. Wilcox, M. A., Newton, C. S. & Johnson, I. R. Paternal influences on birthweight. *Acta Obstet. Gynecol. Scand.* **74**, 15–18 (1995).

85. Gleiberman, L., Harburg, E., DiFranceisco, W. & Schork, A. Familial transmission of alcohol use: v. drinking patterns among spouses, tecumseh, michigan. *Behav. Genet.* **22**, 63–79 (1992).

86. Sutton, G. C. Assortative marriage for smoking habits. *Ann. Hum. Biol.* **7**, 449–456 (1980).

87. Cotch, M. F., Beaty, T. H. & Cohen, B. H. Path analysis of familial resemblance of pulmonary function and cigarette smoking. *Am. Rev. Respir. Dis.* **142**, 1337–1343 (1990).

88. Kalmijn, M. Intergenerational transmission of health behaviors in a changing demographic context: The case of smoking and alcohol consumption. *Soc. Sci. Med.* **296**, 114736 (2022).

89. Bloch, K. V., Klein, C. H., de Souza e Silva, N. A., Nogueira, A. da R. & Salis, L. H. A. Socioeconomic aspects of spousal concordance for hypertension, obesity, and smoking in a community of Rio de Janeiro, Brazil. *Arq. Bras. Cardiol.* **80**, 179–186, 171–178 (2003).

90. Clark, A. E. & Etilé, F. Don't give up on me baby: Spousal correlation in smoking behaviour. *J. Health Econ.* **25**, 958–978 (2006).

91. Ogden, M. W., Morgan, W. T., Heavner, D. L., Davis, R. A. & Steichen, T. J. National incidence of smoking and misclassification among the U.S. married female population. *J. Clin. Epidemiol.* **50**, 253–263 (1997).

92. Espinosa, J. & Evans, W. N. Heightened mortality after the death of a spouse: Marriage protection or marriage selection? *J. Health Econ.* **27**, 1326–1342 (2008).

93. Wilson, S. E. The health capital of families: an investigation of the inter-spousal correlation in health status. *Soc. Sci. Med.* **55**, 1157–1172 (2002).

94. Jackson, S. E., Steptoe, A. & Wardle, J. The influence of partner's behavior on health behavior change: the English Longitudinal Study of Ageing. *JAMA Intern. Med.* **175**, 385–392 (2015).

95. Jeong, S. & Cho, S.-I. Concordance in the health behaviors of couples by age: a cross-sectional study. *J. Prev. Med. Pub. Health* **51**, 6 (2018).

96. Homish, G. G. & Leonard, K. E. Spousal influence on smoking behaviors in a US community sample of newly married couples. *Soc. Sci. Med.* **61**, 2557–2567 (2005).

97. Pai, C.-W., Godboldo-Brooks, A. & Edington, D. W. Spousal concordance for overall health risk status and preventive service compliance. *Ann. Epidemiol.* **20**, 539–546 (2010).

98. Machado, M. P. A. *et al.* Alcohol and tobacco consumption concordance and its correlates in older couples in Latin America. *Geriatr. Gerontol. Int.* **17**, 1849–1857 (2017).

99. Nordsletten, A. E. *et al.* Patterns of nonrandom mating within and across 11 major psychiatric disorders. *JAMA Psychiatry* **73**, 354–361 (2016).

100. Kirillova, G. P., Vanyukov, M. M., Kirisci, L. & Reynolds, M. Physical maturation, peer environment, and the ontogenesis of substance use disorders. *Psychiatry Res.* **158**, 43–53 (2008).

101. Sakai, J. T. *et al.* Mate similarity for substance dependence and antisocial personality disorder symptoms among parents of patients and controls. *Drug Alcohol Depend.* **75**, 165–175 (2004).

102. McCrae, R. R. *et al.* Personality trait similarity between spouses in four cultures. *J. Pers.* **76**, 1137–1164 (2008).

103. Rammstedt, B. & Schupp, J. Only the congruent survive–personality similarities in couples. *Personal. Individid. Differ.* **45**, 533–535 (2008).

104. Chopik, W. J. & Lucas, R. E. Actor, partner, and similarity effects of personality on global and experienced well-being. *J. Res. Personal.* **78**, 249–261 (2019).

105. Vandermeer, M. R. J., Kotelnikova, Y., Simms, L. J. & Hayden, E. P. Spousal Agreement on Partner Personality Ratings is Moderated by Relationship Satisfaction. *J. Res. Personal.* **76**, 22–31 (2018).

106. Watson, D., Beer, A. & McDade-Montez, E. The role of active assortment in spousal similarity. *J. Pers.* **82**, 116–129 (2014).

107. Barelds, D. P. Self and partner personality in intimate relationships. *Eur. J. Personal. Publ. Eur. Assoc. Personal. Psychol.* **19**, 501–518 (2005).

108. van Scheppingen, M. A., Chopik, W. J., Bleidorn, W. & Denissen, J. J. A. Longitudinal Actor, Partner and Similarity Effects of Personality on Well-Being. *J. Pers. Soc. Psychol.* **117**, e51–e70 (2019).

109. Dijkstra, P. & Barelds, D. P. H. Self and partner personality and responses to relationship threats. *J. Res. Personal.* **42**, 1500–1511 (2008).

110. Mosca, I. & McCrory, C. Personality and wealth accumulation among older couples: Do dispositional characteristics pay dividends? *J. Econ. Psychol.* **56**, 1–19 (2016).

111. Thompson, E. R. Development and validation of an international english big-five mini-markers. *Personal. Individ. Differ.* **45**, 542–548 (2008).

112. Dubuis-Stadelmann, E., Fenton, B. T., Ferrero, F. & Preisig, M. Spouse similarity for temperament, personality and psychiatric symptomatology. *Personal. Individ. Differ.* **30**, 1095–1112 (2001).

113. Tambs, K., Sundet, J. M., Eaves, L., Solaas, M. H. & Berg, K. Pedigree analysis of Eysenck Personality Questionnaire (EPQ) scores in monozygotic (MZ) twin families. *Behav. Genet.* **21**, 369–382 (1991).

114. Guttman, R. & Zohar, A. Spouse similarities in personality items: Changes over years of marriage and implications for mate selection. *Behav. Genet.* **17**, 179–189 (1987).

115. Eysenck, H. J. Personality, premarital sexual permissiveness, and assortative mating. *J. Sex Res.* **10**, 47–51 (1974).

116. Patterson, C. H. The relationship of Bernreuter scores to parent behavior, child behavior, urban-rural residence, and other background factors in 100 normal adult parents. *J. Soc. Psychol.* **24**, 3–49 (1946).

117. Terman, L. M. & Buttenwieser, P. Personality factors in marital compatibility: II. *J. Soc. Psychol.* **6**, 267–289 (1935).

118. Rushton, J. P. & Bons, T. A. Mate choice and friendship in twins: evidence for genetic similarity. *Psychol. Sci.* **16**, 555–559 (2005).

119. Farley, F. H. & Davis, S. A. Arousal, Personality, and Assortative Mating in Marriage. *J. Sex Marital Ther.* **3**, 122–127 (1977).

120. Sutton, G. C. Do men grow to resemble their wives, or vice versa? *J. Biosoc. Sci.* **25**, 25–29 (1993).

121. Abdellaoui, A. *et al.* Associations between loneliness and personality are mostly driven by a genetic association with Neuroticism. *J. Pers.* **87**, 386–397 (2019).

122. Hoffeditz, E. L. Family resemblances in personality traits. *J. Soc. Psychol.* **5**, 214–227 (1934).

123. Donnellan, M. B., Conger, R. D. & Bryant, C. M. The Big Five and enduring marriages. *J. Res. Personal.* **38**, 481–504 (2004).

124. Lake, R. I., Eaves, L. J., Maes, H. H., Heath, A. C. & Martin, N. G. Further evidence against the environmental transmission of individual differences in neuroticism from a collaborative study of 45,850 twins and relatives on two continents. *Behav. Genet.* **30**, 223–233 (2000).

125. Ormel, J., Riese, H. & Rosmalen, J. G. M. Interpreting neuroticism scores across the adult life course: immutable or experience-dependent set points of negative affect? *Clin. Psychol. Rev.* **32**, 71–79 (2012).

126. Chopik, W. J. & Johnson, D. J. Modeling dating decisions in a mock swiping paradigm: An examination of participant and target characteristics. *J. Res. Personal.* **92**, 104076 (2021).

127. Sebro, R., Peloso, G. M., Dupuis, J. & Risch, N. J. Structured mating: patterns and implications. *PLoS Genet.* **13**, e1006655 (2017).

128. Kim, H. C. *et al.* Spousal concordance of metabolic syndrome in 3141 Korean couples: a nationwide survey. *Ann. Epidemiol.* **16**, 292–298 (2006).

129. Pennock-Román, M. Assortative marriage for physical characteristics in newlyweds. *Am. J. Phys. Anthropol.* **64**, 185–190 (1984).

130. Staessen, J. *et al.* Familial aggregation of blood pressure, anthropometric characteristics and urinary excretion of sodium and potassium—a population study in two Belgian towns. *J. Chronic Dis.* **38**, 397–407 (1985).

131. Katzmarzyk, P. T., Hebebrand, J. & Bouchard, C. Spousal resemblance in the Canadian population: implications for the obesity epidemic. *Int. J. Obes.* **26**, 241–246 (2002).

132. Wu, D.-M. *et al.* Familial resemblance of adiposity-related parameters: results from a health check-up population in Taiwan. *Eur. J. Epidemiol.* **18**, 221–226 (2003).

133. Zahra, J., Jago, R. & Sebire, S. J. Associations between parenting partners' objectively-assessed physical activity and Body Mass Index: A cross-sectional study. *Prev. Med. Rep.* **2**, 473–477 (2015).

134. Longini Jr, I. M., Higgins, M. W., Hinton, P. C., Moll, P. P. & Keller, J. B. Genetic and environmental sources of familial aggregation of body mass in Tecumseh, Michigan. *Hum. Biol.* 733–757 (1984).

135. Tambs, K. *et al.* Genetic and environmental contributions to the variance of the body mass index in a Norwegian sample of first- and second-degree relatives. *Am. J. Hum. Biol.* **3**, 257–267 (1991).

136. Knuiman, M. W., Divitini, M. L., Bartholomew, H. C. & Welborn, T. A. Spouse correlations in cardiovascular risk factors and the effect of marriage duration. *Am. J. Epidemiol.* **143**, 48–53 (1996).

137. Jee, S. H., Suh, I., Won, S. Y. & Kim, M. Y. Familial correlation and heritability for cardiovascular risk factors. *Yonsei Med. J.* **43**, 160–164 (2002).

138. Davillas, A. & Pudney, S. Concordance of health states in couples: analysis of self-reported, nurse administered and blood-based biomarker data in the UK Understanding Society panel. *J. Health Econ.* **56**, 87–102 (2017).

139. Moll, P. P., Burns, T. L. & Lauer, R. M. The genetic and environmental sources of body mass index variability: the Muscatine Ponderosity Family Study. *Am. J. Hum. Genet.* **49**, 1243 (1991).

140. van Dongen, J., Willemsen, G., Chen, W.-M., de Geus, E. J. C. & Boomsma, D. I. Heritability of metabolic syndrome traits in a large population-based sample[S]. *J. Lipid Res.* **54**, 2914–2923 (2013).

141. Prichard, I. *et al.* Brides and young couples: partners’ weight, weight change, and perceptions of attractiveness. *J. Soc. Pers. Relatsh.* **32**, 263–278 (2015).

142. Province, M. A. & Rao, D. C. Path analysis of family resemblance with temporal trends: applications to height, weight, and Quetelet index in northeastern Brazil. *Am. J. Hum. Genet.* **37**, 178 (1985).

143. Chien, K. L. *et al.* Familial aggregation of metabolic syndrome among the Chinese: Report from the Chin-Shan community family study. *Diabetes Res. Clin. Pract.* **76**, 418–424 (2007).

144. Knight, B. *et al.* Evidence of genetic regulation of fetal longitudinal growth. *Early Hum. Dev.* **81**, 823–831 (2005).

145. Bouchard, C., Perusse, L., Leblanc, C., Tremblay, A. & Theriault, G. Inheritance of the amount and distribution of human body fat. *Int. J. Obes.* **12**, 205–215 (1988).

146. Friedlander, Y., Kark, J. D., Kaufmann, N. A., Berry, E. M. & Stein, Y. Familial aggregation of body mass index in ethnically diverse families in Jerusalem. The Jerusalem Lipid Research Clinic. *Int. J. Obes.* **12**, 237–247 (1988).

147. Silverman-Retana, O. *et al.* Spousal concordance in pathophysiological markers and risk factors for type 2 diabetes: a cross-sectional analysis of The Maastricht Study. *BMJ Open Diabetes Res. Care* **9**, e001879 (2021).

148. Sear, R. & Marlowe, F. W. How universal are human mate choices? Size does not matter when Hadza foragers are choosing a mate. *Biol. Lett.* **5**, 606–609 (2009).

149. Silventoinen, K., Kaprio, J., Lahelma, E., Viken, R. J. & Rose, R. J. Assortative mating by body height and BMI: Finnish twins and their spouses. *Am. J. Hum. Biol.* **15**, 620–627 (2003).

150. Mueller, W. H. & Malina, R. M. Differential contribution of stature phenotypes to assortative mating in parents of Philadelphia black and white school children. *Am. J. Phys. Anthropol.* **45**, 269–276 (1976).

151. Heude, B. *et al.* Anthropometric relationships between parents and children throughout childhood: the Fleurbaix–Laventie Ville Santé Study. *Int. J. Obes.* **29**, 1222–1229 (2005).

152. Sanchez-Andres, A. & Mesa, M. Assortative mating in a Spanish population: effects of social factors and cohabitation time. *J. Biosoc. Sci.* **26**, 441–50 (1994).

153. Salces, I., Rebato, E. & Susanne, C. Evidence of phenotypic and social assortative mating for anthropometric and physiological traits in couples from the Basque country (Spain). *J. Biosoc. Sci.* **36**, 235–250 (2004).

154. Ahmad, M. & Gilbert, R. I. Assortative mating for height in Pakistani arranged marriages. *J. Biosoc. Sci.* **17**, 211–214 (1985).

155. Ajala, O. *et al.* The relationship of height and body fat to gender-assortative weight gain in children. A longitudinal cohort study (EarlyBird 44). *Int. J. Pediatr. Obes.* **6**, 223–228 (2011).

156. Annest, J. L., Sing, C. F., Biron, P. & Mongeau, J. G. Familial aggregation of blood pressure and weight in adoptive families: III. analysis of the role of shared genes and shared household environment in explaining family resemblance for height, weight and selected weight/height indices. *Am. J. Epidemiol.* **117**, 492–506 (1983).

157. Burgess, E. W. & Wallin, P. Homogamy in personality characteristics. *J. Abnorm. Soc. Psychol.* **39**, 475 (1944).

158. Byard, P. J., Poosha, D. V. R. & Satyanarayana, M. Genetic and environmental determinants of height and weight in families from Andhra Pradesh, India. *Hum. Biol.* 621–633 (1985).

159. Byard, P. J., Mukherjee, B. N., Bhattacharya, S. K., Russell, J. M. & Rao, D. C. Familial aggregation of blood pressure and anthropometric variables in patrilocal households. *Am. J. Phys. Anthropol.* **79**, 305–311 (1989).

160. Dasgupta, I., Dasgupta, P. & Daschaudhuri, A. B. Familial resemblance in height and weight in an endogamous Hahisya caste population of rural West Bengal. *Am. J. Hum. Biol. Off. J. Hum. Biol. Assoc.* **9**, 7–9 (1997).

161. Eckman, R. E., Williams, R. & Nagoshi, C. Marital assortment for genetic similarity. *J. Biosoc. Sci.* **34**, 511–523 (2002).

162. Ellis, J. A. *et al.* Comprehensive multi-stage linkage analyses identify a locus for adult height on chromosome 3p in a healthy Caucasian population. *Hum. Genet.* **121**, 213–222 (2007).

163. Ginsburg, E., Livshits, G., Yakovenko, K. & Kobyliansky, E. Major gene control of human body height, weight and BMI in five ethnically different populations. *Ann. Hum. Genet.* **62**, 307–322 (1998).

164. Harrison, G. A., Gibson, J. B. & Hiorns, R. W. Assortative marriage for psychometric, personality and anthropometric variation in a group of Oxfordshire villages. *J. Biosoc. Sci.* **8**, 145–153 (1976).

165. Knuiman, M. W., Divitini, M. L. & Bartholomew, H. C. Spouse selection and environmental effects on spouse correlation in lung function measures. *Ann. Epidemiol.* **15**, 39–43 (2005).

166. Luo, Z. C., Albertsson-Wikland, K. & Karlberg, J. Target height as predicted by parental heights in a population-based study. *Pediatr. Res.* **44**, 563–571 (1998).

167. Mascie-Taylor, C. G. N. Assortative mating in a contemporary British population. *Ann. Hum. Biol.* **14**, 59–68 (1987).

168. McManus, I. C. & Mascie-Taylor, C. G. Human assortative mating for height: non-linearity and heteroscedasticity. *Hum. Biol.* **56**, 617–623 (1984).

169. Mukhopadhyay, N. *et al.* A genome-wide scan for loci affecting normal adult height in the Framingham Heart Study. *Hum. Hered.* **55**, 191–201 (2003).

170. Nagoshi, C. T. & Johnson, R. C. Between- vs. within-family analyses of the correlation of height and intelligence. *Soc. Biol.* **34**, 110–113 (1987).

171. Nance, W. E., Corey, L. A. & Eaves, L. J. A model for the analysis of mate selection in the marriages of twins application to data on stature. *Acta Genet. Medicae Gemellol. Twin Res.* **29**, 91–101 (1980).

172. Pearson, K. & Lee, A. On the laws of inheritance in man: I. Inheritance of physical characters. *Biometrika* **2**, 357–462 (1903).

173. Pieper, U. Assortative mating in the population of a German and a Cameroon city. *J. Hum. Evol.* **10**, 643–645 (1981).

174. Pomerat, C. M. Homogamy and infertility. *Hum. Biol.* **8**, 19 (1936).

175. Raychaudhuri, A., Ghosh, R., Vasulu, T. S. & Bharati, P. Heritability estimates of height and weight in Mahishya caste population. *Int. J. Hum. Genet.* **3**, 151–154 (2003).

176. Roberts, D. F., Billewicz, W. Z. & McGregor, I. Heritability of stature in a West African population. *Ann. Hum. Genet.* **42**, 15–24 (1978).

177. Seki, M., Ihara, Y. & Aoki, K. Homogamy and imprinting-like effect on mate choice preference for body height in the current Japanese population. *Ann. Hum. Biol.* **39**, 28–35 (2012).

178. Siniarska, A. Assortative mating of parents and sib-sib similarities in offspring. *Stud Hum Ecol* **5**, 95–112 (1984).

179. Smith, M. A research note on homogamy of marriage partners in selected physical characteristics. *Am. Sociol. Rev.* **11**, (1946).

180. Stulp, G., Buunk, A. P. & Pollet, T. V. Women want taller men more than men want shorter women. *Personal. Individ. Differ.* **54**, 877–883 (2013).

181. Stulp, G., Buunk, A. P., Pollet, T. V., Nettle, D. & Verhulst, S. Are human mating preferences with respect to height reflected in actual pairings? *PLoS One* **8**, e54186 (2013).

182. Stulp, G., Mills, M., Pollet, T. V. & Barrett, L. Non-linear associations between stature and mate choice characteristics for American men and their spouses. *Am. J. Hum. Biol.* **26**, 530–537 (2014).

183. Susanne, C. Heritability of anthropological characters. *Hum. Biol.* 573–580 (1977).

184. Tambs, K. *et al.* Genetic and environmental contributions to the variance of body height in a sample of first and second degree relatives. *Am. J. Phys. Anthropol.* **88**, 285–294 (1992).

185. Tenesa, A., Rawlik, K., Navarro, P. & Canela-Xandri, O. Genetic determination of height-mediated mate choice. *Genome Biol.* **16**, 1–8 (2015).

186. To, W. W. K., Cheung, W. & Kwok, J. S. Y. Paternal height and weight as determinants of birth weight in a Chinese population. *Am. J. Perinatol.* **15**, 545–548 (1998).

187. Willoughby, R. R. Somatic homogamy in man. *Hum. Biol.* **5**, 690 (1933).

188. Xu, J. *et al.* Major recessive gene(s) with considerable residual polygenic effect regulating adult height: confirmation of genomewide scan results for chromosomes 6, 9, and 12. *Am. J. Hum. Genet.* **71**, 646–650 (2002).

189. Stulp, G., Simons, M. J. P., Grasman, S. & Pollet, T. V. Assortative mating for human height: a meta-analysis. *Am. J. Hum. Biol.* **29**, e22917 (2017).

190. Donahue, R. P., Prineas, R. J., Gomez, O. & Hong, C. P. Familial resemblance of body fat distribution: the Minneapolis Children's Blood Pressure Study. *Int. J. Obes. Relat. Metab. Disord. J. Int. Assoc. Study Obes.* **16**, 161–167 (1992).

191. Hippisley-Cox, J. Married couples' risk of same disease: cross sectional study. *BMJ* **325**, 636–636 (2002).

192. McLeod, J. D. Spouse concordance for depressive disorders in a community sample. *J. Affect. Disord.* **27**, 43–52 (1993).

193. Al-Sharbatti, S. S., Abed, Y. I., Al-Heety, L. M. & Basha, S. A. Spousal concordance of diabetes mellitus among women in Ajman, United Arab Emirates. *Sultan Qaboos Univ. Med. J.* **16**, e197 (2016).

194. Sun, J. *et al.* Prevalence of diabetes and cardiometabolic disorders in spouses of diabetic individuals. *Am. J. Epidemiol.* **184**, 400–409 (2016).

195. Patel, S. A. *et al.* Chronic disease concordance within Indian households: a cross-sectional study. *PLoS Med.* **14**, e1002395 (2017).

196. Jun, S. Y., Kang, M., Kang, S. Y., Lee, J. A. & Kim, Y. S. Spousal Concordance regarding Lifestyle Factors and Chronic Diseases among Couples Visiting Primary Care Providers in Korea. *Korean J. Fam. Med.* **41**, 183–188 (2020).

197. Watanabe, T., Sugiyama, T., Takahashi, H., Noguchi, H. & Tamiya, N. Concordance of hypertension, diabetes and dyslipidaemia in married couples: cross-sectional study using nationwide survey data in Japan. *BMJ Open* **10**, e036281 (2020).

198. Tama, B. A., Rodiyatul F. S., R. F. S. & Hermansyah, H. An early detection method of type-2 diabetes mellitus in public hospital. *TELKOMNIKA Telecommun. Comput. Electron. Control* **9**, 287 (2011).

199. McLeod, J. D. Social and psychological bases of homogamy for common psychiatric disorders. *J. Marriage Fam.* 201–214 (1995).
